# Walking Alone to the North: The Origin and Historical Expansion of a Polyploid Parthenogenetic Lineage in a Weevil

**DOI:** 10.64898/2026.08.06.743411

**Authors:** Shota Murakami, Po-Wei Hsu, Takashi Sato, Isao Matoba, Shigeto Dobata

## Abstract

**Background:** Polyploid parthenogenetic organisms are distributed nonrandomly with respect to their diploid sexual relatives, and this pattern has been well documented in plants. Comparable cases are rare in animals, and the genomic origins of such lineages have been reconstructed in only a few taxa. Additional animal examples are therefore needed to distinguish general eco-evolutionary processes from taxonomic idiosyncrasies.

**Results:** We determined the sex of 2,223 individuals of the flightless weevil *Catapionus nebulosus* species group from 294 localities across Japan. The sex ratio shifted sharply toward females in northern Japan. Phylogenetic analyses using mitochondrial and genome-wide SNP data assigned all female-only populations to a single lineage. This lineage consisted of tetraploid and pentaploid individuals, as determined by flow cytometry. Parthenogenesis was confirmed by the production of fertile female offspring from virgin females reared in isolation. Admixture analyses and model choice supported hybrid ancestry of this lineage from two divergent sexual lineages. Phylogeographic analyses showed that this lineage expanded rapidly in both range and effective population size. In the sexual populations, larval density and individual heterozygosity were both lower toward the northern range margin, including south of the co-occurrence zone with the parthenogenetic lineage.

**Conclusions:** The all-female lineage of the *Catapionus nebulosus* species group is a single polyploid parthenogenetic lineage with hybrid ancestry, although the order in which these traits arose remains unresolved. This lineage expanded northward and now occurs where sexual populations are sparse or absent. This pattern is consistent with several mutually non-exclusive hypotheses about polyploidy and parthenogenesis, including mate limitation as described in Baker’s law. This weevil is a new animal example of a syndrome well documented in plants, in which polyploid parthenogens with hybrid ancestry occur at higher latitudes than their sexual relatives. It offers a valuable system for studying the eco-evolutionary processes underlying this geographic distribution.

## Background

Polyploidy, the possession of more than two complete chromosome sets, has arisen repeatedly across eukaryotes and has been a major driver of their evolutionary diversification [1]. Polyploid organisms have long been known to be distributed nonrandomly with respect to their diploid relatives, often occupying cold and climatically fluctuating environments, such as higher latitudes [2–6]. A comparable bias is observed for asexual organisms and is known as geographic parthenogenesis [7, 8]. These convergent patterns are often empirically inseparable, because polyploidy is closely associated with asexual reproduction, which often originates through hybridization [9, 10]. This co-occurrence points to shared evolutionary processes that shape the geographic distribution of polyploid and asexual genomes.

Several ecological and evolutionary theories have been proposed to explain the geographic distribution of polyploid lineages. Genome duplication increases cell size and alters physiological performance, which may offer a fitness benefit in harsher environments [11, 12]. Polyploidy enables individuals to carry more than two allelic variants per locus, especially in allopolyploids, which buffers deleterious mutations [1]. Moreover, the resulting genomic redundancy provides duplicated gene copies with opportunities for sub- and neofunctionalization over evolutionary time [1]. Even in the absence of such fitness advantages, the biased distribution can be explained by interactions with diploid relatives. Newly arisen polyploids are reproductively swamped in dense diploid populations (minority cytotype exclusion [13, 14]), favoring establishment where diploids are sparse, such as at an expanding range edge [15]. At the range edge, small population sizes also intensify genetic drift, which can fix a rare polyploid variant by chance rather than through selection [16, 17]. When polyploidy is associated with asexuality, a single uniparental founder can establish a population where mates are scarce (Baker’s law [18, 19]). The clonal genotypes can broaden or shift environmental tolerance relative to sexual progenitors [1, 20], either through a tolerant general-purpose genotype [21] or through specialized clones of recurrent origins (frozen niche variation [22, 23]). The above mechanisms likely operate jointly in the formation of geographic polyploidy and/or geographic parthenogenesis, and resolving their relative contributions requires empirical studies that link the genomic origin of polyploid lineages with eco-evolutionary studies of ploidy difference [24].

Polyploidy is far more common in plants than in animals [25], producing a marked bias in research effort [26, 27]. In angiosperms, roughly a third of extant species are polyploid [28] and cytogeographic surveys have been integrated with phylogenomic reconstruction of polyploid origin [29, 30] and tests of ecological consequences [31, 32]. In animals, polyploids comprise only ∼0.8% of species even in well-sampled taxa [2], and no comparable synthesis exists. In vertebrates, genetic origins of polyploid lineages have been reconstructed only recently (fish [33–35]; amphibians [36]; reptiles [37]). In invertebrates, polyploidy is documented in several phyla but systematically studied in only a few, such as crustaceans [38, 39] and earthworms [40]. In insects, polyploidy has long been recognized, particularly in weevils, where diploid to decaploid parthenogenetic forms have been reported [41–43]. However, the origins of these polyploids have not been reconstructed with genome-wide data, an approach that has been applied only in stick insects [44]. Distinguishing general ecological and evolutionary processes from taxonomic idiosyncrasy of well-studied clades therefore requires convergent evidence from phylogenetically independent origins.

Here, we studied the flightless weevil genus *Catapionus* Schoenherr, 1842 (Coleoptera: Curculionidae). In Japan, *Catapionus* weevils are univoltine: adults are found on Asteraceae plants, including thistles (*Cirsium* spp.) and butterbur (*Petasites* spp.), from May to August, and larvae develop in the soil on the roots of the same host plants [45]. Two morphologically similar species of uncertain phylogenetic relationship occur in Japan: *Catapionus nebulosus* (Motschulsky, 1860), formerly known as *C. gracilicornis* Roelofs, 1873 [46], which ranges from Sakhalin and the Kuril Islands through Hokkaido to Honshu, and *C. modestus* Roelofs, 1873, which occurs in central and western Honshu and Shikoku [47]. *C. nebulosus* has also been divided into several subspecies, but intraspecific phylogenetic relationships remain unresolved [48]. We therefore treat the two species together as the “*Catapionus nebulosus* species group” throughout this study. *Catapionus nebulosus* also includes all-female polyploid forms. Early karyological studies found mainly tetraploid and pentaploid forms in Hokkaido and northern Honshu [43, 49, 50], whereas diploid sexual individuals have been reported only from Honshu [51, 52]. Eggs laid by individually reared females of polyploid forms hatched at high rates [45], although parthenogenesis remained unconfirmed because the females’ mating history was not controlled. In addition, females of a bisexual subspecies near the Cape Tappi-zaki were described as resembling those of the all-female forms [47]. The sexual and all-female forms may therefore have been difficult to distinguish morphologically, making their distributions unclear. Their phylogenetic relationships and the evolutionary origin of the all-female polyploid form remain unknown.

We aimed to clarify the origin of the all-female polyploid form of *C. nebulosus* and the processes that produced its northern distribution. Because the range-wide distribution of the all-female form was unknown, we characterized the geographic distribution of sex ratio across Japan. We then determined whether the all-female polyploid populations form a single lineage or several independent lineages. Parthenogenesis in animals often arises through hybridization [20, 37], so we also tested whether hybridization contributed to the ancestry of the all-female form. Because parthenogenesis had not been confirmed, we reared females in isolation from the egg stage over two generations. We also tested whether females from populations with males require fertilization. Several hypotheses predict that a polyploid parthenogen establishes where sexual relatives are sparse [13, 14, 18, 19]. We therefore characterized trends in SNP-based heterozygosity and larval density toward higher latitudes in the sexual populations.

## Methods

### Sample collection, sex ratio estimation

We surveyed 176 localities across Japan, from Hokkaido to Shikoku, between June 2014 and June 2025, collected 1,624 adults of the *C. nebulosus* species group from host plants including butterbur (*Petasites* spp.) and thistles (*Cirsium* spp.), and preserved them at −80 °C or −30 °C until DNA extraction. In our field surveys, adults were collected by visual search on host plants or by beating, usually over several hundred host plants within about 100 m. All adults found were collected to minimize potential bias in sex ratio estimates. We also examined museum and private collections: the Kushiro City Museum (22 specimens from 13 localities), the Kanagawa Prefectural Museum of Natural History (51 from 11), the Museum of Nature and Human Activities, Hyogo (196 from 36), and the collections of co-authors Matoba (142 from 54) and Sato (188 from 21). Coordinates of all records were rounded to the nearest minute of latitude and longitude (about 2 km), and records with the same rounded coordinates were treated as one locality. Records without coordinates were assigned to localities by place name. The final dataset comprised 2,223 adults from 294 localities (**Table S1; Figure S1**). Published and social media records were used only to guide field sampling and were not included in the analyses. Sex was determined by genitalic inspection. The locality-level female proportion was modeled against latitude with a beta-binomial GLM in the R package glmmTMB [53] in R v4.5.2 [54]. The number of females out of the total number of individuals examined at each locality was used as the response, and all 294 localities, including those with few individuals, were included in the model. Individuals and localities for the genetic analyses, the rearing experiment, and the larval survey were chosen only to cover the geographic range as evenly as possible. Genome size was measured only in individuals of the nuclear clade that includes the all-female lineage (**Supplementary Methods; Figure S2**).

### Mitochondrial phylogeography

Genomic DNA was extracted from a middle or hind leg of each specimen (**Table S2**) using the DNeasy Blood & Tissue Kit (QIAGEN, Hilden, Germany). Partial sequences of mitochondrial cytochrome *c* oxidase subunits I (COI) and II (COII) were amplified by PCR with TaKaRa Ex Taq (Takara Bio, Shiga, Japan). Information about primers and thermal cycling conditions is given in **Table S3**. Amplicons were purified with ExoSAP-IT Express (Thermo Fisher Scientific, Waltham, MA, USA) and Sanger-sequenced by Eurofins Genomics (Tokyo, Japan). Sequences were assembled in MEGA7 [55] and aligned with MAFFT v7 [56].

Bayesian phylogenetic analysis was conducted in BEAST v10.5.0 [57, 58] with COI and COII as separate partitions. Substitution models were selected under the BIC in ModelTest-NG [59]: TIM2+G for COI and TN93+G for COII (**Table S4**). Rate heterogeneity was modeled with a discrete gamma distribution (four categories [60]) without an invariable-sites parameter, as their joint estimation is unreliable at the intraspecific level [61, 62]. We used an uncorrelated relaxed lognormal clock, with the mean COI rate calibrated to 1.68% per Myr [63] under a normal prior and the mean COII rate estimated under a lognormal prior. A constant-size coalescent tree prior was used, with the starting tree constrained to enforce monophyly of *Dermatoxenus* Marshall, 1916, *Catapionus viridimetallicus* (Motschulsky, 1860), and the *Catapionus nebulosus* species group. MCMC chains were run for 300 million generations, with thinning at every 30,000 generations. Convergence was confirmed in Tracer v1.7.2 [64] by visual inspection and with effective sample sizes of > 200. Runs were combined in LogCombiner v10.5.0 and summarized as a maximum clade credibility tree in TreeAnnotator v10.5.0 [57]. Node support was reported as posterior probabilities and divergence times as median estimates with 95% HPD intervals.

For each major mitochondrial lineage, we reconstructed demographic and spatial dispersal history in BEAST v10.5.0. Effective population size through time was estimated with the Bayesian Skyride coalescent prior [65], and geographic diffusion was modeled simultaneously under the relaxed random walk model in continuous space with a Cauchy distribution [66]. A strict clock was applied to each partition. Substitution rates and the root height were assigned informative priors based on the posterior estimates from the preceding analysis. Sampling coordinates were jittered by ±0.01° at shared localities to avoid poor model performance. MCMC chains were run for 100 million generations, with thinning at every 10,000 generations. Convergence was confirmed as described above. We quantified range expansion across 900 post-burn-in trees with the R package seraphim [67]. This package computes weighted branch dispersal velocity by dividing the total great-circle distances spanned by all branches by their total duration. Posterior median and its 95% credible intervals (km Myr□¹) were reported for each lineage.

### Reference genome assembly and whole-genome comparison

We generated reference genome assemblies for *Catapionus nebulosus* (Akita Prefecture) and *C. modestus* (Shiga Prefecture) from PacBio HiFi reads. Both are morphological species within the species group, and we use the names only as labels for the two assemblies. For each species, whole-body tissues (excluding head and elytra) from three adults preserved at −80 °C were pooled for high-molecular-weight (HMW) DNA extraction with the NucleoBond HMW DNA kit (Macherey-Nagel, Düren, Germany); a single individual did not yield sufficient HMW DNA for library preparation. DNA fragments shorter than 4 kb were removed with the Short Read Eliminator XL kit (Pacific Biosciences, Menlo Park, CA, USA), and HiFi sequencing was performed by Novogene (Beijing, China).

After adapter trimming with HiFiAdapterFilt [68], reads were assembled with hifiasm v0.19.9-r616 [69] with --n-hap 6 for the six haplotypes of the pooled samples. Three rounds of purge_dups v1.2.6 [70] were applied to reduce haplotig redundancy. Contigs were fragmented into 2,000-bp segments and screened for non-eukaryotic contamination by blastn [71] against the NCBI nt database (downloaded 16 April 2022). Contigs with at least 50% non-eukaryotic best hits were removed. Assembly completeness at each stage was assessed with Benchmarking Universal Single-Copy Orthologs (BUSCO) v6.0.0 [72] in genome mode using the insecta_odb10 lineage dataset. The two assemblies were aligned with nucmer in MUMmer4 [73], using *C. nebulosus* as the reference and *C. modestus* as the query. One-to-one best matches (minimum alignment length 1 kb, minimum identity 80%) were retained with delta-filter. The retained matches were converted to UCSC chain format and used to generate liftover indices with LevioSAM2 [74].

### Variant calling of MIG-seq

Genome-wide SNPs were generated by MIG-seq [75] from 118 individuals (113 of the *C. nebulosus* species group and five *C. viridimetallicus*), using the DNA extracts described above. Libraries were prepared following [76] with modifications. The first PCR was run for 28 cycles instead of the published cycle number. The forward-primer tail was replaced with an anchor sequence that matched the second PCR design, in which the second PCR reverse primer carried a 6-bp index. Libraries were sequenced by Novogene on an Illumina NovaSeq platform with 150-bp paired-end reads.

Raw reads were demultiplexed with process_shortreads in Stacks v2.68 [77], adapter- and primer-trimmed with cutadapt v1.15 [78], and quality-filtered with Trimmomatic v0.39 [79]. To reduce reference bias from genetic divergence, reads were mapped under a reference-flow framework [80]: all reads were first mapped to the *C. nebulosus* genome with BWA-MEM v0.7.17 [81], and alignments with MAPQ ≥ 20 were retained. Unmapped and low-confidence reads were remapped to the *C. modestus* genome, and resulting alignments with MAPQ ≥ 20 from this second round were lifted to *C. nebulosus* coordinates with LevioSAM2 [74].

Because MIG-seq relies on multiplex PCR, read counts can be biased between alleles, making it difficult to reliably infer allele dosage from the read ratios of polyploid individuals. We therefore called genotypes as diploid-like hard calls without estimating allele dosage [82, 83]. Variants and genotypes were called with gstacks in Stacks v2.68 under its default model (marukilow; --var-alpha 0.01, --gt-alpha 0.05) [84]. This model considers only the two most frequent alleles at each site and assigns each individual the most likely of three diploid genotypes (0/0, 0/1, or 1/1), leaving the genotype missing when the likelihood ratio is not significant. All SNPs were therefore biallelic, as in a previous study of a mixed-ploidy complex [85]. Genotypes with a read depth below three were treated as missing. The resulting VCF was processed in PLINK v2.0.0 [86]. Individuals with a genotyping rate below 3% were removed (--mind 0.97). Sites were filtered at five thresholds (--geno 0.3, 0.4, 0.5, 0.6, and 0.7), and each downstream analysis was run separately on all five datasets. Results from --geno 0.5 were reported as the main result and the rest in Additional files 1 and 2. Each dataset was LD-pruned by removing one SNP from each pair with r² > 0.1 within 100-kb windows (--indep-pairwise 100kb 1 0.1). To test the sensitivity of the main results to depth and individual missingness, we repeated the heterozygosity analysis, SVDquartets, STRUCTURE, Dsuite, OrientAGraph, and DIYABC-RF with a minimum depth of five, a stricter individual missingness threshold (--mind 0.95), or both.

### Nuclear phylogenetic relationships and population structure

To visualize relationships among individuals without assuming a bifurcating history, we constructed Neighbor-Net networks from the five LD-pruned SNP datasets in SplitsTree6 [87], with 1,000 bootstrap replicates per dataset. To obtain a backbone topology of relationships among the resulting clusters, we inferred a species tree with SVDquartets [88] in PAUP* v4.0a169 [89], using 1,000 bootstrap replicates and 400,000 randomly sampled quartets per replicate. This topology was used for root inference and in the ABBA–BABA tests (see below). To infer the root position, we optimized branch lengths on each bootstrap topology in RAxML-NG v1.2.2 [90] under the GTR+G+ASC_LEWIS model.

Population structure was inferred in STRUCTURE v2.3.4 [91, 92], which performs well on mixed-ploidy data with unknown allele dosage [93]. Polyploid genotypes were encoded following Falush et al. [94] and Meirmans et al. [82]: we set PLOIDY = 5, so that each individual had five allele rows. For each individual, as many rows as its ploidy, estimated from genome size (**Supplementary Methods; Table S2**), were filled by repeating the observed alleles of each call, and the remaining rows were coded as missing. We also set RECESSIVEALLELES = 1 and coded the recessive-allele row as missing at every locus, so that allele copy numbers were treated as unknown. We ran the admixture model with correlated allele frequencies [91], without treating sampling locality as prior information, for K = 1–10 with 50 replicate runs per K. Each run used a 100,000 burn-in and 100,000 MCMC iterations. For each K, replicate runs were grouped into modes by their pairwise similarity (G′ ≥ 0.9), following the approach of CLUMPAK [95], and the runs of each mode were aligned and averaged with CLUMPP v1.1.2 [96], which was run from the R package pophelper v2.3.1 [97]. We refer to the mode containing the most runs as the major mode and to the others as minor modes. Rather than selecting a single optimal K, we report a range of K; the range was guided by the mean log-likelihood, ln Pr(X|K), the ΔK statistic [98], and the Puechmaille estimators [99], calculated with sampling localities as groups and a membership threshold of 0.5.

To account for clonal reproduction, we grouped mig-Np individuals into multilocus lineages (MLLs) by farthest-neighbor clustering of pairwise allelic mismatch, excluding missing sites. The clustering was cut just below the smallest pairwise distance between individuals of the sexual lineages [100]. We ran STRUCTURE with the same settings on a clone-corrected dataset that retained, for each MLL, the individual with the fewest missing genotypes. To check the number of clusters without assuming Hardy–Weinberg or linkage equilibrium, we performed PCA on the clone-corrected dataset and selected the number of clusters by k-means clustering with find.clusters() in the R package adegenet [101]. We retained all principal components, because reducing their number is unnecessary for k-means clustering [102], and chose the smallest K after which the Bayesian information criterion (BIC) increased, for K = 1–20 [103].

### Detection of hybridization and admixture

*C. viridimetallicus* shared too few homologous loci to serve as a reliable outgroup. We therefore inferred root placement with the minimal ancestor deviation (MAD) method [104], applying it to each SVDquartets bootstrap topology. MAD assigned each individual a probability of falling on the outgroup side in each bootstrap replicate; we averaged these probabilities across the individuals of each nuclear cluster to obtain a lineage-level frequency. The basal lineages identified by MAD method were then used as alternative outgroups in admixture analyses.

Allele-sharing asymmetries were tested with ABBA–BABA statistics in Dsuite v0.6 [105], using unpruned SNP datasets to maximize informative sites. Dtrios was run with 20 jackknife blocks under each candidate rooting assumption. The topology for each run was taken from the corresponding SVDquartets species tree. *f*-branch statistics were computed with Dsuite Fbranch and visualized with dtools.py.

Admixture graphs were inferred with OrientAGraph [106] under each candidate rooting assumption and all five SNP datasets. Migration edges from 0 to 10 were evaluated, with block size varied from 8 to 57 SNPs (50 runs per edge count). The optimal edge count was selected with the R package OptM [107]. At the optimal edge count, 100 jackknife and 1,000 bootstrap replicates were performed with block size fixed at 10 SNPs.

Alternative origin scenarios for the all-female lineage were compared with approximate Bayesian computation and random-forest model choice in DIYABC-RF v1.1.54 [108]. Models were evaluated under four rooting scenarios: the three candidate rootings and a root-agnostic set. Each scenario was crossed with all five SNP datasets, for 20 combinations in total. Allele frequencies were summarized by lineage with a minimum allele frequency threshold of 1%. Each model was simulated 10,000 times. Model selection used abcranger v1.16.69 [109], which grew a random forest of 2,000 trees in each of 10 replicate runs. Uniform priors ranged from 1,000 to 10 for effective population sizes, from 10 to 10 generations for divergence times, and from 0.01 to 0.99 for the admixture rate.

### Rearing experiment to confirm parthenogenesis

We conducted a rearing experiment that controlled mating history to test whether virgin females of the all-female populations produce fertile daughters and whether females of the sexual populations require mating to produce viable eggs. In 2021, adult females were collected at four localities where only females were found and at five localities where both sexes occurred (**Table S1**). We also collected the only female found at AO-03, where one male was also collected. Genotyping assigned this female to the lineage found at all-female localities (see Results), so she was grouped with the females from those localities. Some of the field-collected mothers had been included in the molecular and flow cytometry analyses (**Table S2).** Each female was reared individually with butterbur (*Petasites* spp.) at 20 °C under a 16L:8D photoperiod and was allowed to oviposit. Hatched larvae were reared on carrot and burdock root at 20 °C, held at 4 °C from December to March, and returned to 20 °C until adult emergence. At some localities, larval rearing containers were labeled only by locality, so the mothers of the resulting F1 females are unknown. Rearing in isolation from the egg stage to adult emergence ensured that all F1 adults, which emerged in 2022, were virgin. We then assessed reproduction in 39 virgin F1 females, which developed from at least 182 individually reared larvae. F1 females from localities with both sexes were kept unmated for 15–23 days and then paired with an F1 male from the same locality for 24 h (mating treatment). Eggs laid before and after the mating treatment were assigned to the unmated and mated phases, respectively. For each female, hatching success in each phase was calculated as the number of hatched larvae divided by the number of eggs laid. F1 females from all-female localities and AO-03 were kept unmated throughout, and their F2 offspring were reared to adulthood under the same isolated conditions. For these unmated F1 and F2 females, hatching success was calculated in the same way.

### Latitudinal trends in individual heterozygosity and larval density

Individual heterozygosity was estimated from the five unpruned SNP datasets with PLINK v2.0.0, as the proportion of genotyped sites that were heterozygous. Under random mating, the expected heterozygosity of an individual equals that of its population, so we treat individual heterozygosity as a measure of population genetic diversity [110]. We excluded the all-female polyploid lineage to restrict the analysis to the sexual lineages. Sexual individuals of *C. nebulosus* and *C. modestus* are diploid, as shown by previous karyological studies [51, 52] and by our chromosome observations (**Figure S3**). The analysis was further restricted to localities south of 39°04′N, where the all-female lineage has not been found in our survey. We modeled the relationship between latitude and heterozygosity with a beta-binomial GLMM in glmmTMB. We specified the numbers of heterozygous and homozygous sites in each individual as the response, standardized genotyping rate as a covariate, and locality as a random intercept. The linear model included latitude as a fixed effect, and the quadratic model included latitude and its square. Null, linear, and quadratic models were compared by AIC and likelihood-ratio tests.

We surveyed larval density at 21 localities from Hokkaido to Fukui Prefecture in autumn 2021, starting at the northernmost locality and proceeding southward. Latitude was therefore partly confounded with sampling date and potentially with host plant species. At each locality, perennial Asteraceae host plants were selected at random, and a 30-cm soil cube centered on the root zone of each plant was hand-sorted for weevil larvae. Samples collected from butterbur (*Petasites* spp.) were excluded because they may contain larvae of other Entiminae species such as *Scepticus* sp. [111–113]. The analysis was restricted to localities south of 39°04′N, where the all-female lineage has not been found in our survey. We modeled the relationship between latitude and larval abundance with a GLMM in glmmTMB. We specified the number of larvae in each soil cube as the response, latitude as a fixed effect, and locality as a random intercept. Poisson and negative binomial models were compared by AICc, and overdispersion and zero inflation were checked with simulated residuals in the R package DHARMa [114]. Linear and quadratic models were compared by a likelihood-ratio test.

## Results

### Geographic variation in sex ratio

To characterize the latitudinal distribution of sex ratio, we determined the sex of all 2,223 adults from 294 localities in Japan (**Figure 1; Table S1**). A beta-binomial regression of the locality-level female proportion on latitude returned a strongly positive slope (β = 0.451, SE = 0.039, *z* = 11.47, *P* < 0.001; **Figure 2**). The odds of an adult being female increased about 1.57-fold per degree of latitude. South of 39°N, males were detected at 125 of 148 localities (84.5%). The remaining 23 localities yielded only females, but each had five or fewer adults. At these sample sizes, an all-female sample does not deviate significantly from a 1:1 sex ratio (two-sided exact binomial test, *P* ≥ 0.0625), so these localities are consistent with chance sampling under a balanced sex ratio. In contrast, north of 39°N, 131 of 146 localities (89.7%) yielded only females (**Figures 1 and 2; Table S1**). At each of the 24 northern female-only localities with six or more adults, the same test rejected a 1:1 sex ratio (*P* ≤ 0.031). At least at these localities, the absence of males is unlikely to result from small sample sizes. At a regional scale, all 304 adults collected from 85 localities in Hokkaido were female (**Table S1**). In northern Honshu (39–42°N), males were detected at 15 localities, mostly in the western part of the region. Most of these localities were near the Cape Tappi-zaki at the northern tip of Honshu or in the Shirakami Mountains on the Akita–Aomori boundary (**Figures 1 and S1**).

**Figure 1.**
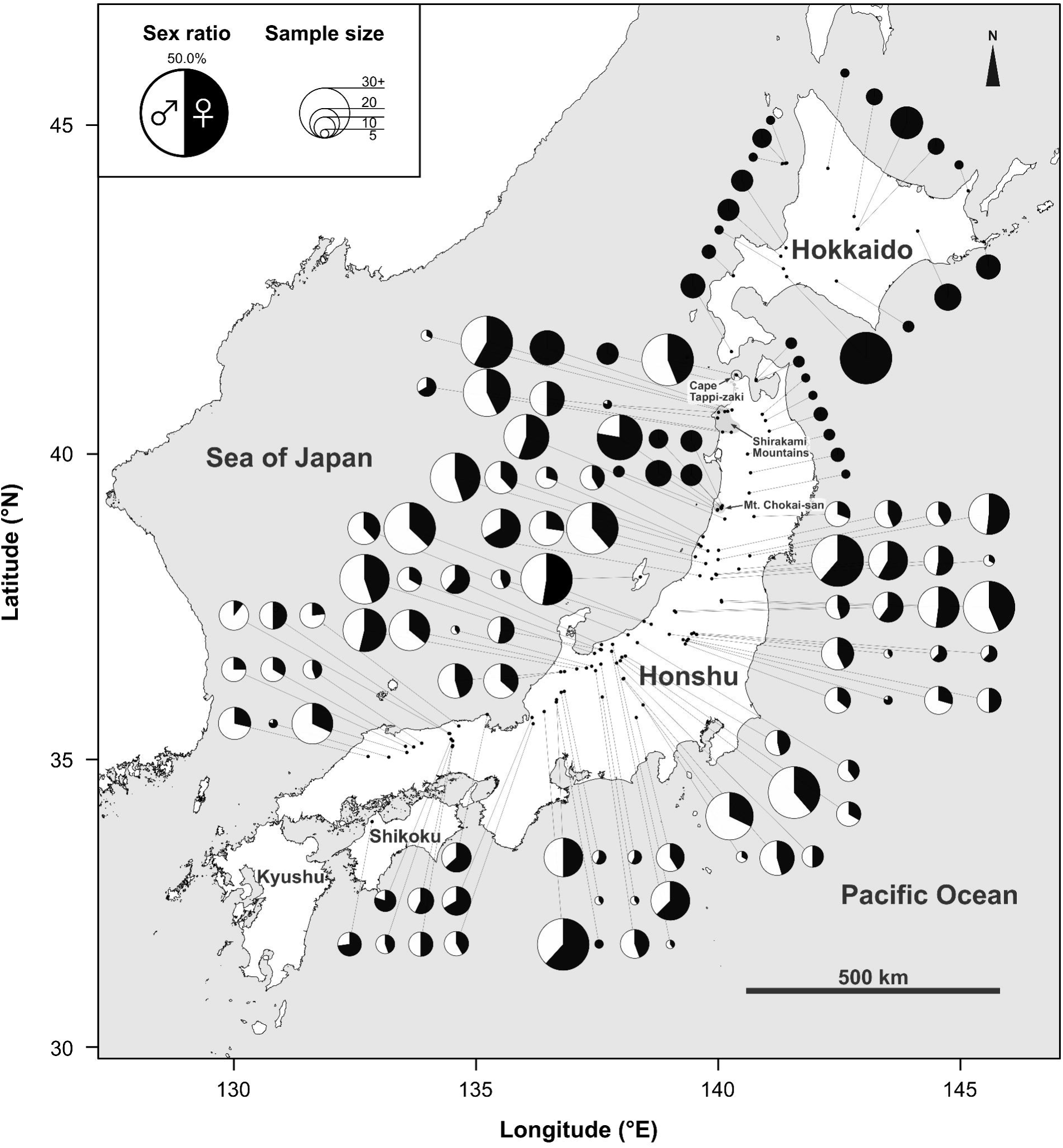
Geographic variation in sex ratio across Japanese populations of the *Catapionus nebulosus* species group. Only the 122 of 294 localities with five or more individuals are shown. Each pie chart shows the proportion of females (black) and males (white), with its size proportional to the number of individuals. Locality codes are given in Figure S1 and Table S1.

**Figure 2.**
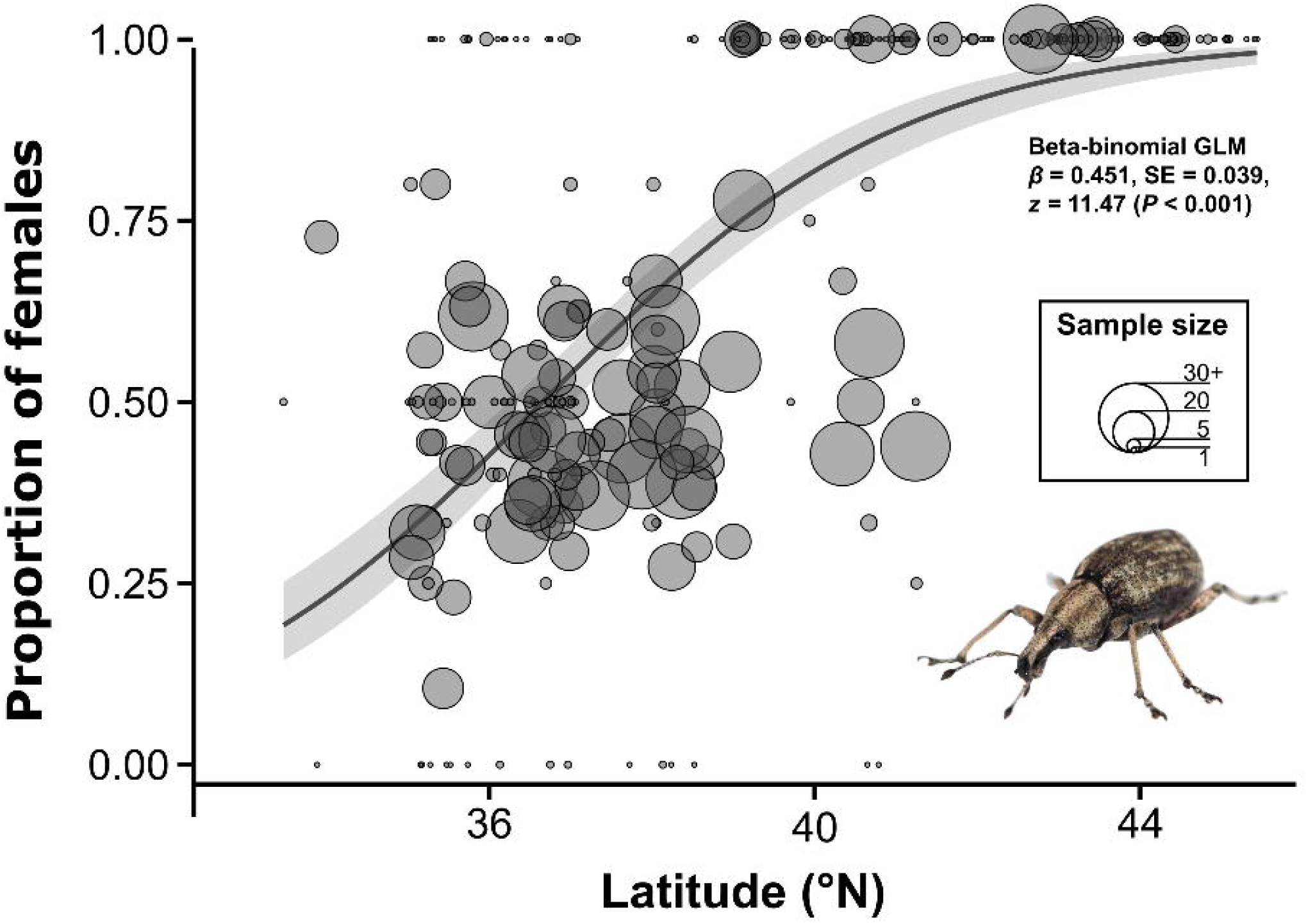
Relationship between latitude and sex ratio in the *Catapionus nebulosus* species group. Each circle represents one of the 294 localities, with its size proportional to the number of individuals. The curve shows the fit of a beta-binomial GLM to all localities, with shading indicating the 95% confidence interval.

### Mitochondrial phylogeny and geographic distribution

To examine the maternal origin of the all-female lineage, we reconstructed the mitochondrial phylogeny of the *C*. *nebulosus* species group, with the congener *C. viridimetallicus* and *Dermatoxenus* as outgroups (**Figure 3**). The concatenated COI and COII alignment was 1,235 bp long and included 177 individuals of the *C. nebulosus* species group, 26 of *C. viridimetallicus*, and 4 of *Dermatoxenus*. The phylogeny recovered nine well-supported, geographically structured clades along a northeast–southwest axis, which we named with the prefix “mt-” to distinguish them from the nuclear clusters defined later. Individuals that did not fall into any of these clades formed a paraphyletic assemblage, mt-X, confined to inland central Hokuriku. mt-Np was the most widely distributed, extending from Hokkaido southward through inland areas of the Tohoku region, with its southern range margin near Mt. Chokai-san (YG-02, 39°04′N). mt-N1 was restricted to the Sea of Japan coast of northern Tohoku, whereas mt-N2, mt-N3, and mt-N4 were confined to southern Tohoku. mt-C extended from southern Tohoku to eastern Hokuriku along the Sea of Japan coast, including Sado Island. mt-S1 and mt-S2 overlapped from central to western Hokuriku, and mt-S1 also occurred on Sado Island. mt-F extended from western Hokuriku southwestward across the mountains of Chugoku and Shikoku. Neighboring lineages were typically parapatric rather than separated by clear geographic gaps (**Figures 3A and 3B**).

**Figure 3.**
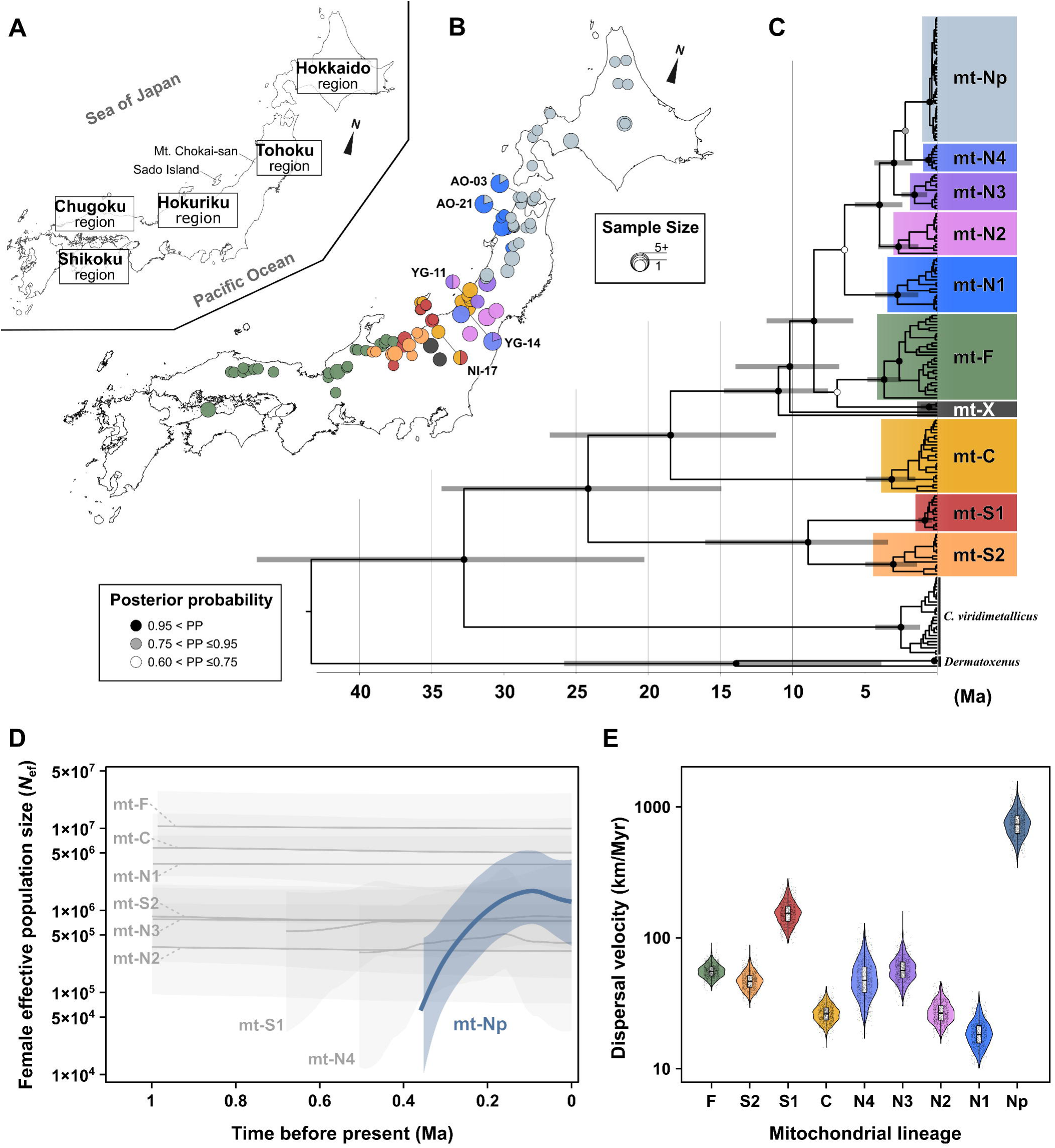
Mitochondrial phylogeny, demographic history, and spatial expansion of the *Catapionus nebulosus* species group. (A) Map of Japan showing the regions and places mentioned in the text. (B) Geographic distribution of the mitochondrial lineages. Each pie chart shows the lineage composition at one locality, with its size proportional to the number of individuals sequenced. Colors follow panel C. (C) Time-calibrated Bayesian tree of COI and COII (1,235 bp) from 177 individuals of the *C. nebulosus* species group, 26 of *C. viridimetallicus*, and four of *Dermatoxenus*. Node symbols denote posterior probability, and gray bars indicate 95% HPD intervals of node ages. (D) Bayesian Skyride reconstruction of female effective population size of mt-Np (colored) and the other lineages (gray). Lines and shading show the posterior median and 95% HPD interval. Individual lineages are shown in Figure S4. (E) Dispersal velocity of each mitochondrial lineage under a continuous phylogeographic model, from 900 posterior trees. Boxes show the median and interquartile range, and whiskers extend to 1.5 times the interquartile range. Values are given in Table S7.

Males were found in all lineages except mt-Np (**Table S2**). Genome size measurements showed that individuals of mt-Np were polyploid, whereas those of mt-N1 to mt-N4 were diploid, except for one tetraploid-like female of mt-N1 (Cneb1362) (**Supplementary Methods; Figure S2; Table S2**). mt-Np was monophyletic (PP = 1.00; **Figure 3C**), indicating that its individuals share a common maternal ancestor.

Divergence times estimated under a relaxed lognormal clock calibrated at 1.68% per Myr placed the tMRCA of *Catapionus* in Japan at 31.97 Ma (95% HPD: 20.31–47.12 Ma) and that of the *Catapionus nebulosus* species group at 23.60 Ma (95% HPD: 14.97–34.35 Ma; **Figure 3C; Table S5**). The closest sexual relative of mt-Np was mt-N4, restricted to inland southern Tohoku near Mt. Chokai-san (PP = 0.80). The tMRCA of mt-Np, 0.46 Ma (95% HPD: 0.25–0.79 Ma), was the youngest among the major lineages (0.53–3.60 Ma, **Table S5**). This date represents the age of the most recent common maternal ancestor of the sampled sequences, not necessarily that of the origin of the all-female lineage.

Bayesian Skyride reconstruction showed a marked expansion only in mt-Np. Its median female effective population size increased roughly 29-fold over the sampled timeframe, from 60,120 to 1,718,000 (**Figures 3D and S4; Table S6**). The lower limit of the 95% HPD interval at about 0.1 Ma exceeded the upper limit at the time of the most recent common ancestor (**Figure S4**). This supports a substantial increase in female effective population size, even after accounting for the uncertainty of the reconstruction. Part of this increase may reflect growth from the founding female(s). All sexual lineages remained stable over the same period, and their median female effective population size changed by no more than 32% (**Table S6**). Continuous phylogeographic analysis showed the same contrast in dispersal rate. The weighted branch dispersal velocity of mt-Np was 740.5 km Myr□¹ (95% CI: 467.7–1149.6), roughly five times higher than the fastest sexual lineage, mt-S1 (154.0 km Myr□¹; range across sexual lineages: 18.3–154.0 km Myr□¹; **Figure 3E; Table S7**). The all-female lineage expanded both in effective population size and in space at a pace unmatched by any sexual lineage of the species group.

### Reference genome assembly and whole-genome comparison

We generated de novo reference assemblies for *Catapionus nebulosus* (Akita Prefecture) and *C. modestus* (Shiga Prefecture) from PacBio HiFi reads (**Tables S8 and S9**). Each assembly was built from three individuals pooled from a single locality, using the same pipeline. After removal of non-eukaryotic contigs, the *C. nebulosus* assembly comprised 5,800 contigs (N50 = 305 kb; 1.086 Gb), with 95.7% complete BUSCOs (91.0% single-copy and 4.7% duplicated). The *C. modestus* assembly comprised 5,933 contigs (N50 = 431 kb; 1.831 Gb), with 98.2% complete BUSCOs (53.8% single-copy and 44.4% duplicated). The *C. modestus* assembly was thus 1.7 times longer and contained far more duplicated BUSCOs. The *C. modestus* source population was diploid (**Supplementary Methods; Figure S3**), consistent with previous reports for both species [115], so these differences do not reflect ploidy. Because these assemblies served only as mapping references, we did not examine its cause further. Pairwise alignment with nucmer recovered one-to-one hits for 90.4% of *C. nebulosus* contigs and 92.7% of *C. modestus* contigs, and this alignment provided the liftover chain used to translate read coordinates between the two assemblies in reference-flow mapping.

Because the *C. modestus* assembly may retain duplicated haplotypes, we checked whether this redundancy affected the genotypes. If duplicated contigs had split the two alleles at a locus, heterozygosity would be lower at SNPs on *C. modestus* contigs. Within individuals, heterozygosity at these SNPs was not lower than at SNPs on *C. nebulosus* contigs at any --geno threshold; it was similar at three thresholds and higher at two, including the primary threshold (odds ratio 1.31, 95% CI 1.19–1.43 at --geno 0.5; **Figure S5**). These results suggested that assembly redundancy did not substantially increase homozygous calls through allele splitting.

### MIG-seq data processing

We processed MIG-seq data from 118 individuals (113 *C. nebulosus* species group and 5 *C. viridimetallicus*). Sequencing produced 56,376,070 raw paired-end reads, of which demultiplexing retained 51,460,892 (436,109 ± 292,414 per sample, mean ± SD). Adapter trimming and quality filtering retained 77.2 ± 8.4% per sample. The two groups yielded similar numbers of trimmed reads per individual (255,939 and 228,668). Reference-flow mapping nonetheless retained a much higher fraction of reads in the *C. nebulosus* species group (40.4 ± 12.5%) than in the more divergent *C. viridimetallicus* (24.0 ± 8.9%).

Variant calling with gstacks in Stacks identified 168,427 SNP sites at a minimum read depth of three. Missingness across these unfiltered sites was high (per individual: median = 87.0%; range: 74.9–100.0%). We therefore filtered the data in PLINK v2.0.0. Removal of individuals whose genotyping rates fell below 3% (--mind 0.97) excluded 17 samples. Sites were then filtered at the five missingness thresholds (--geno 0.3–0.7, see Methods). Neighbor-Net used all 101 individuals; the remaining analyses used 99, excluding two samples that fell outside all clusters. All analyses except Dsuite used LD-pruned SNPs (4,307 at the primary threshold), whereas Dsuite used unpruned SNPs (12,025), because its block jackknifing accounts for linkage among sites. Mean genotyping rate decreased from 78.6% to 49.9% as filtering was relaxed. In the primary dataset (minimum depth of three, --mind 0.97, and --geno 0.5), individual genotyping rates ranged from 24.8% to 97.0% (median 65.6%; **Figure S6**), and every individual retained at least 2,963 called genotypes. The median read depth of the called genotypes ranged from 7 to 30 among individuals (median 15; **Figure S7**), so in every individual at least half of the genotypes were supported by seven or more reads. In this dataset, genotyping rate and depth were higher in individuals of the polyploid all-female lineage (medians 93.3% and 17) than in those of the sexual lineages (48.9% and 13.5). The higher genotyping rate in the polyploid lineage was consistent across all filtering conditions. Missing data and low depth were therefore not concentrated in the polyploid individuals, in which allelic dropout is of particular concern. Individual and SNP counts for each combination are given in **Table S10**, and the genotyping rate and read depth of each individual in **Table S11**. Results under stricter depth and individual missingness thresholds are reported in the relevant sections below.

### Nuclear phylogenetic structure

To test whether the all-female polyploid lineage identified in the mitochondrial data also forms a distinct group in the nuclear genome, we reconstructed relationships among individuals from the LD-pruned MIG-seq SNPs. Neighbor-Net analyses identified seven major clusters, denoted with the prefix “mig-” to distinguish them from the mitochondrial lineages. Cluster recovery was most robust at intermediate thresholds (--geno 0.4–0.6). Resolution declined at the extremes, where --geno 0.3 sharply reduced informative SNPs and --geno 0.7 admitted more missing data (**Figures 4A and S8; Table S12**). mig-F, mig-N1, mig-N2, and mig-N3 had consistently high bootstrap support across all five datasets (79.2–100.0%; **Table S12**). mig-Np formed a discrete, shallow, star-like grouping, with internal branches too short to yield stable bootstrap support (**Figures 4A and S8**). Cneb1186 (NN-08) fell within mig-C in four of the five datasets but outside it at --geno 0.4 (**Figure S8B; Table S12**). We treated this individual as a member of mig-C, because it grouped with mig-C in most analyses. Cneb0397 (NI-20) and Cneb1184 (GN-02) fell outside all clusters at all five thresholds. We refer to them as the mig-X assemblage. They were excluded from analyses of relationships among clusters (**Figures 4A and S8**). mig-S fragmented into weakly supported subclades only under extreme filtering (**Table S12**) and was treated as one cluster.

**Figure 4.**
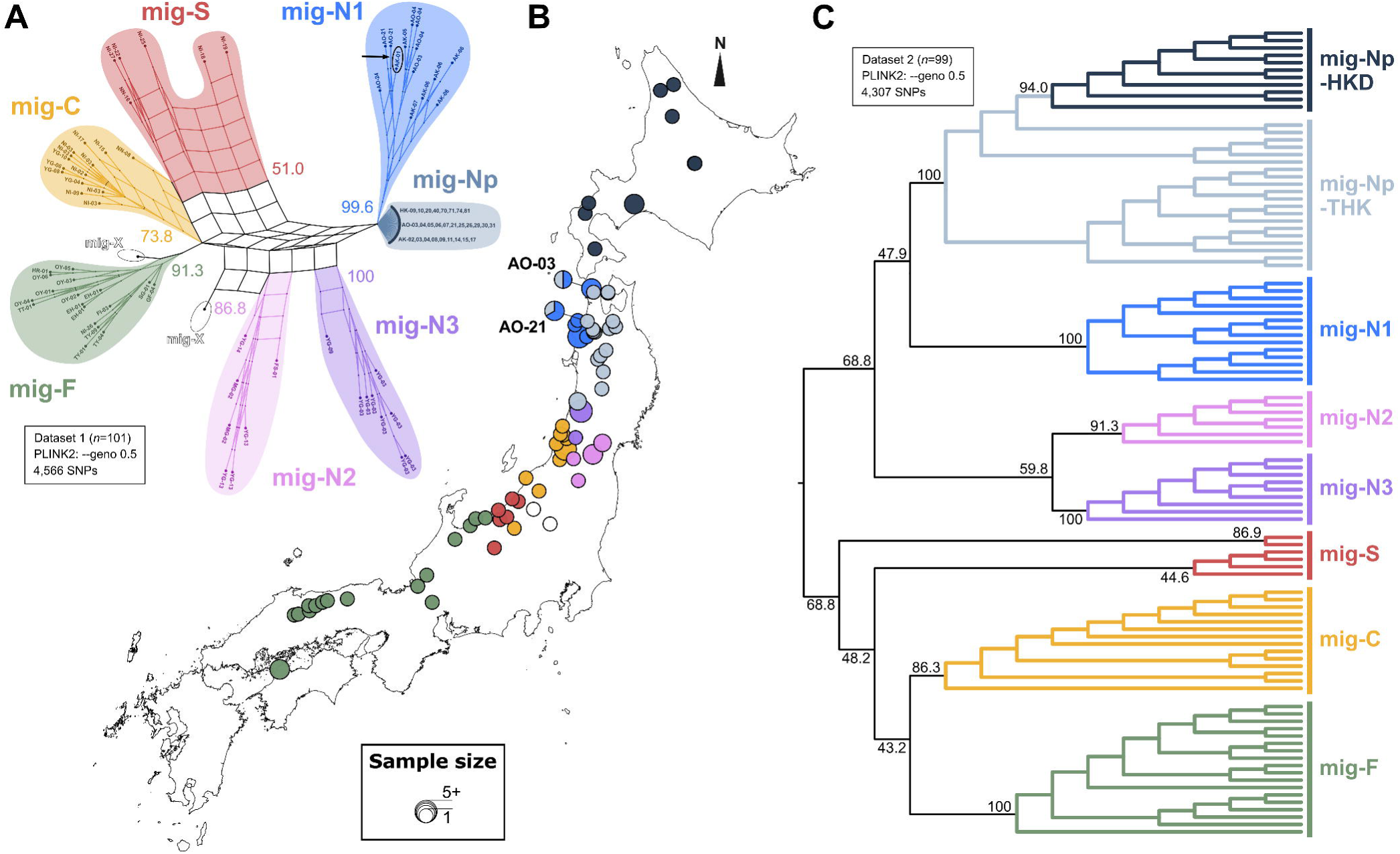
Nuclear phylogeny of the *Catapionus nebulosus* species group from MIG-seq SNPs. (A) Neighbor-Net split network of the --geno 0.5 dataset (n = 101; 4,566 LD-pruned SNPs). Tip labels give locality codes, and numbers give bootstrap support for the split subtending each lineage. The arrow marks Cneb1362 (AK-01), a tetraploid-like individual identified by flow cytometry. Other --geno thresholds are shown in Figure S8. (B) Geographic distribution of the nuclear lineages. Colors follow panel A. (C) SVDquartets tree of the --geno 0.5 dataset (n = 99; 4,307 LD-pruned SNPs), with bootstrap support at nodes. Other --geno thresholds are shown in Figure S9.

Four clusters corresponded one-to-one with a single mitochondrial lineage: all individuals of mig-F, mig-N1, mig-N3, and mig-Np carried mt-F, mt-N1, mt-N3, and mt-Np respectively, and no individual of these mitochondrial lineages fell outside the matching cluster (**Table S2**). The other three clusters mixed mitochondrial lineages. mig-C was the most heterogeneous, comprising mt-C alongside mt-N4, mt-S1, and mt-S2. mig-S combined mt-S1 and mt-S2 in equal numbers. mig-N2 comprised mt-N2 together with one mt-N4 individual from southern Tohoku (**Figure 4B**). Three mitochondrial lineages were split between two nuclear clusters each: mt-N4 between mig-N2 and mig-C, and mt-S1 and mt-S2 between mig-C and mig-S.

SVDquartets recovered the same seven clusters as Neighbor-Net, six of them as discrete clades. Across the five SNP datasets, mig-F and mig-N1 reached 100% bootstrap support, mig-Np 99.4–100.0%, mig-N3 99.3–100.0%, mig-C 69.2–98.5%, and mig-N2 51.8–93.4%, mostly higher than the corresponding Neighbor-Net values (**Figures 4C and S9; Table S13**). Only mig-S received weak support at every threshold (26.3–53.8%), consistent with its fragmentation in Neighbor-Net. Across 1,000 bootstrap replicates of each SVDquartets topology, the lineage that fell on the outgroup side shifted depending on the dataset (**Table S14**). At --geno 0.3, mig-F was sister to all remaining lineages (frequency 0.90). At --geno 0.4, mig-S occupied that position (0.78). At --geno 0.5–0.7, mig-F, mig-S, and mig-C formed a clade sister to the remaining lineages (1.00 for mig-F and mig-C, and 0.91–0.99 for mig-S). We treated these three lineages as candidate outgroups and ran the admixture analyses under each rooting independently. mig-N1, mig-N2, mig-N3, and mig-Np formed a clade (BS = 68.8–96.0%), within which mig-Np was sister to mig-N1 (47.9–91.7%; **Figures 4C and S9; Table S13**). All three candidate roots lay outside the northern clusters, so relationships among those clusters were unaffected by root placement. The mitochondrial data instead placed mt-N4 as the closest relative of mt-Np (PP = 0.80; **Figure 3**). Within mig-Np, a monophyletic Hokkaido group (hereafter mig-Np-HKD) was nested inside a paraphyletic Honshu assemblage (hereafter mig-Np-THK) rather than forming its sister group (**Figures 4C and S9**). Bootstrap support for mig-Np-HKD increased with SNP number. The clade was unresolved at --geno 0.3, where the fewest SNPs were retained, and reached 78.9%, 94.0%, 98.1%, and 99.0% at --geno 0.4, 0.5, 0.6, and 0.7. This topology places the Hokkaido populations within the Honshu assemblage.

We examined individual ancestry with STRUCTURE for K = 1–10, with ploidy assigned from genome size (**Figure S2; Table S2**). Individuals of mig-Np-HKD were treated as tetraploid, individuals of mig-Np-THK as pentaploid, and all other individuals as diploid. STRUCTURE was run on the five --geno datasets with a minimum depth of three and --mind 0.97, as well as on the --geno 0.5 datasets under the three stricter combinations of depth and individual missingness thresholds. Because potential clonal reproduction in the all-female lineage mig-Np could bias clustering through repeated multilocus genotypes, we also analyzed a clone-corrected version of the primary dataset (minimum depth of three, --mind 0.97, and --geno 0.5). The smallest distance between two individuals of the sexual lineages in this dataset was 0.0099, so we set the MLL threshold at 0.0098. With this threshold, the 31 mig-Np individuals formed 10 MLLs, and all 11 mig-Np-HKD individuals, from eight localities, belonged to a single MLL (**Figure S10**). The clone-corrected dataset therefore comprised 78 individuals.

Across all nine datasets, ΔK reached its maximum at K = 2, whereas the mean log-likelihood continued to increase up to K = 10 without a clear optimum (**Figure S11**). In the primary dataset, all four Puechmaille estimators, with sampling localities as groups, indicated seven clusters at K = 8 and did not increase further (**Figure S12C**). Across the nine datasets, the median-based estimators (MedMedK and MedMeaK) mostly leveled off at seven, and the maximum-based estimators (MaxMedK and MaxMeaK) at seven or eight (**Figure S12**). In the PCA of the clone-corrected dataset, the BIC of k-means clustering on all principal components reached its minimum at K = 7 (**Figure S13C**). These seven clusters matched the seven groups recovered by Neighbor-Net, except that one individual of mig-C, Cneb1186, was assigned to the mig-S cluster (**Figures S13A and S13B**).

For each K, replicate runs fell into one or more modes, and we describe the major mode unless stated otherwise. At K = 2, one ancestry component predominated in mig-F and mig-C and the other predominated in mig-Np. The remaining lineages carried both components in varying proportions, forming a gradient along the northeast–southwest axis (**Figure 5A**). mig-Np separated from every other lineage at K = 3 and remained distinct at all higher K. At K = 7, the two most frequent modes (27 and 20 of 50 runs) each failed to distinguish one sexual lineage, mig-N2 or mig-C, as a separate cluster (Figure S11C). At K = 8 (44 of 50 runs), each of the six sexual lineages and mig-Np formed its own cluster (**Figure 5A**). In the clone-corrected dataset, mig-Np shared one component with mig-N1 up to K = 5 and separated from it at K = 6 (**Figure S11F**). At K = 8 (45 of 50 runs), the same seven groups were recovered as in the full dataset (**Figure 5B**). mig-Np-THK and mig-Np-HKD were not separated in the major mode at any K. mig-Np-THK nonetheless carried a greater proportion of the component predominant in mig-N1 than mig-Np-HKD (median at K = 8: 4.2% vs. 0.0%; Wilcoxon rank-sum test, P < 0.001; **Figures 5A and S14; Table S15**). This difference was significant under all filtering conditions (P ≤ 0.008; **Figure S14; Table S15**).

**Figure 5.**
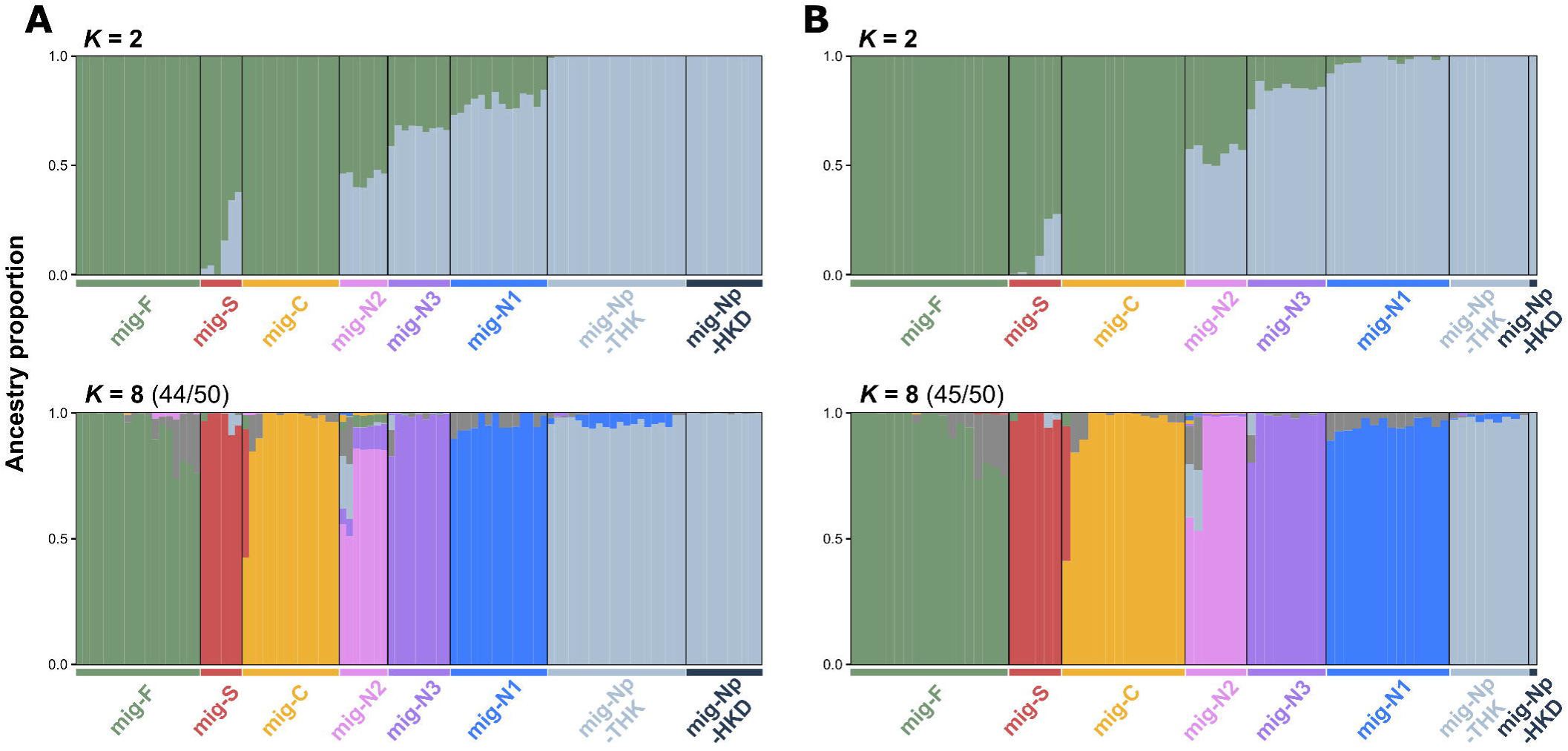
Population structure of the *Catapionus nebulosus* species group inferred with STRUCTURE (minimum depth of three, --mind 0.97, and --geno 0.5; 4,307 LD-pruned SNPs). (A) All individuals (n = 99). (B) Clone-corrected dataset with one individual per multilocus lineage (MLL; n = 78). Each bar represents one individual, and colors show the estimated ancestry proportions at K = 2 and K = 8. Colored bars below the plots indicate nuclear lineages. The major mode is shown with the number of runs it contains out of 50. K = 8 is the smallest K at which each of the six sexual lineages and mig-Np formed its own cluster; the eighth component (gray) did not predominate at any locality. Other K, minor modes, and other filtering conditions are shown in Figure S11.

### Hybrid ancestry of the all-female lineage

The mitochondrial and nuclear trees placed the all-female lineage differently: mt-Np was closest to mt-N4, whereas mig-Np was sister to mt-N1 (**Figures 3C and 4C**). Both hybridization between divergent lineages and incomplete lineage sorting can produce such discordance. We used three analyses to distinguish them: allele-sharing tests in Dsuite, admixture-graph inference in OrientAGraph, and coalescent model choice in DIYABC-RF.

To test for allele-sharing asymmetries consistent with introgression, we computed *D* and *f*-branch statistics in Dsuite under each candidate rooting. The *f*-branch statistics showed an excess of allele sharing between mig-S, or its ancestral lineage, and the ancestor of mig-Np (**Figure S15**). This signal appeared in all ten combinations of SNP dataset and rooting that did not assume mig-S as the outgroup. In analyses that assumed mig-S as the outgroup, this signal cannot be detected, because the outgroup cannot also serve as the P3 taxon. Under the primary filtering conditions, all 12 trios with mig-S as P3 and mig-Np-THK or mig-Np-HKD as P2 were significant (*f* -ratio = 0.07–0.12; **Figure S16**). Across the robustness conditions, these trios were significant in all 132 tests at --geno 0.5–0.7 and in 72 of 96 tests at --geno 0.3–0.4. An additional signal of excess allele sharing was detected between mig-C and mig-N2 in nine of the ten testable combinations, indicating that excess allele sharing was not restricted to mig-Np. Both this signal and the mig-S–mig-Np signal remained significant after Bonferroni correction (*D* = 0.227–0.860 for the mig-S signal; **Table S16**). These statistics detect excess allele sharing but do not establish the direction or timing of gene flow. We also tested whether allele sharing with other lineages differed between the Honshu and Hokkaido populations of mig-Np. Under the primary filtering conditions and all three rooting assumptions, mig-N1 showed significantly greater allele sharing with mig-Np-THK than with mig-Np-HKD (*D* = 0.43–0.57, *f* -ratio = 0.18–0.22; **signal c in Figure S15; Figure S17; Table S16**). Across the robustness conditions, this excess was significant in all 45 tests at --geno 0.4–0.7 and in 4 of 12 tests at --geno 0.3.

To estimate the number of admixture events and their positions, we fitted admixture graphs in OrientAGraph. OptM selected a single migration edge (*m* = 1) in every combination of SNP dataset and rooting (**Figure S18**). The recipient was the ancestor of mig-Np in all 33 analyses (**Figure S19; Table S17**). With mig-F or mig-C as the outgroup, the donor was mig-S in 18 of the 22 analyses and the ancestor of mig-S and mig-F in one. In the other three analyses, and in all 11 analyses with mig-S as the outgroup, the donor was placed on a branch ancestral to the recipient. Such placements are difficult to interpret and may reflect the limited identifiability of root placement and of edge direction in these models, particularly without a reliable outgroup [116]. In all 19 analyses in which the donor was mig-S or the ancestor of mig-S and mig-F, the migration edge was significant (*P* < 0.01; **Table S17**), and bootstrap support for the recipient node exceeded 96% at --geno 0.5–0.7 under the primary filtering conditions (**Figures 6A and S19**).

**Figure 6.**
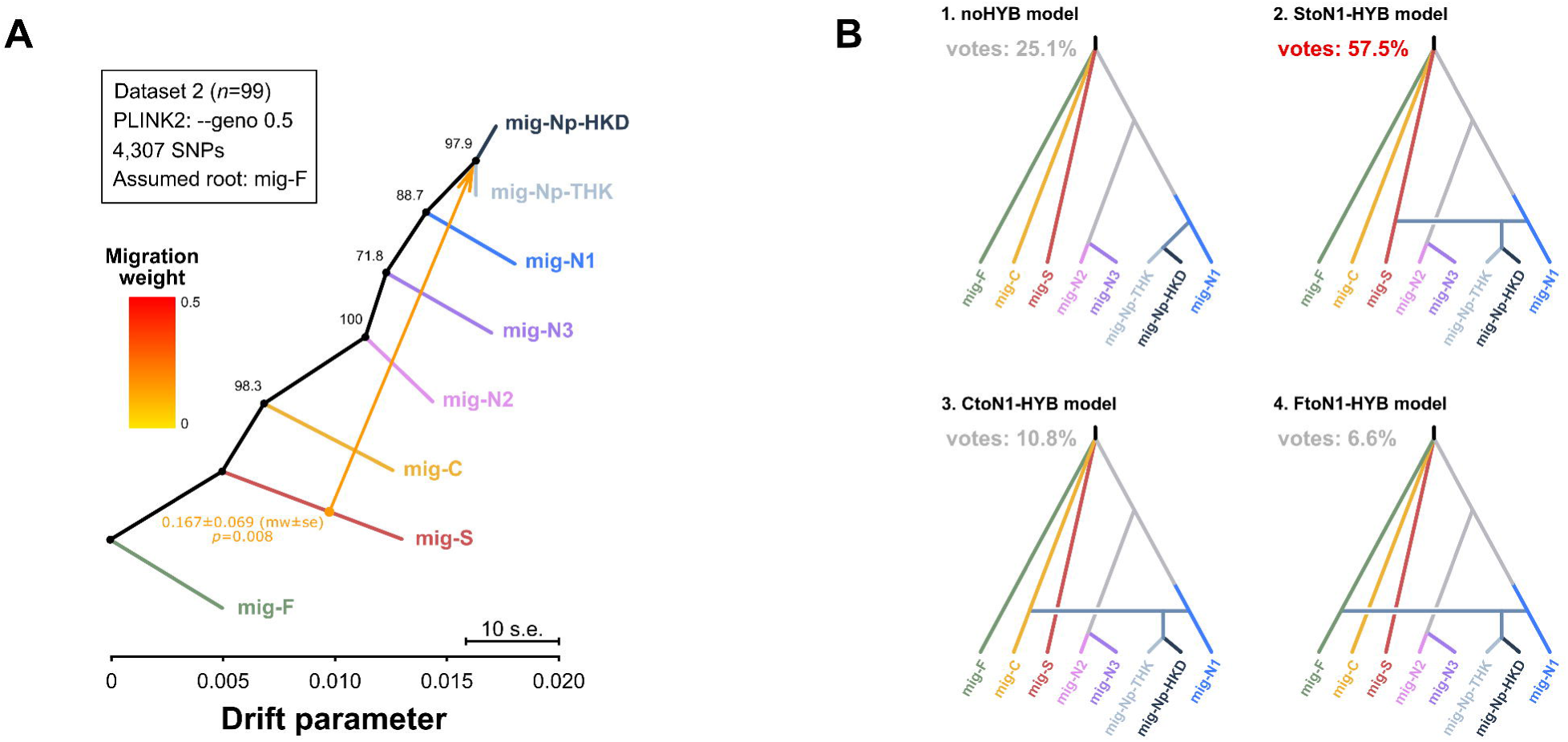
Tests of the hybrid ancestry of mig-Np. (A) Admixture graph of the --geno 0.5 dataset (n = 99; 4,307 LD-pruned SNPs) rooted with mig-F. The arrow shows the migration edge with its weight, standard error, and *P* value from jackknifing, and numbers at nodes give bootstrap support. The selection of the number of edges and the results for other datasets and rootings are shown in Figures S18 and S19 and Table S17. (B) Four origin scenarios compared with DIYABC-RF for the --geno 0.5 dataset under the root-agnostic setting, in which the branching order among mig-S, mig-F, and mig-C was not specified. Each scenario is labeled with the mean percentage of votes across 10 replicate random forests of 2,000 trees. Other datasets and rootings are shown in Figure S20 and Table S18.

To compare explicit scenarios while accounting for coalescence and population-size change, we performed approximate Bayesian computation with random-forest model choice in DIYABC-RF. We compared a non-hybrid model, in which the ancestor of mig-Np diverged directly from mig-N1, with three hybrid models in which it arose from mig-N1 crossed with mig-S, mig-F, or mig-C (**Figures 6B and S20**). We included alternative hybrid models involving mig-F and mig-C to test whether mig-N1 × mig-S was specifically favored over other hybrid models. Under the primary filtering conditions, the mig-N1 × mig-S model was selected in the majority of the 10 replicate runs in all 20 combinations of SNP dataset and rooting, with posterior probabilities of 0.596–0.736 (**Table S18**). Under the stricter conditions, it was selected in the majority of runs in 59 of the 60 combinations (**Table S18**).

### Experimental confirmation of parthenogenesis

To test whether the polyploid, all-female lineage reproduces by parthenogenesis, we reared females in isolation from the egg stage and recorded egg production and hatching in virgin females from localities of known lineage (**Figure 7A; Table S1**9). The 22 virgin F1 females came from four all-female localities and AO-03. At AO-03, the only female collected (Cneb0782) was assigned to mig-Np-THK and had a pentaploid genome size, whereas the male (Cneb0783) was assigned to mig-N1 (**Table S2**). Cneb0782 was the mother of two of the F1 females. All 22 virgin F1 females laid eggs that hatched, with a mean hatching success of 79.3% per female (range 42.4–97.2%). The two geographic groups gave similar results: 80.3% in mig-Np-HKD (*n* = 18) and 74.6% in mig-Np-THK (*n* = 4). We then reared three F2 females, the daughters of two of the mig-Np-HKD females, in the same way. Each laid eggs that hatched without mating (mean 67.7% per female, range 44.0–90.9%). All offspring that emerged as adults were female in both generations. Parthenogenesis was thus confirmed in the all-female lineage: unmated F1 females produced F2 females, which in turn reproduced without mating.

**Figure 7.**
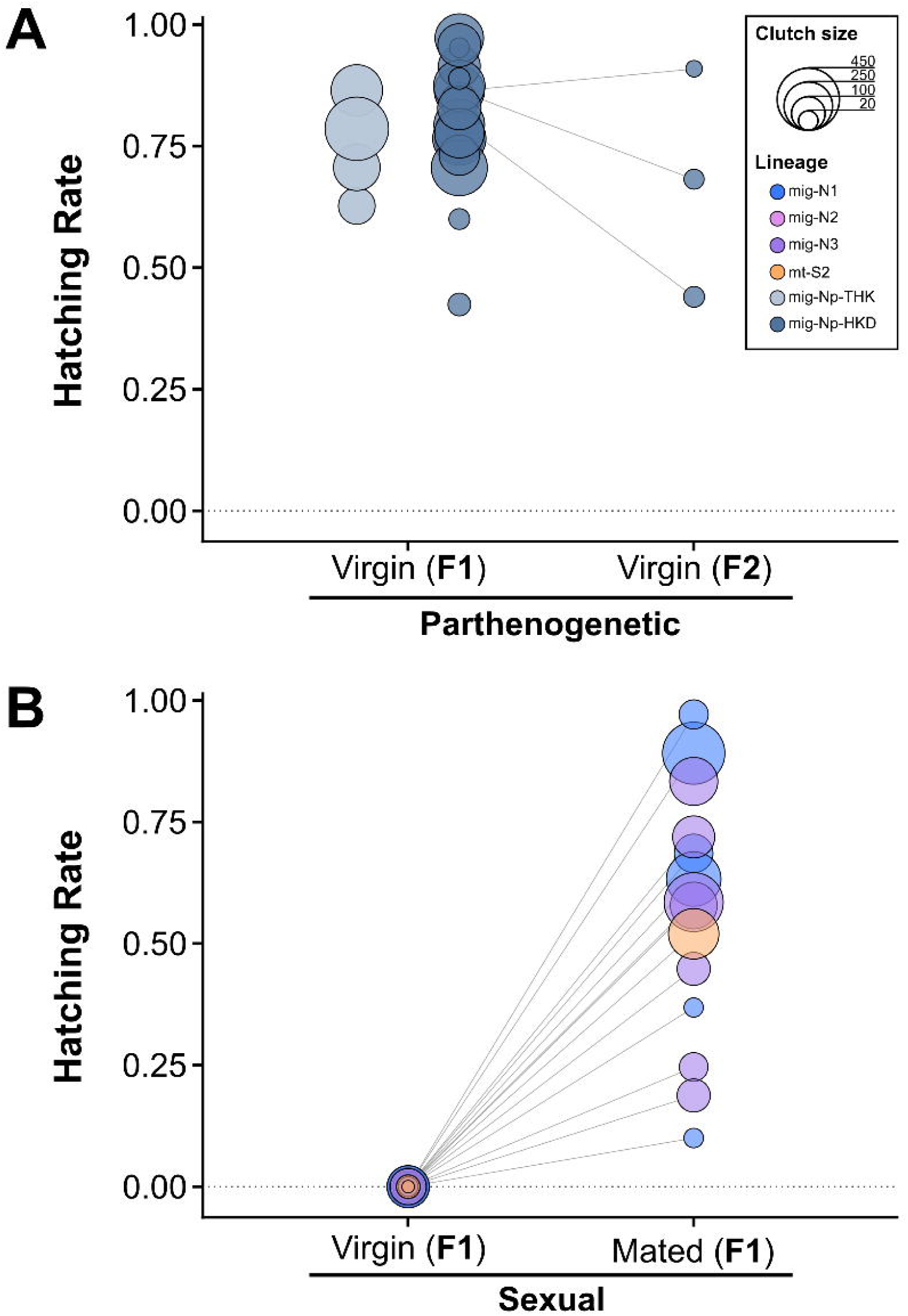
Hatching success of eggs laid by females of the *Catapionus nebulosus* species group under laboratory conditions. Each circle represents one clutch, with its size proportional to clutch size and its color indicating the lineage of the source locality. (A) Unmated females from all-female localities and AO-03 over two generations: 22 F1 females (18 from localities of mig-Np-HKD and four from localities of mig-Np-THK) and three F2 females from two F1 females of mig-Np-HKD localities. (B) Seventeen females from five localities with both sexes, representing four sexual lineages (mig-N1, mig-N2, mig-N3, and mt-S2), before and after pairing with a male. Lines connect the clutches of the same female. Counts for each female are given in Table S19.

Virgin females from localities with both sexes also laid eggs, but none hatched (**Figure 7B; Table S19**). The 17 virgin F1 females came from five localities of four sexual lineages (mig-N1, mig-N2, mig-N3, and mt-S2). At AO-04, both candidate mothers (Cneb0784 and Cneb0785) were assigned to mig-N1 and had diploid genome sizes (**Table S2**). At AK-06, four of the five candidate mothers (Cneb0760–Cneb0763) were assigned to mig-N1 and had diploid genome sizes; the fifth was not genotyped. Of the 17 females, 16 laid eggs, and none of their 978 eggs hatched. Of these 16 females, 14 were then paired with a male, and 13 laid eggs that produced larvae (mean hatching success of 53.3% per female, range 10.0–97.1%). Both sexes appeared among their offspring under the same rearing conditions. Egg development in the sexual lineages therefore required fertilization.

### Latitudinal patterns in heterozygosity and larval density

To examine the latitudinal distribution of genetic diversity in the sexual lineages, we calculated individual SNP-based heterozygosity from the MIG-seq data for sexual populations south of 39°04′N, the latitude of the southernmost locality of the all-female lineage (YG-02). In a beta-binomial GLMM with locality as a random effect and genotyping rate as a covariate, the quadratic model fit better than the null and linear models at the primary threshold (β = −0.060, SE = 0.025; *P* = 0.021, likelihood-ratio test). Under all 20 combinations of read depth, individual missingness, and site missingness thresholds, the quadratic model had the lowest AIC, and its quadratic term was negative and significant (**Table S20**). Genotyping rate had no significant effect on heterozygosity under any condition (*P* ≥ 0.24). Heterozygosity peaked near the center of the analyzed range and was lower toward both its northern and southern ends (**Figures 8A, 8B, and S21**).

**Figure 8.**
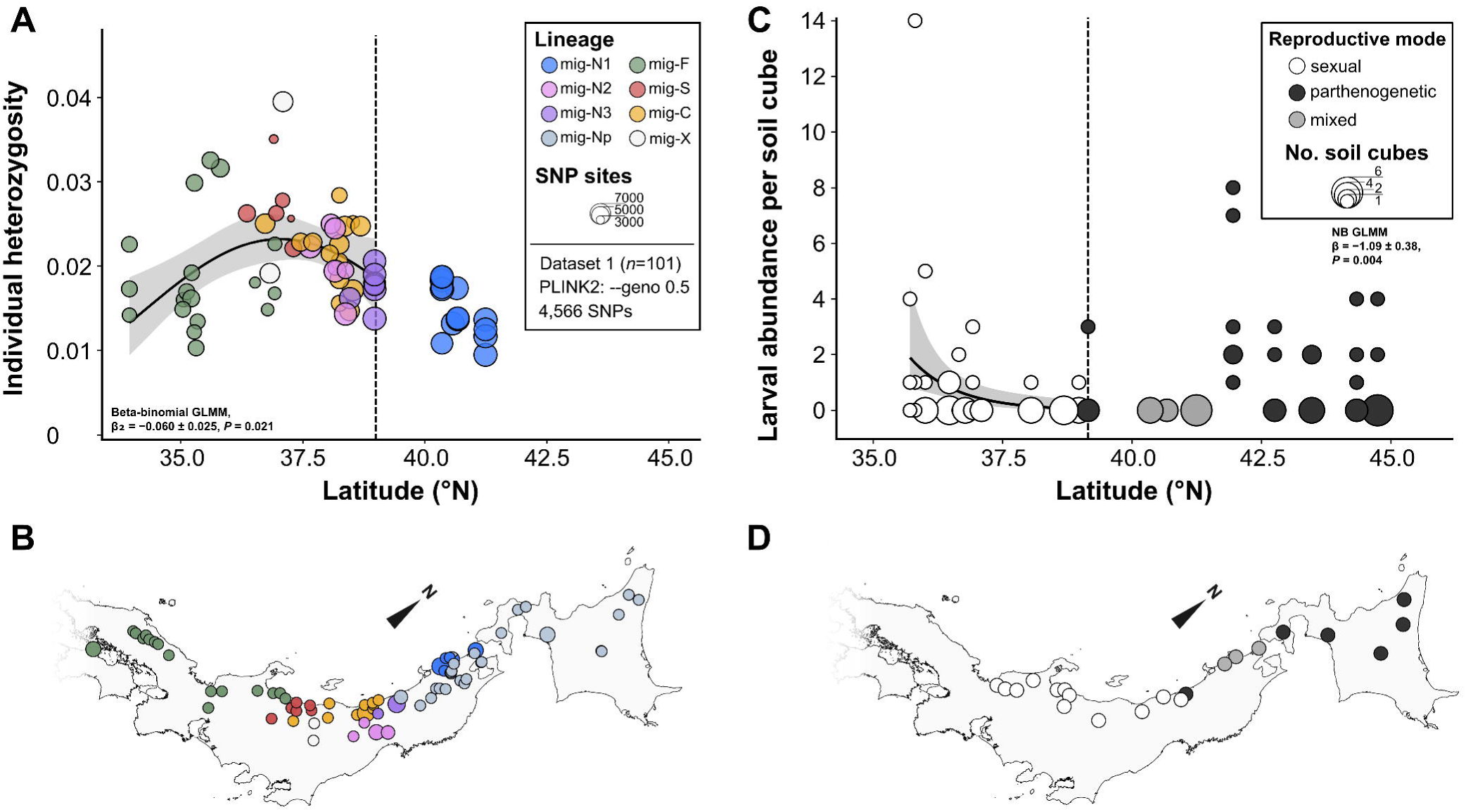
Latitudinal patterns in heterozygosity and larval density in the *Catapionus nebulosus* species group. (A) Individual heterozygosity (--geno 0.5 dataset) in relation to latitude. Each circle represents one individual, with its size proportional to the number of genotyped sites and its color indicating nuclear lineage; white circles indicate the mig-X assemblage. The curve shows the fit of a quadratic beta-binomial GLMM with genotyping rate held at its mean, fitted to sexual individuals south of 39°04′N (dashed line), the southernmost record of mig-Np. Other filtering conditions are shown in Figure S21. (B) Localities of the individuals in panel A and of mig-Np. (C) Number of larvae per soil cube in relation to latitude at 21 localities: 12 with sexual lineages only (white), three with both sexual lineages and mig-Np (gray), and six with mig-Np only (black). Circle size is proportional to the number of cubes with the same count at a locality. The line and shading show the fit of a negative binomial GLMM and its 95% confidence interval, fitted to the 12 sexual localities south of 39°04′N (dashed line). (D) Localities in panel C. Colors follow panel C.

To examine the latitudinal distribution of population density, we counted larvae in soil cubes taken from the rhizospheres of 97 host plants at 21 localities and collected 85 larvae in total (**Table S1**). No larvae were detected at six localities, although adults had been collected at these localities in summer. In the 12 localities of sexual lineages south of 39°04′N, the Poisson GLMM was overdispersed, and the negative binomial GLMM had a lower AICc (ΔAICc = 27.3). Residual diagnostics of the negative binomial GLMM showed neither overdispersion nor excess zeros (*P* = 0.38 and 0.96, respectively). A quadratic latitude term was not supported (*P* = 0.20, likelihood-ratio test). Larval abundance decreased with latitude (estimate = −1.09, SE = 0.38, *P* = 0.004; **Figures 8C and 8D**). Although heterozygosity and larval abundance were measured at different localities, both were lower toward the north within the analyzed range. By contrast, the six localities occupied only by mig-Np yielded 1.37 larvae per soil cube (48 larvae from 35 cubes), more than twice the density at the 15 localities with sexual lineages (0.60 larvae per cube; 37 larvae from 62 cubes).

## Discussion

Our results showed that the all-female lineage mig-Np, restricted to the northern part of the range of the species group, is polyploid, reproduces by parthenogenesis, and descends from a single common ancestor (**Figures 1, 3, 4, 7, and S2**). Three different analyses further indicated that hybridization with the divergent sexual lineage mig-S contributed to its ancestry (**Figures 6, S15, S16, S19, and S20**). *Catapionus* thus provides a rare animal example in which parthenogenesis, polyploidy, and a single origin of the lineage have all been shown.

### The cytogenetic origin of the polyploid lineage

Cold and fluctuating temperatures can raise the rate of unreduced gamete formation in plants and animals [1, 117, 118]. This may explain why polyploids are more frequent at high latitudes and in deserts [3–6, 119]. In *Catapionus*, low temperature induced polyploidized embryos [120, 121], but whether they can develop has not been tested. The restriction of mig-Np to the northern part of the range of the species group is consistent with a cold-driven origin. One tetraploid-like female, Cneb1362 (4.99 Gb per nucleus; **Figure S2**), was found within the otherwise diploid sexual lineage mig-N1 near the northern edge of its range. It carried a mitochondrial haplotype belonging to mt-N1 and fell within mig-N1 rather than in an intermediate position between mig-N1 and mig-Np (**Figure 4A**). Because mig-Np produces no males, a hybrid involving mig-Np would carry mt-Np. A backcross origin would require repeated backcrossing of polyploids to mig-N1 over several generations despite minority cytotype exclusion [13, 14]. Cneb1362 may therefore have arisen by genome duplication within mig-N1. This individual suggests that polyploidization can occur within sexual populations of *Catapionus* and that the resulting polyploid can develop into an adult.

All three admixture analyses indicated a genetic contribution from mig-S to the ancestor of mig-Np, and DIYABC-RF favored a cross between mig-N1 and mig-S among the four scenarios tested (**Figures 6, S15, S19, and S20**). In a clonal polyploid, however, allele-sharing tests cannot tell whether the contribution from mig-S predates the origin of parthenogenesis or was added after the lineage became parthenogenetic. Polyploid parthenogenetic lineages of hybrid origin occur in plants, algae, and animals [37, 44, 122, 123]. Hybridization between divergent lineages can disrupt meiosis, producing unreduced eggs that develop without fertilization [20, 21], and in whiptail lizards, parthenogens have arisen only from crosses between deeply divergent species [37]. The inferred parental lineages of mig-Np, mig-N1 and mig-S, are also deeply divergent **(Figure 4C**), which is consistent with this pattern. Crosses between sexual lineages that differ in genetic distance could test whether meiotic failure and unreduced egg production in hybrids depend on the divergence between the parental lineages.

Both the mitochondrial and nuclear trees showed that the two cytotypes of mig-Np form a single clade. The most parsimonious explanation is therefore a single polyploidization event from a diploid ancestor, after which ploidy variation arose within mig-Np. These cytotypes correspond to the tetraploid and pentaploid forms recorded earlier by Takenouchi [50]. The two cytotypes also differed in genome composition. The pentaploid Honshu populations (mig-Np-THK) shared more alleles with mig-N1 than the tetraploid Hokkaido populations (mig-Np-HKD) did (**Figure S17; Table S16**) and carried a small proportion of the STRUCTURE component predominant in mig-N1, which was nearly absent from mig-Np-HKD (median 0.3–4.2% vs. 0.0%; **Figures 5 and S14; Table S15**). In parthenogenetic weevils, ploidy is thought to increase stepwise through occasional fertilization by males of the same or related species [124]. Genome addition from mig-N1 males, which occur in northern Honshu but not in Hokkaido, would explain both the excess allele sharing and the difference in genome size, which corresponds to about one genome copy of mig-N1 (**Figure S2**). Alternatively, mig-Np-HKD may have arisen from a pentaploid ancestor by losing one chromosome set. Given the nested position of mig-Np-HKD within mig-Np-THK (Figures 4C and S9), this is the more parsimonious explanation of the topology. However, genome addition from mig-N1 to mig-Np-THK could also affect the inferred phylogenetic relationships and produce the same nested pattern. Genome loss would explain the difference in allele sharing only if the lost copy of every chromosome came from mig-N1, unless whole chromosome sets were lost at once [41]. We therefore consider genome addition more likely than genome loss, although our data cannot exclude the latter.

mig-Np thus combines polyploidy, parthenogenesis, and hybrid ancestry. Such combinations are well known in plants [10], and mig-Np provides an example in an animal. The combination of polyploidy and parthenogenesis may itself have allowed the polyploid lineage to persist. A newly arisen polyploid among diploids may be eliminated by minority cytotype exclusion [13, 14], but parthenogenesis can remove this barrier because it requires no mate [125]. Conversely, polyploidy may reduce the effects of deleterious mutations that accumulate in the absence of recombination [126]. Each trait may therefore facilitate the persistence of the other [24]. Consistent with this, we found no sexual polyploid population. Apart from Cneb1362, all individuals measured from the sexual lineages of the mig-N clade were diploid (**Figure S2**).

Understanding how polyploidy originated and how it relates to the co-occurring traits will require resolving the order of polyploidization, parthenogenesis, and hybridization, which our data could not determine. Despite range-wide sampling, we found no diploid parthenogen, the “missing link” reported in other weevils [42, 124] that would have helped clarify whether parthenogenesis preceded polyploidization [37]. Our MIG-seq genotypes were diploid-like biallelic calls without allele dosage [75, 83, 127]. They cannot resolve subgenomic composition, which will require haplotype-resolved whole-genome sequencing [128]. The lack of allele dosage information can also bias allele-frequency estimates and thus the magnitude of admixture estimates. However, such calls do not create alleles absent from an individual, so the shared alleles themselves are unlikely to be artifacts. Our inference of hybrid ancestry is supported by the consistent signal of a contribution from mig-S across all three admixture analyses. Biallelic SNPs from mixed-ploidy samples have likewise been used to detect interploidy introgression in a plant polyploid complex [85]. In addition, we did not examine reproductive endosymbionts. *Wolbachia* induces parthenogenesis in some arthropods and is associated with parthenogenesis in weevils of the tribe Naupactini [129–131], so screening mig-Np and its sexual relatives will be needed.

### Northward establishment and persistence of the polyploid parthenogens

The parental lineages of mig-Np, mig-N1 and mig-S, occur in Honshu and are absent from Hokkaido (**Figure 4**). Within mig-Np, the Hokkaido populations (mig-Np-HKD) are nested within the paraphyletic Honshu populations (mig-Np-THK; **Figures 4C and S9**). The polyploid parthenogenetic lineage therefore originated in Honshu, the southern part of its present range, and expanded northward into Hokkaido, where no sexual lineage of the species group occurs. The expansion was rapid in both spatial and demographic terms (**Figures 3D**, **3E, and S4**). All 11 individuals of mig-Np-HKD, from eight localities, belonged to a single MLL (**Figure S10**), consistent with colonization of Hokkaido by a few closely related founders within a short period.

In the sexual lineages south of the range of mig-Np, heterozygosity peaked near the center of the analyzed range and was lower toward its northern end, and larval density decreased toward the north (**Figure 8**). Genetic diversity reflects both demographic history and evolutionary processes, including mutation, genetic drift, selection, and gene flow [132]. MIG-seq targets anonymous regions between simple sequence repeats across the genome [75]. These loci are assumed to be largely neutral, and their mutation rates are likely similar among these closely related lineages. Gene flow is also limited in flightless weevils [133], so differences in heterozygosity among sexual populations may largely reflect differences in long-term effective population size. Heterozygosity would then reflect population size over many generations, whereas larval density reflects current abundance. The lower values of both toward the north therefore suggest that the northern sexual populations have been small for a long time, not only at present. Lower genetic diversity toward range margins is a common pattern and is often attributed to smaller population sizes there [134]. This interpretation should nevertheless be regarded as tentative, because the two variables were measured at different localities and heterozygosity can also be affected by admixture between lineages. Although our data cannot reveal how population sizes were distributed or how mig-Np spread when it originated, mig-Np at least currently occupies the region where sexual populations are sparse or absent (**Figures 1 and 8**). Several density-dependent hypotheses, which are not mutually exclusive, are consistent with this distribution. Minority cytotype exclusion is weaker where diploids are sparse [13, 14], and mate limitation in sparse populations favors reproduction without mates (Baker’s law [18, 19, 135]). Low density may also favor asexuality through sexual conflict [136]. A lineage at an expanding range front can also be fixed by drift alone [16, 17]. The density decline alone cannot separate these hypotheses. Testing them will require crosses between cytotypes with measurements of offspring fitness, and field measurements of the fertilization rate of sexual females.

The larval density of mig-Np was more than twice that at localities occupied by sexual lineages (**Figure 8C**). This difference may reflect higher reproductive output, lower mortality, or both. Parthenogenetic females produce no males and need not search for mates, which alone can double population growth [137]. Polyploidy and hybrid ancestry may lower mortality by broadening ecological tolerance [1, 20, 21]. Alternatively, mig-Np may have adapted locally after colonization, independently of polyploidy and hybrid ancestry, because the absence of recombination constrains but does not preclude adaptation [138]. In mig-Np, polyploidy, parthenogenesis, and hybrid ancestry are associated, and our design cannot separate their effects. Comparing fitness and environmental tolerance between sexual and parthenogenetic lineages under common conditions would show whether mig-Np performs better, but not which trait is responsible. Comparing the tetraploid and pentaploid cytotypes of mig-Np would help to test the effect of ploidy.

## Conclusions

We showed that the all-female populations of the *Catapionus nebulosus* species group form a single polyploid parthenogenetic lineage. Among the origin scenarios tested, hybrid ancestry from two divergent sexual lineages, mig-N1 and mig-S, was best supported. The tetraploid and pentaploid cytotypes of this lineage differed in genome composition, and the pentaploids may have arisen when tetraploid females gained an additional chromosome set from local sexual males. The order in which polyploidization, parthenogenesis, and hybridization arose remains unresolved. The lineage expanded northward and now occurs where sexual populations are absent or, near its southern limit, show low heterozygosity and larval density. This pattern is consistent with several mutually non-exclusive hypotheses, including mate limitation as described in Baker’s law. *Catapionus* thus provides a new animal example in which polyploidy, parthenogenesis, and hybrid ancestry coincide with a distribution biased toward higher latitudes, consistent with a pattern long recognized in plants. Further studies across animal taxa may help determine whether this combination reflects processes shared with plants or taxonomic idiosyncrasies.

## Declarations

### Ethics approval and consent to participate

This study did not require approval from an institutional ethics committee. It involved only field collection, non-invasive rearing, and molecular analysis of an invertebrate species (Coleoptera: Curculionidae) not subject to animal-ethics regulation under Japanese institutional or national frameworks.

### Consent for publication

Not applicable.

### Availability of data and materials

Mitochondrial COI and COII sequences are deposited in GenBank (BioProject PRJNA1507911; accessions PZ797942-PZ798144, PZ803735-PZ803937, PZ866819-PZ866822, and PZ882232-PZ882235). Raw PacBio HiFi reads used for the reference genome assemblies are deposited in the SRA (BioProject PRJNA1508320; accessions SRR40039004-SRR40039005). The corresponding genome assemblies of *Catapionus nebulosus* and *C. modestus* are deposited in GenBank under the same BioProject (accessions JCBZJO000000000 and JCBZJN000000000, respectively). Raw MIG-seq reads are deposited in the SRA (BioProject PRJNA1508350; accessions SRR40037598-SRR40037715).

### Authors’ contributions

Designed the research: SM, SD; Collected samples: SM, TS, IM; Analyzed the data: SM, PWH; Performed the statistical analysis: SM, SD; Wrote the manuscript: SM, SD; Contributed to revising the manuscript: SM, PWH, SD.

## Acknowledgements

We dedicate this study to the memory of the late Yasushi Takenouchi, whose discovery of polyploidy and parthenogenesis in *Catapionus* provided the foundation and primary motivation for this work. We thank Keisuke Tsuchiya (Kushiro City Museum), Kyohei Watanabe (Kanagawa Prefectural Museum of Natural History), and Kazutaka Yamada (University of Hyogo / Museum of Nature and Human Activities, Hyogo) for granting access to specimens in their care. We are grateful to Yasuhiro Tada (Kyoto Tachibana University) for advice on the ecology of the target species, and to Yutaka Okuzaki (Osaka Metropolitan University) for encouraging and supporting our field surveys. We thank Tatsumi Kudo (Kyoto University) and Seikan Kurata (University of Hyogo) for collecting specimens and Haruka Asanabe (University of Tokyo) for providing photographs of the target species. We also thank Takayoshi Yamamoto (Tokyo Gakugei University) for preparing the *Xenopus laevis* standard samples used in flow cytometry. We thank Chizuko Nishida (Hokkaido University) for teaching us chromosome preparation techniques. Haruka Maeta and Senga Touma provided valuable advice on figure design. We thank Hiroki Serikawa, Masatoshi Ohsawa, Junko Ishiyama, Yuko Ueda, Sorako (@cicindela_jp_), and delta_mu (@insects_birds) for permission to use their online observation records, which guided our field surveys.

