## Supplemental Figures for "Walking Alone to the North: The Origin and Historical Expansion of a Polyploid Parthenogenetic Lineage in a Weevil"

**Additional file 1: Supplementary information for:**

**Table of Contents:**

| **Additional file 1: Supplementary Methods** | Page 2 |
| --- | --- |
| **Figure S1** | Page 3 |
| **Figure S2** | Page 4 |
| **Figure S3** | Page 5 |
| **Figure S4** | Page 6 |
| **Figure S5** | Page 7 |
| **Figure S6** | Page 8 |
| **Figure S7** | Page 9 |
| **Figure S8** | Page 10 |
| **Figure S9** | Page 11 |
| **Figure S10** | Page 12 |
| **Figure S11** | Pages 13–21 |
| **Figure S12** | Page 22 |
| **Figure S13** | Page 23 |
| **Figure S14** | Page 24 |
| **Figure S15** | Page 25 |
| **Figure S16** | Page 26 |
| **Figure S17** | Page 27 |
| **Figure S18** | Page 28 |
| **Figure S19** | Page 29 |
| **Figure S20** | Page 30 |
| **Figure S21** | Page 31 |
| **References** | Page 32 |

**Additional file 1: Supplementary Methods**

**Genome size estimation**

To characterize the geographic distribution of ploidy, we estimated genome size (2C DNA content) by flow cytometry in 54 individuals of the nuclear clade mig-N, which comprises the parthenogenetic lineage mig-Np and the sexual lineages mig-N1, mig-N2, and mig-N3 (**Table S2**). Adult brain tissue was dissected in Galbraith's buffer [1] and homogenized in 200 µL of extraction buffer (CyStain PI Absolute T kit; Sysmex, Kobe, Japan). Each homogenate was mixed with 1.5 mL of propidium iodide staining solution from the same kit, filtered through a CellTrics 50 µm mesh (Sysmex), and stained for 30 min. Fluorescence from at least 1,000 nuclei per sample was acquired on a CyFlow Ploidy Analyser (Sysmex) and calibrated against *Xenopus laevis* (2C = 6.2 Gb) or chicken (*Gallus gallus*) blood (2C = 2.4 Gb) as internal standards.

Genome size followed a clear trimodal distribution (**Figure S2**). The 27 individuals of the sexual lineages, excluding Cneb1362, included both sexes and formed a diploid class (2C = 2.36 ± 0.18 Gb, range 2.02–2.74 Gb; mig-N1, n = 13; mig-N2, n = 5; mig-N3, n = 9). All measured individuals of mig-Np were female and polyploid. The eight from Hokkaido (mig-Np-HKD) formed a tetraploid class (4.30 ± 0.20 Gb, range 4.05–4.66 Gb), and the 18 from Tohoku (mig-Np-THK) a pentaploid class (5.49 ± 0.19 Gb, range 5.14–5.82 Gb). One further female of mig-N1 (Cneb1362, AK-01) had a genome size of 4.99 Gb, more than twice that of any other individual of mig-N1 (**Figure S2**).

**Chromosome observation**

For chromosome observation, testes from three males of the Shiga population used for reference genome assembly (see “Reference genome assembly and whole-genome comparison” in the main text) were dissected and processed using the air-drying method [2]. Tissues were treated with hypotonic solution, fixed, and stained with 3% Giemsa solution for 10 min before microscopic observation.

Mitotic spreads of spermatogonia from the Shiga population consistently showed 22 chromosomes, confirming a diploid chromosome number of 2n = 22 (**Figure S3**).

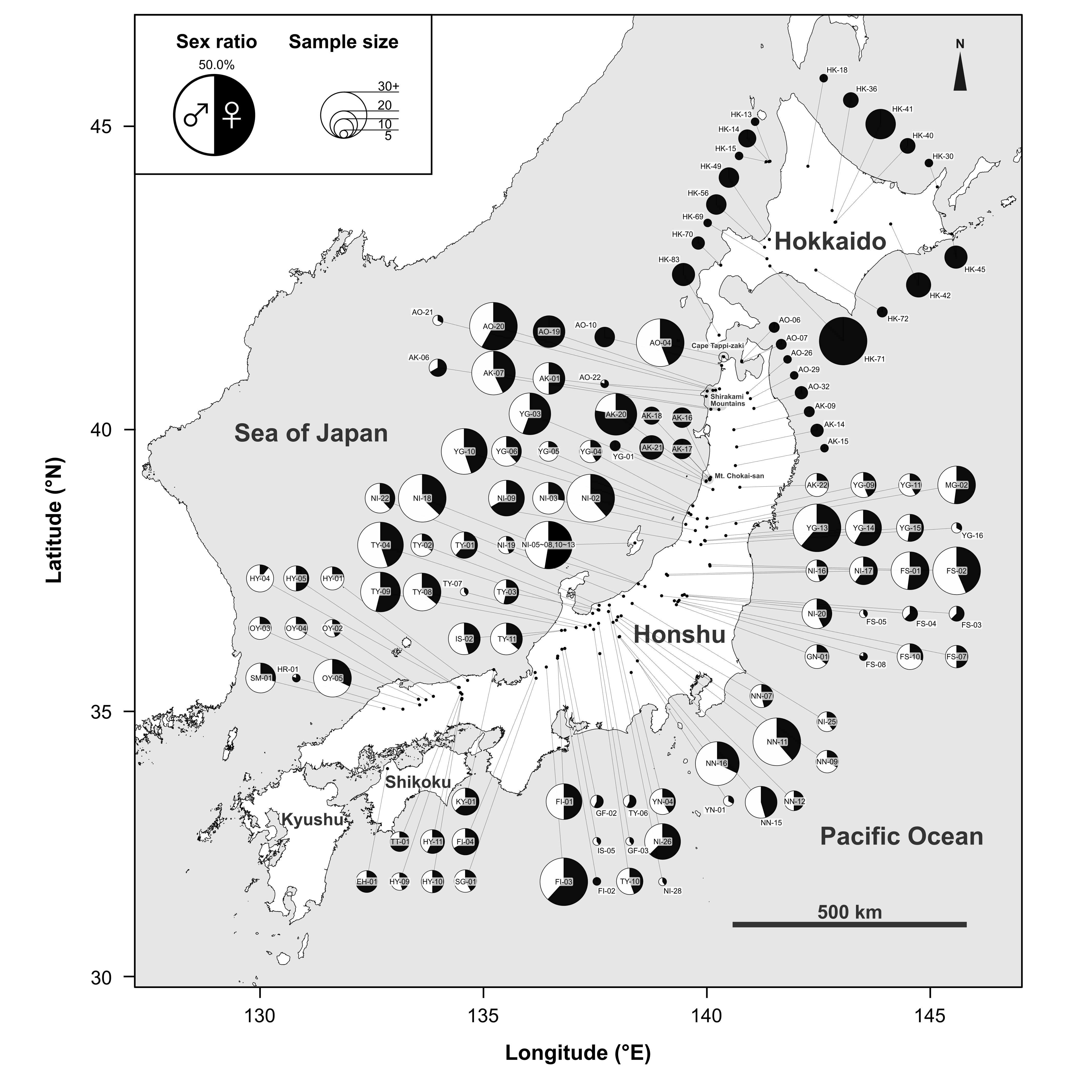
**Figure S1.** Geographic variation in sex ratio across Japanese populations of *Catapionus nebulosus* species group, with locality codes. Codes correspond to Table S1.

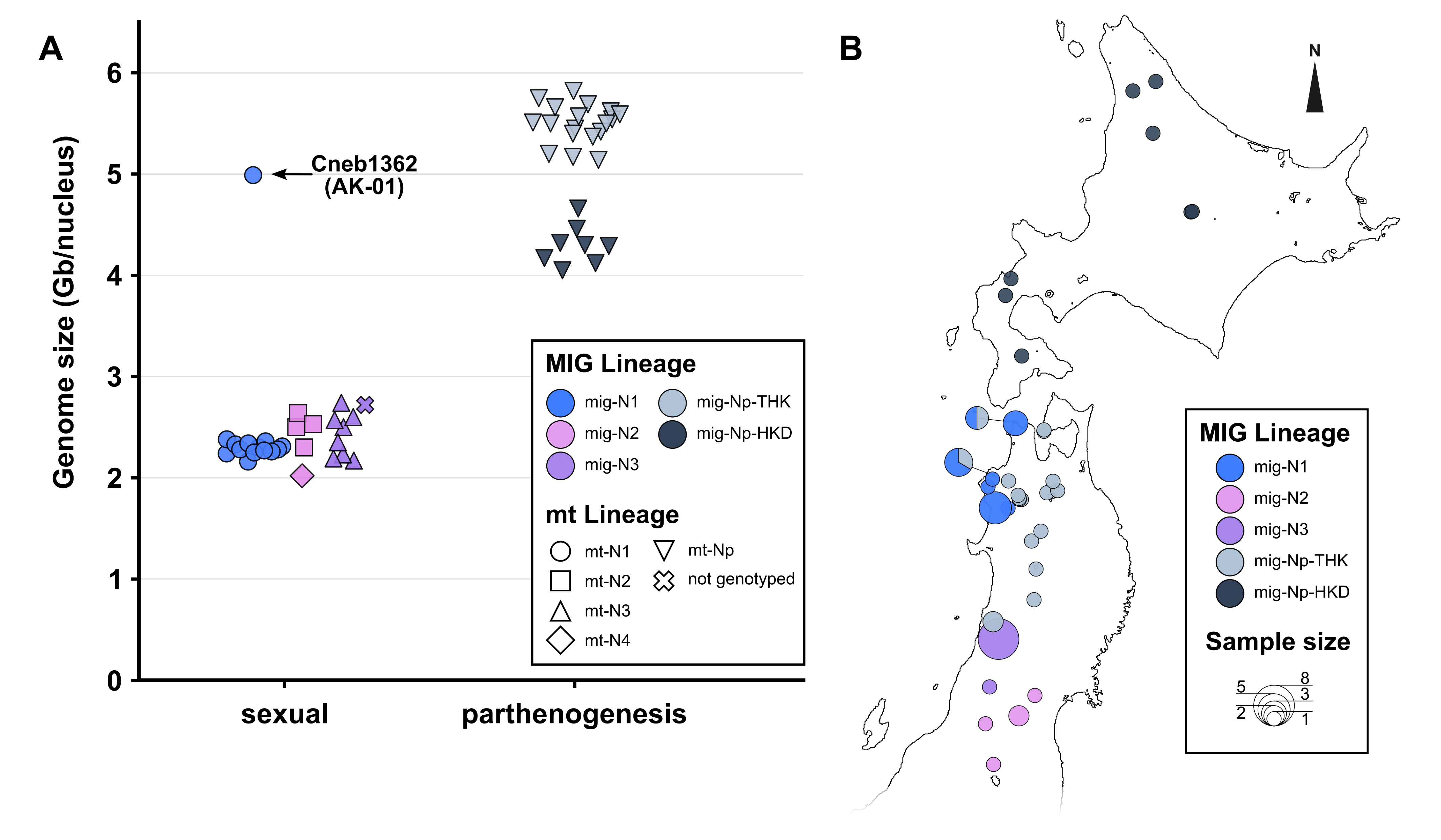
**Figure S2.** Genome size and its geographic distribution in the mig-N clade of the *Catapionus nebulosus* species group. (A) Nuclear DNA content per somatic nucleus, measured by flow cytometry. Colors indicate nuclear lineages. The three modes correspond to putative diploid (mig-N1, mig-N2, and mig-N3, excluding Cneb1362), tetraploid (mig-Np-HKD), and pentaploid (mig-Np-THK) classes. Cneb1362 (AK-01) is a tetraploid-like female recovered within the otherwise diploid lineage mig-N1 (see also Figures 4A and S8). Individual values are given in Table S2. (B) Localities of the measured individuals. Each pie chart shows the lineage composition, with its size proportional to the number of individuals.

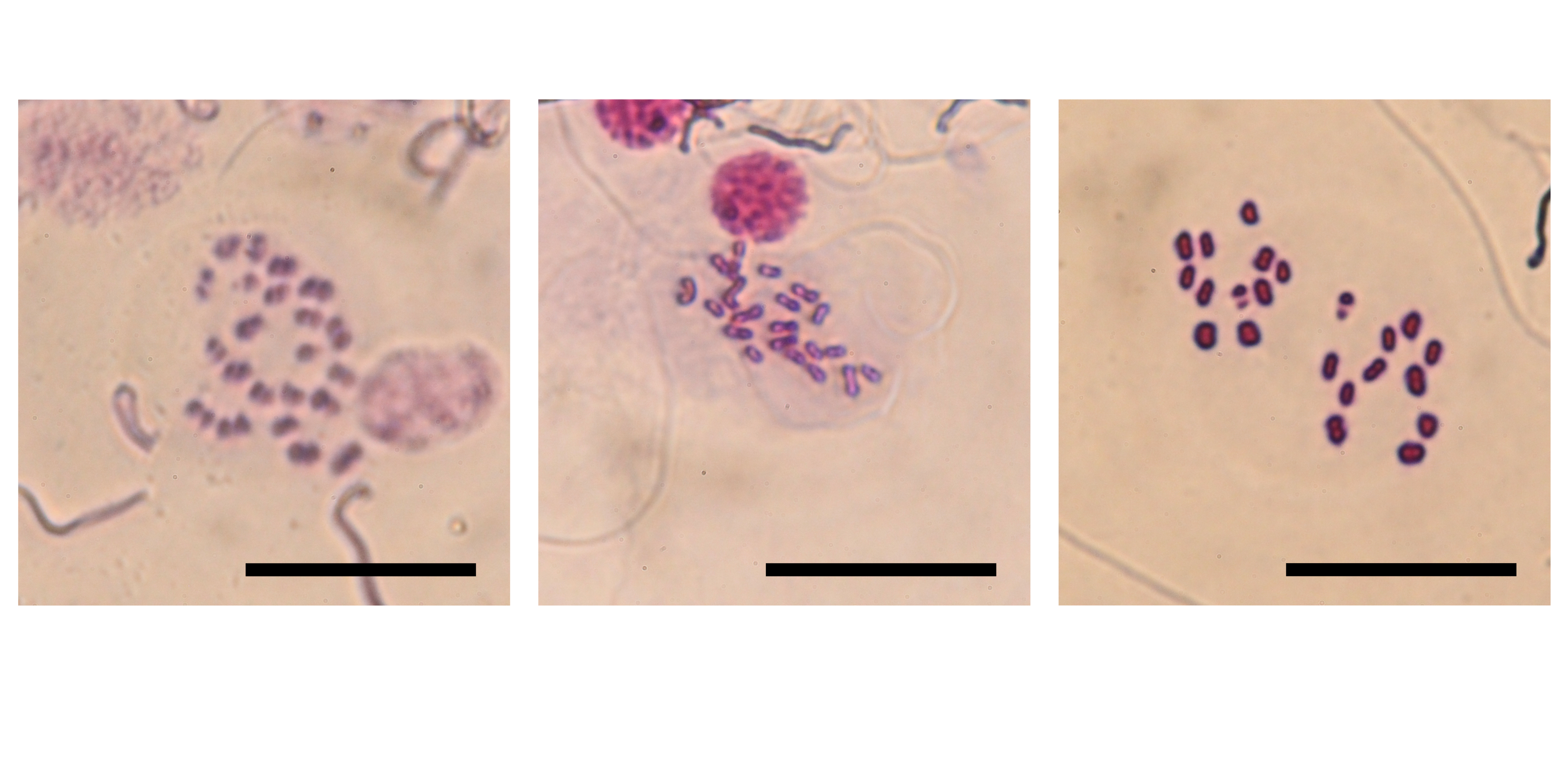
**Figure S3.** Mitotic spermatogonial chromosome spreads of three *C. modestus* individuals from the Shiga population, showing a diploid chromosome number of 2n = 22. Scale bars, 30 μm.

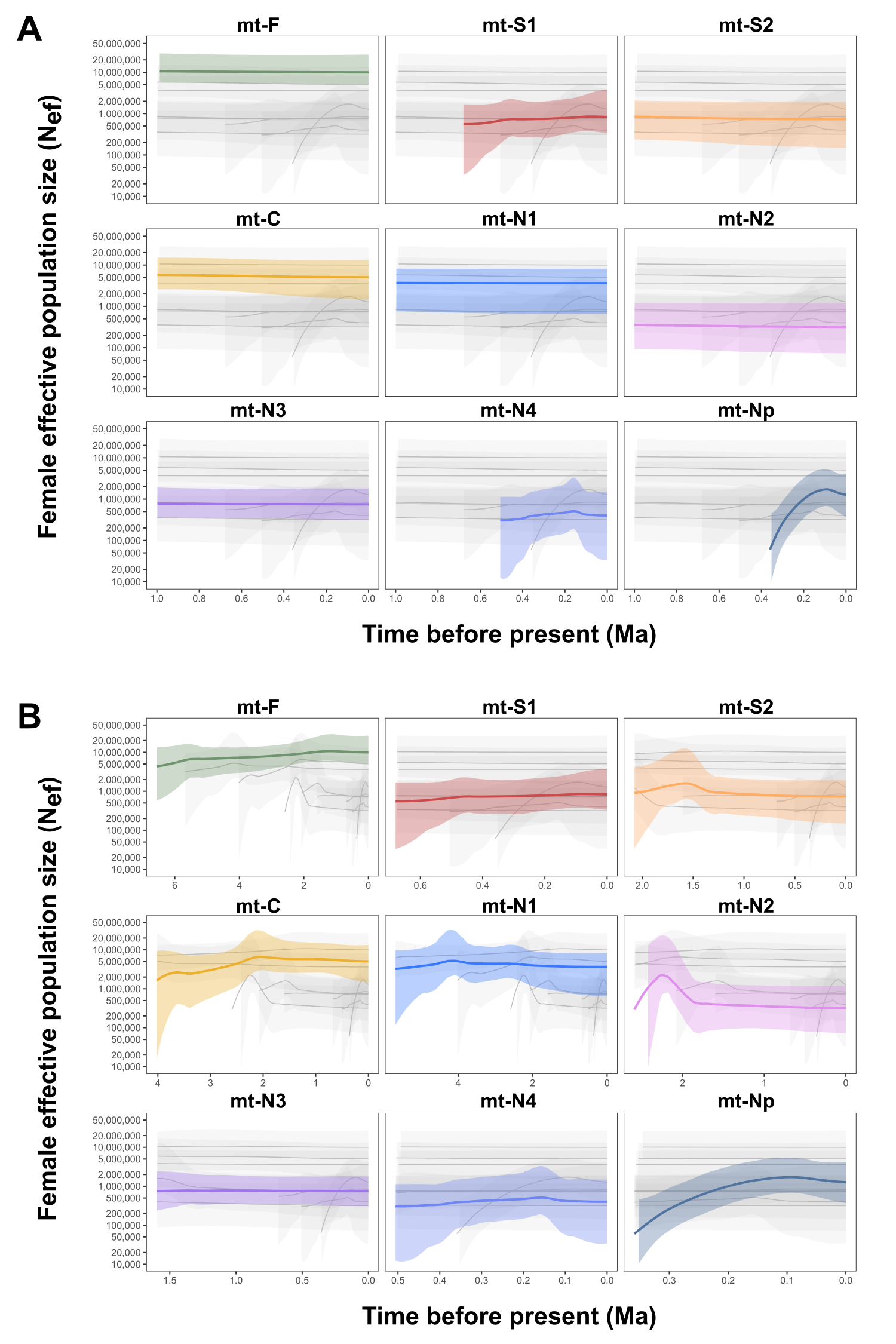
**Figure S4.** Bayesian Skyride reconstructions of female effective population size through time for each mitochondrial lineage of the *Catapionus nebulosus* species group, showing (A) over the past 1 million years ago, and (B) over the full time span of each lineage. In each panel the colored solid line indicates the median estimate for the focal lineage and the colored boundary indicates the 95% HPD interval. Gray lines and shading repeat the other lineages. Summaries are given in Table S6.

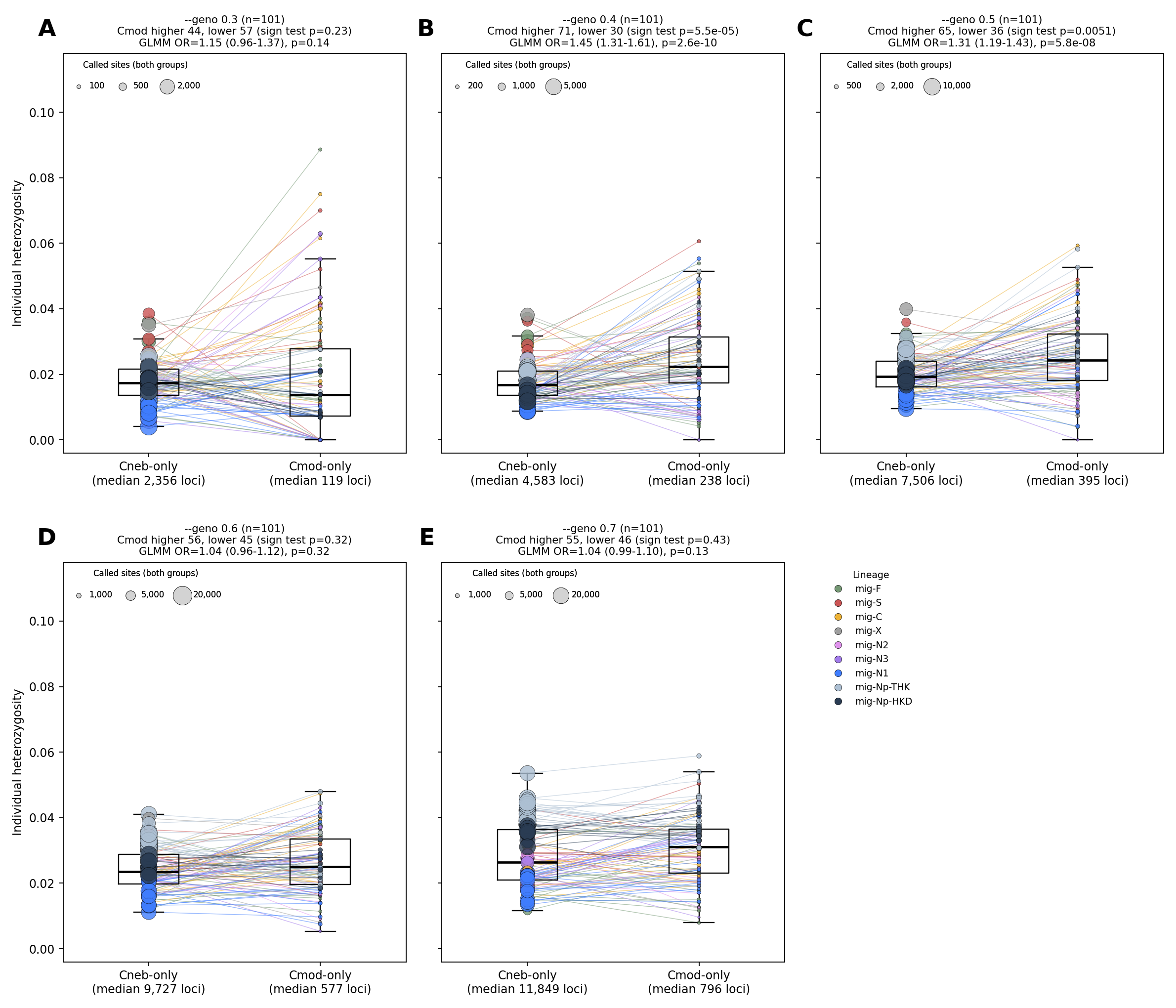

**Figure S5.** Individual heterozygosity at SNPs mapped to the *C. nebulosus* reference (Cneb-only) and to the *C. modestus* reference (Cmod-only) in the reference-flow procedure (minimum depth of three, --mind 0.97; n = 101). (A–E) --geno 0.3–0.7. Lines connect the two values of the same individual, and point area is proportional to the number of called sites. Boxes show the median and interquartile range, and whiskers extend to 1.5 times the interquartile range. Panel titles give the numbers of individuals with higher and lower heterozygosity at Cmod-only SNPs (sign test) and the odds ratio (Cmod-only / Cneb-only) from a binomial GLMM with individual as a random effect, fitted by penalized quasi-likelihood.

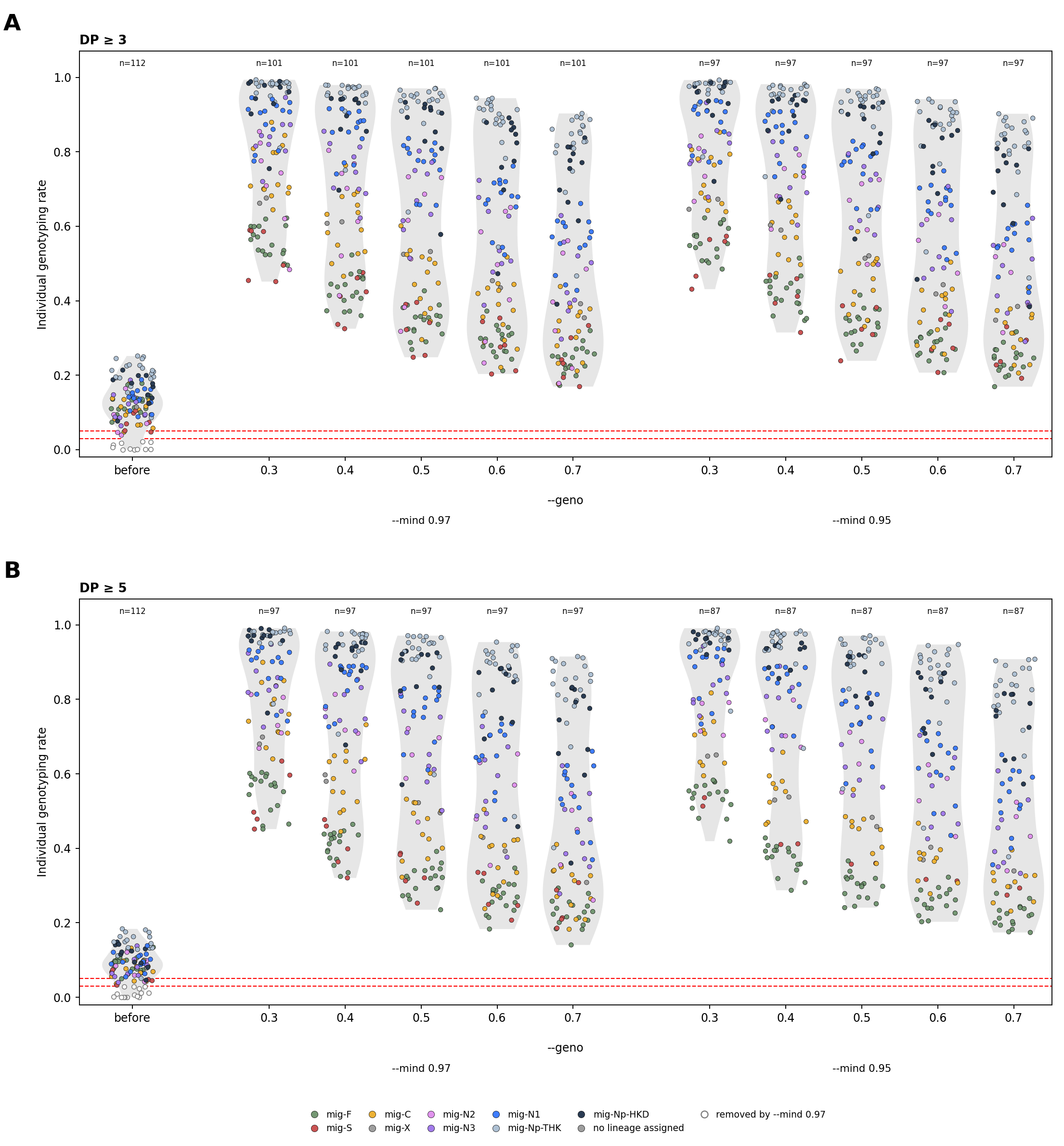

**Figure S6.** Individual genotyping rate before and after filtering. (A) Minimum depth of three. (B) Minimum depth of five. In each panel, the leftmost column shows the genotyping rate of each individual at all SNP sites before filtering. The other columns show it after site filtering at each --geno threshold, for the individuals retained with --mind 0.97 (left) or --mind 0.95 (right). The dashed lines indicate the individual thresholds of 3% (--mind 0.97) and 5% (--mind 0.95). Open circles indicate individuals removed by --mind 0.97. Colors indicate nuclear lineages. Violins show the distribution across individuals, and the number of individuals is given above each column. The primary dataset has a minimum depth of three, --mind 0.97, and --geno 0.5.

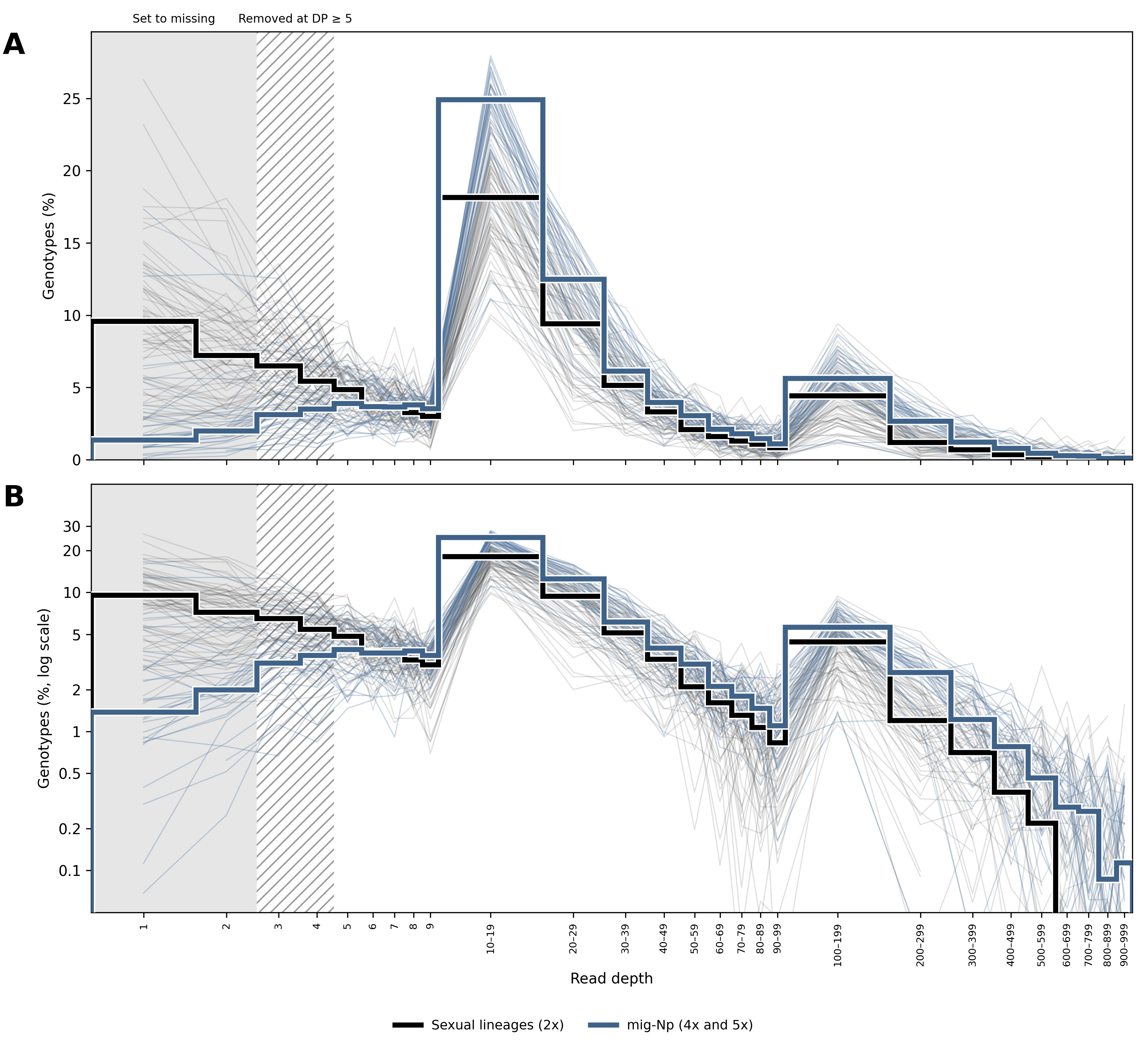
**Figure S7.** Read depth of genotypes in the primary dataset (minimum depth of three, --mind 0.97, and --geno 0.5; 11,940 SNPs before LD pruning; n = 101). (A) Linear scale. (B) Logarithmic scale. Thin lines represent individuals of the sexual lineages (gray) and of mig-Np (blue), and thick lines show the median across individuals of each group. Values are percentages of all genotypes called by Stacks at these SNPs in each individual. Bins are single depths up to 9, intervals of 10 from 10 to 99, and intervals of 100 from 100 to 999, so values are not comparable across these ranges. Genotypes with a depth of one or two (gray shading) were set to missing. Genotypes with a depth of three or four (hatched) were used in the main analyses and were removed at a minimum depth of five. Genotypes with a depth of 1,000 or more (0.36% of all genotypes) are not shown.

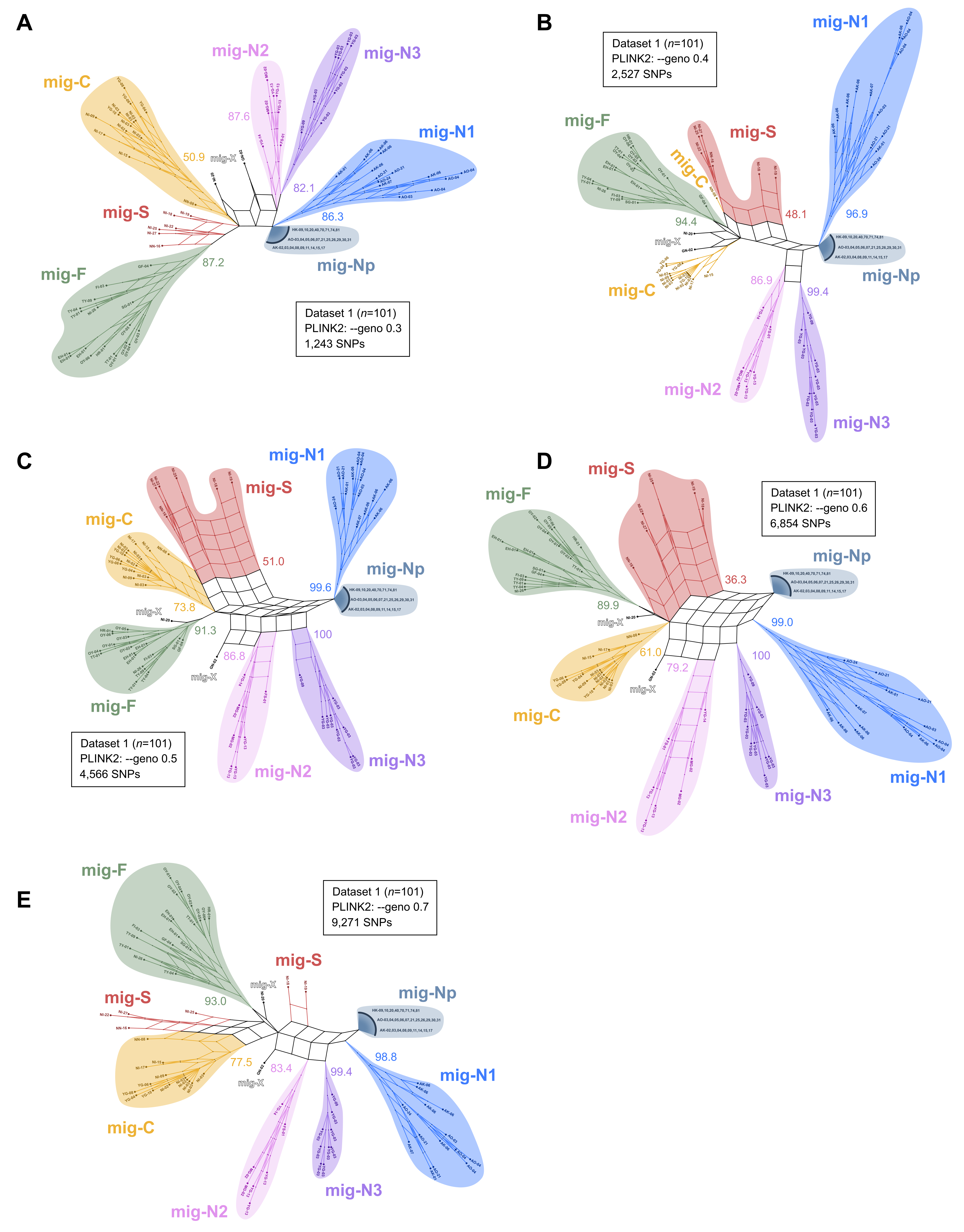
**Figure S8.** Neighbor-Net split networks of the 101 individuals of the *Catapionus nebulosus* species group at five --geno thresholds: (A) --geno 0.3, (B) 0.4, (C) 0.5, (D) 0.6, and (E) 0.7. SNP counts and bootstrap support values are given in **Table S12**.

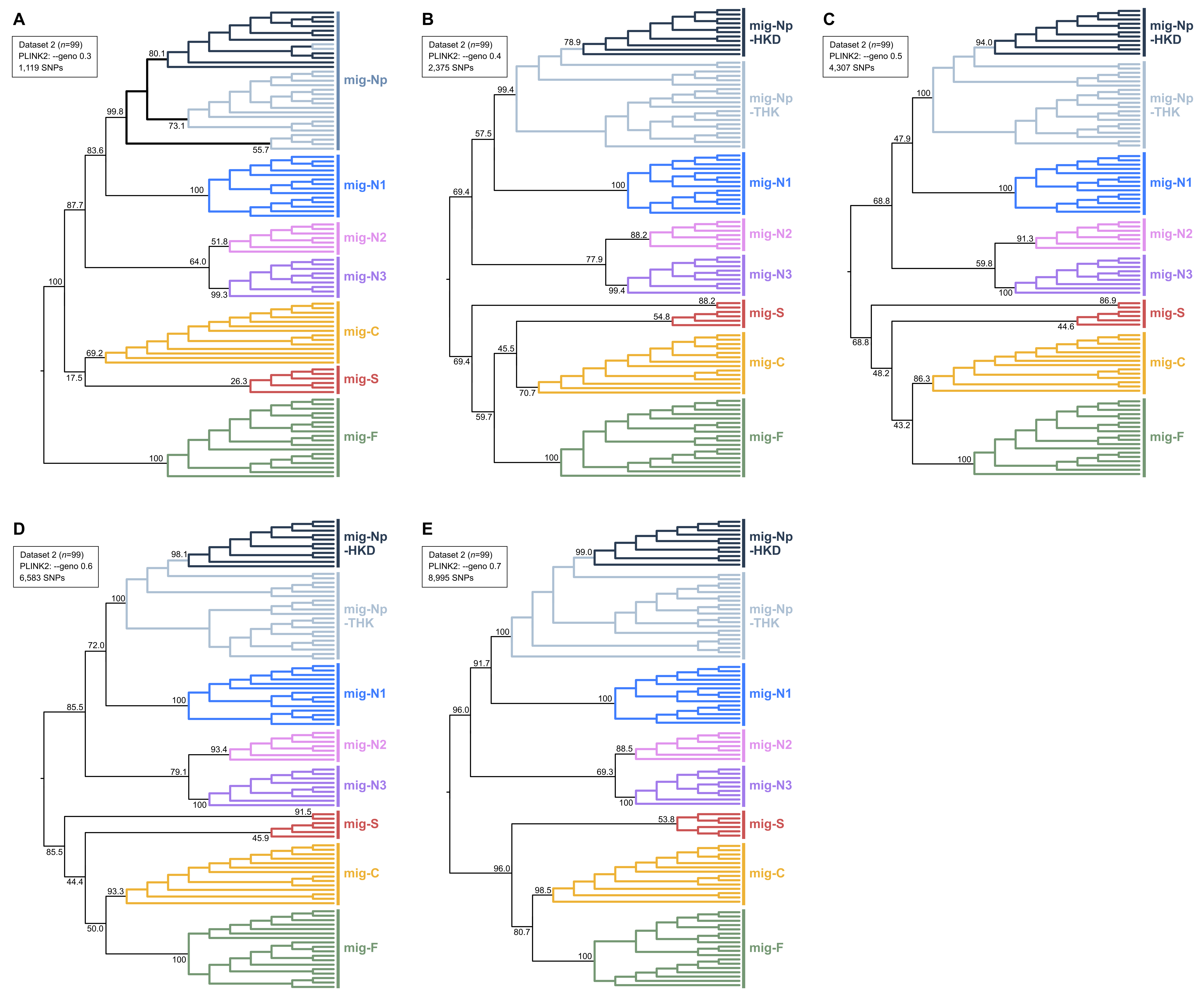
**Figure S9.** SVDquartets species trees of the 99 individuals of the *Catapionus nebulosus* species group at five --geno thresholds: (A) --geno 0.3, (B) 0.4, (C) 0.5, (D) 0.6, and (E) 0.7. SNP counts and bootstrap support values are given in **Table S13**.

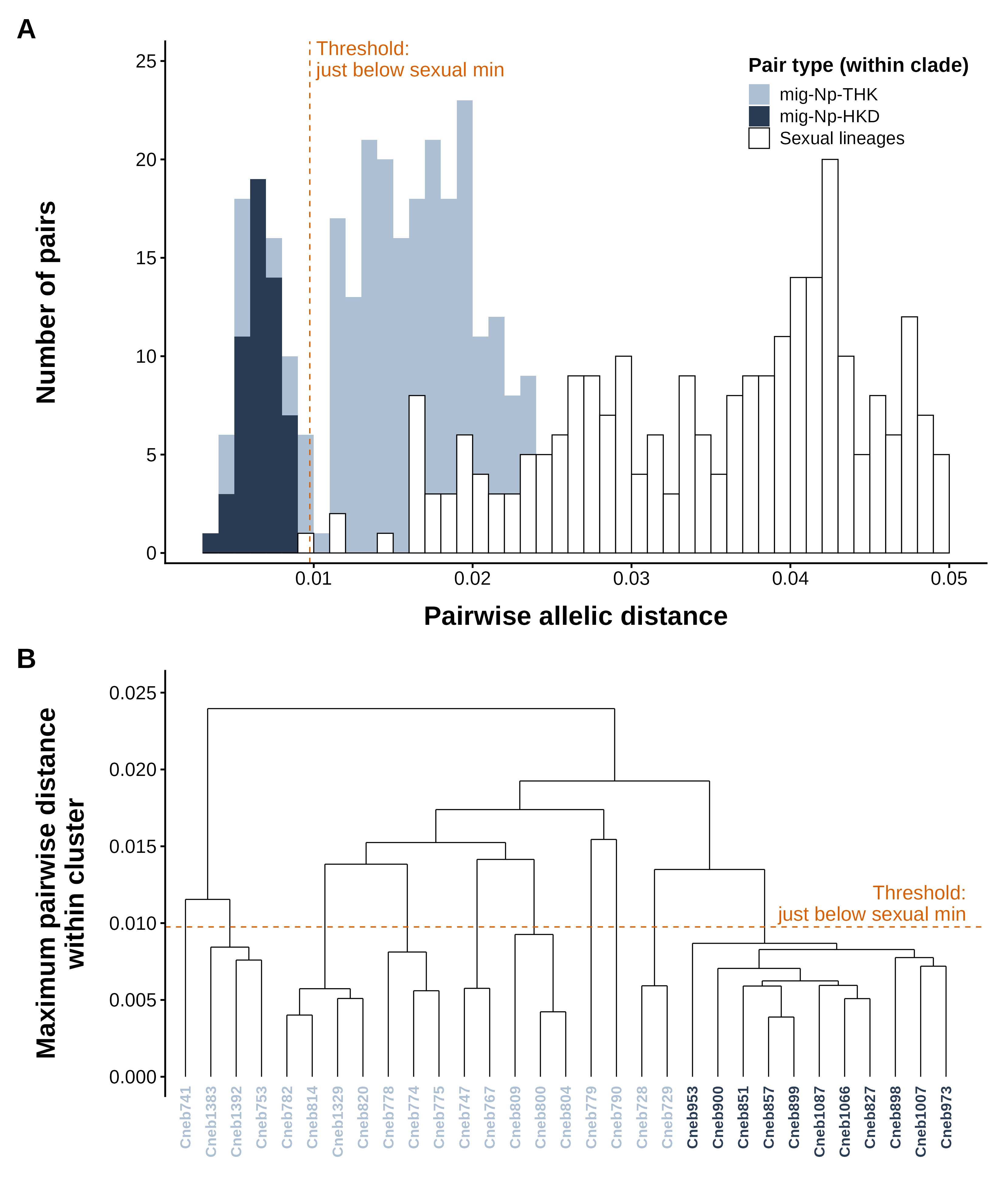
**Figure S10.** Definition of multilocus lineages (MLLs) within mig-Np (minimum depth of three, --mind 0.97, and --geno 0.5). (A) Distribution of pairwise allelic distances up to 0.05 within mig-Np-THK, within mig-Np-HKD, and between individuals of the sexual lineages. Dashed lines show the candidate thresholds 0.0047, 0.0098 (adopted), and 0.0125. The smallest distance between two sexual individuals was 0.0099, above the adopted threshold. (B) Farthest-neighbor (complete-linkage) dendrogram of the 31 mig-Np individuals, with dashed lines as in panel A. Tip labels are colored as in panel A. Cutting at 0.0098 gave 10 MLLs.

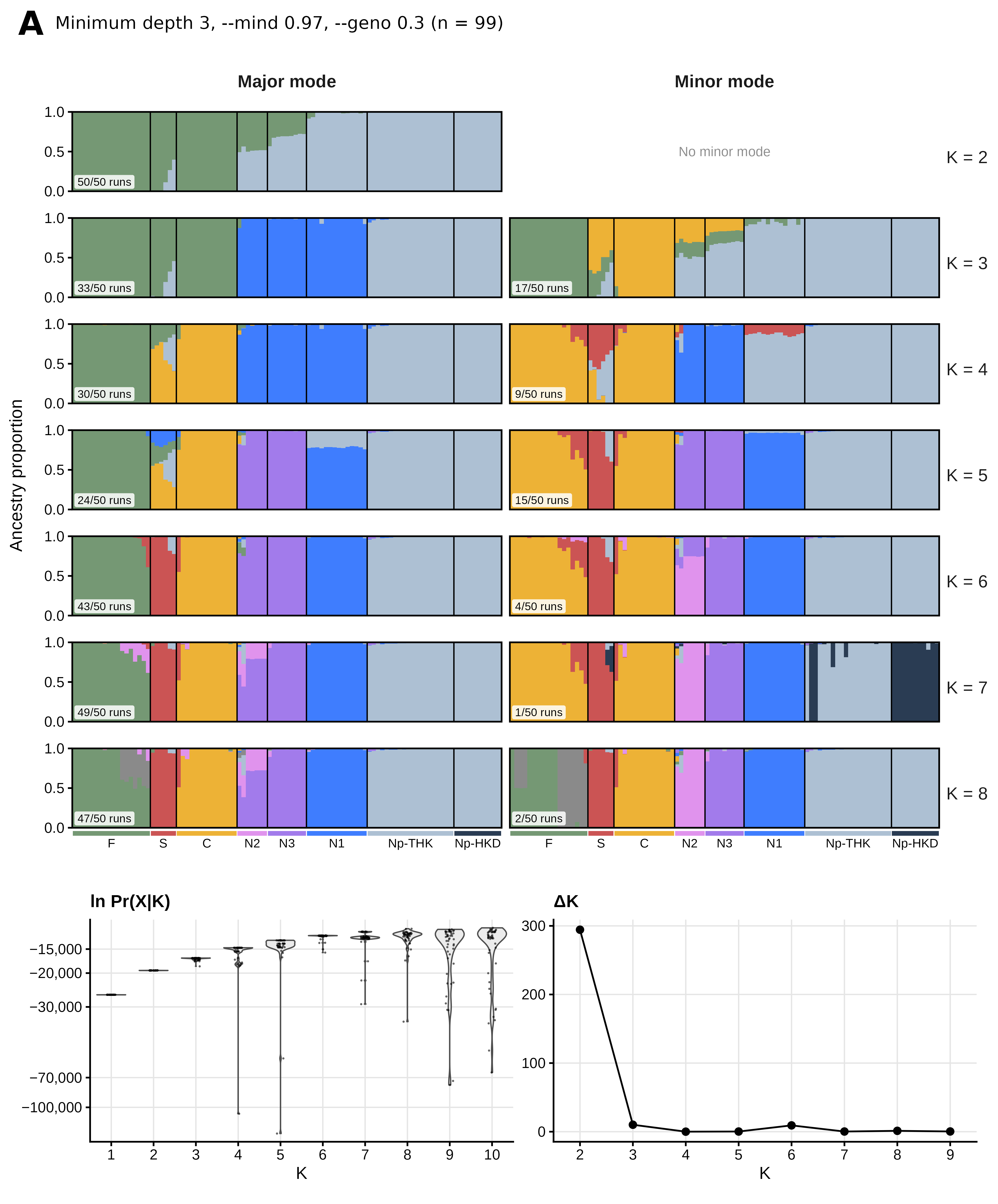
**Figure S11**. STRUCTURE results for all filtering conditions. Each page shows one dataset: (A–E) minimum depth of three and --mind 0.97 at --geno 0.3, 0.4, 0.5, 0.6, and 0.7 (C, primary dataset); (F) clone-corrected --geno 0.5 dataset, which retained one individual per multilocus lineage; (G–I) --geno 0.5 datasets with a minimum depth of three and --mind 0.95 (G), a minimum depth of five and --mind 0.97 (H), and a minimum depth of five and --mind 0.95 (I). For each K, replicate runs were grouped into modes (G′ ≥ 0.9). The major mode and the second most frequent mode are shown for K = 2–8, with the number of runs in each mode out of 50. The lower panels show the distribution of ln Pr(X|K) across the 50 runs on a logarithmic scale for K = 1–10, and ΔK for K = 2–9. Colors follow Figure 5. Lineage labels omit the prefix “mig-”.

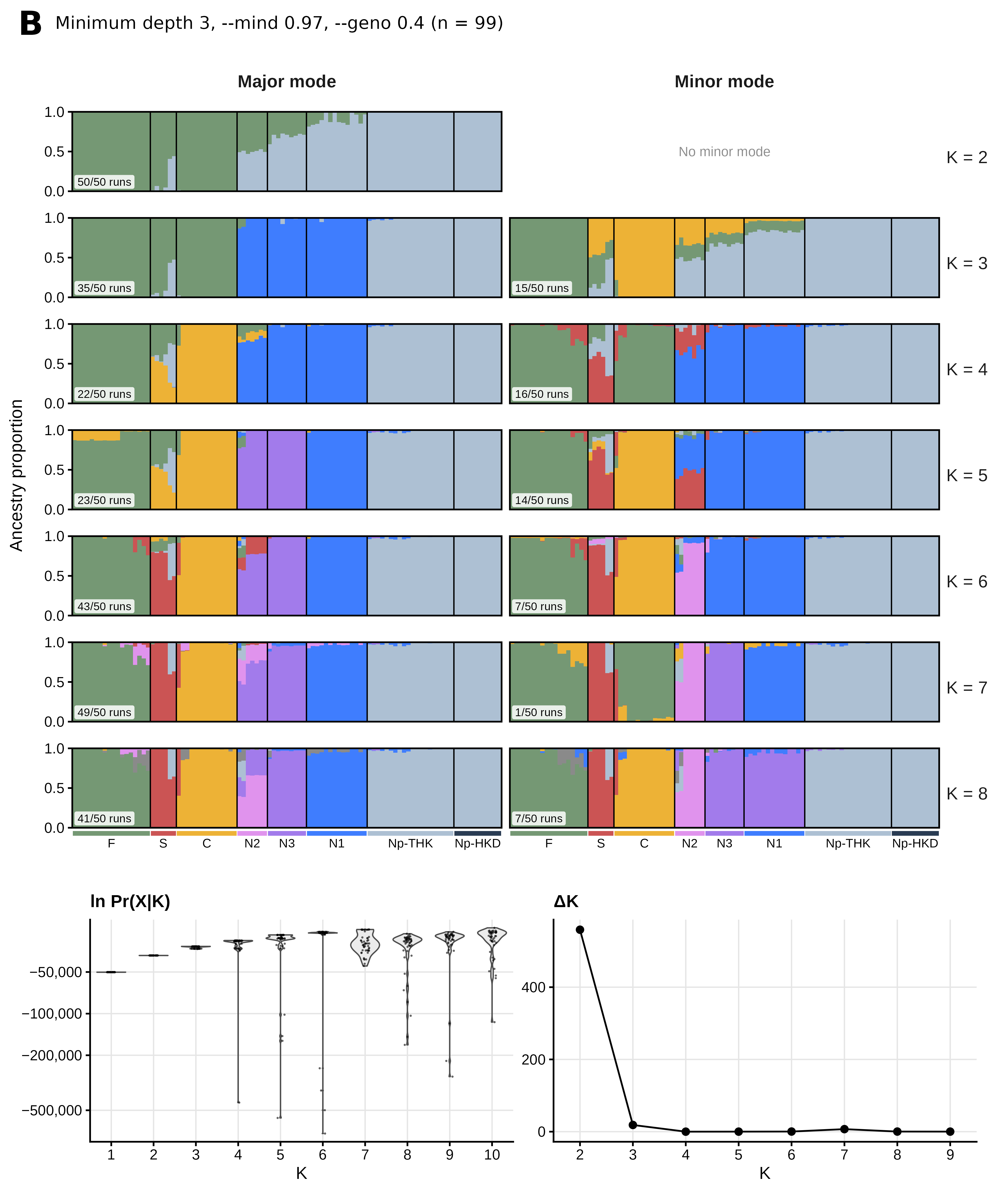

**Figure S11 (continued)**.

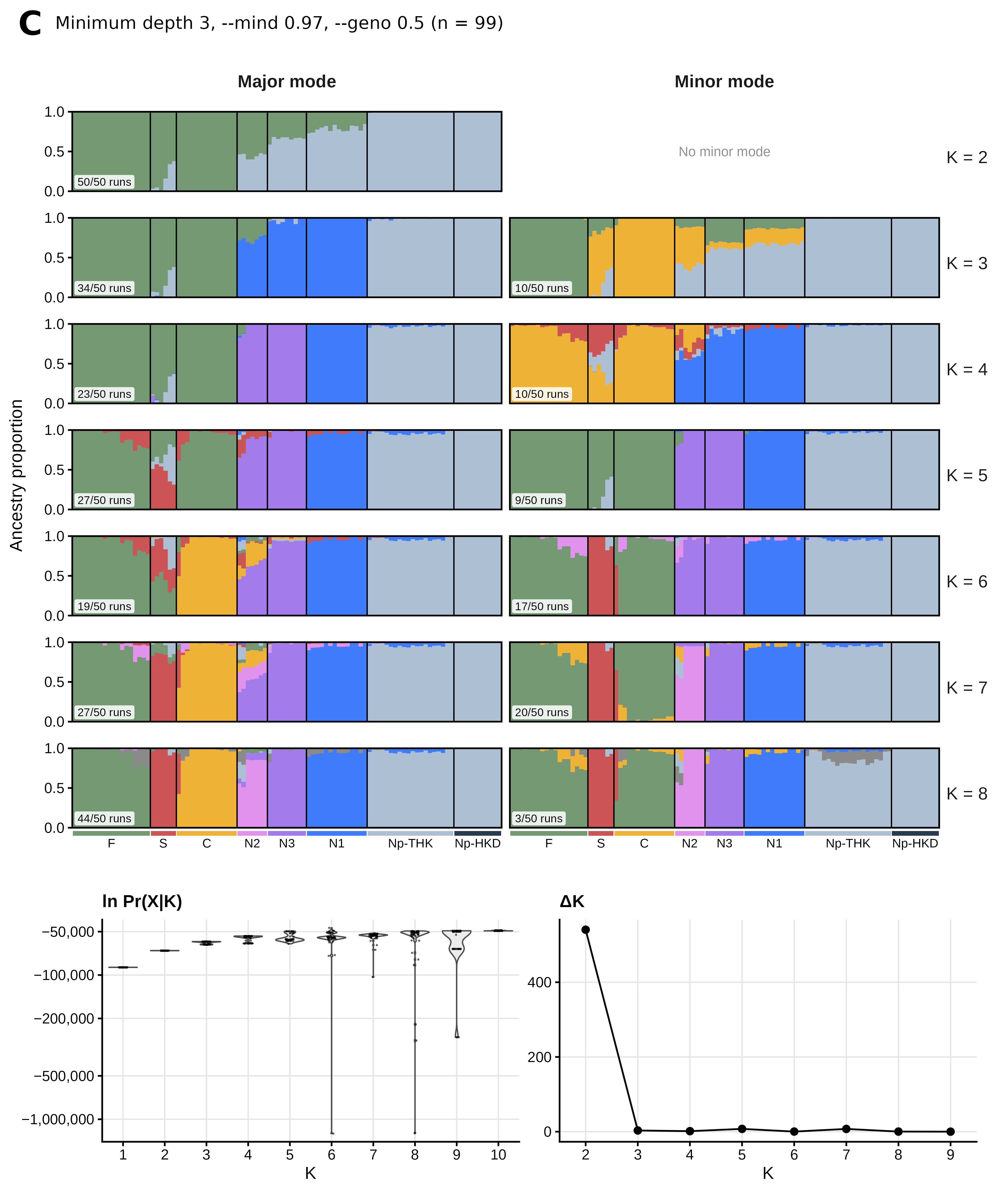

**Figure S11 (continued)**.

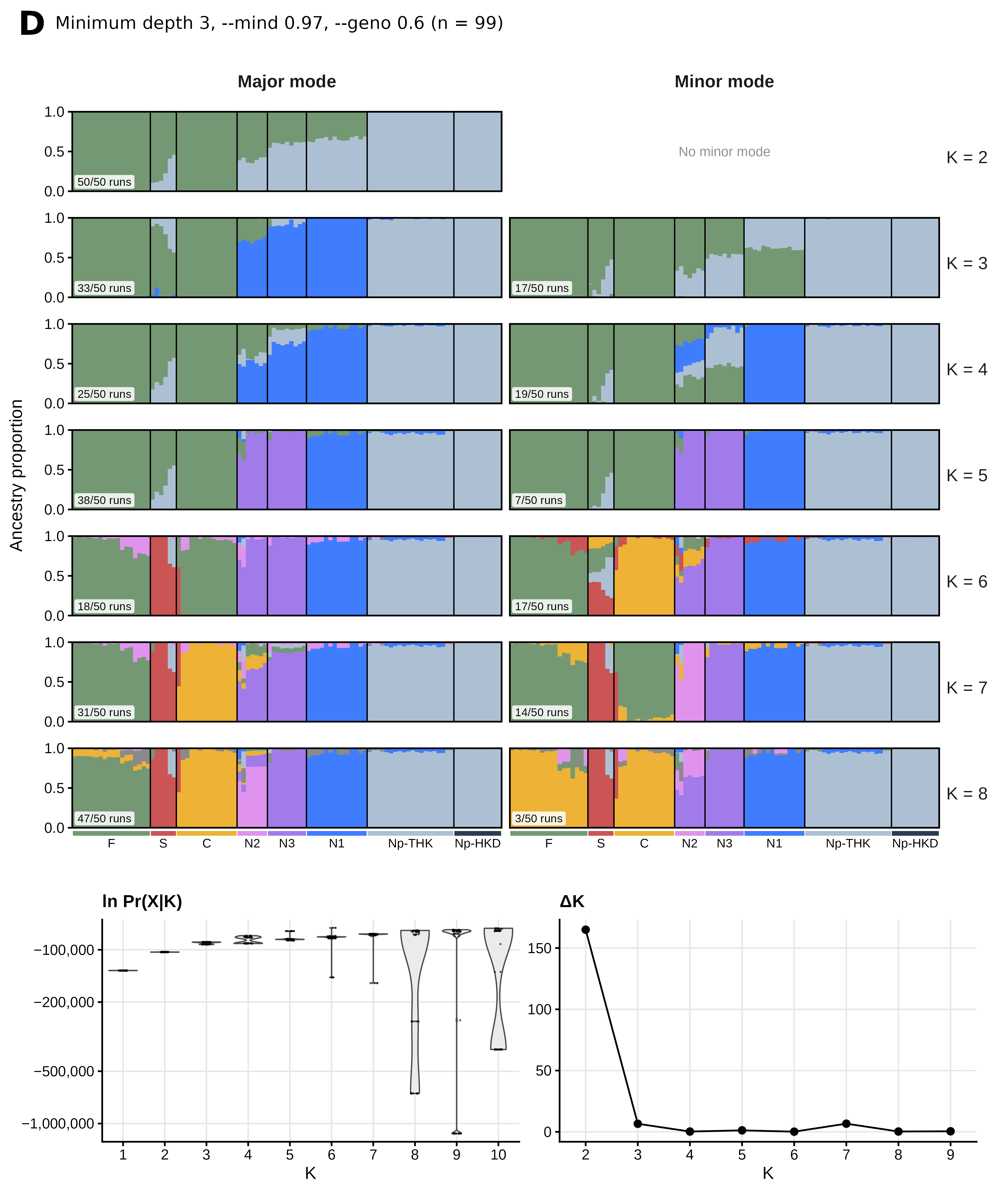

**Figure S11 (continued)**.

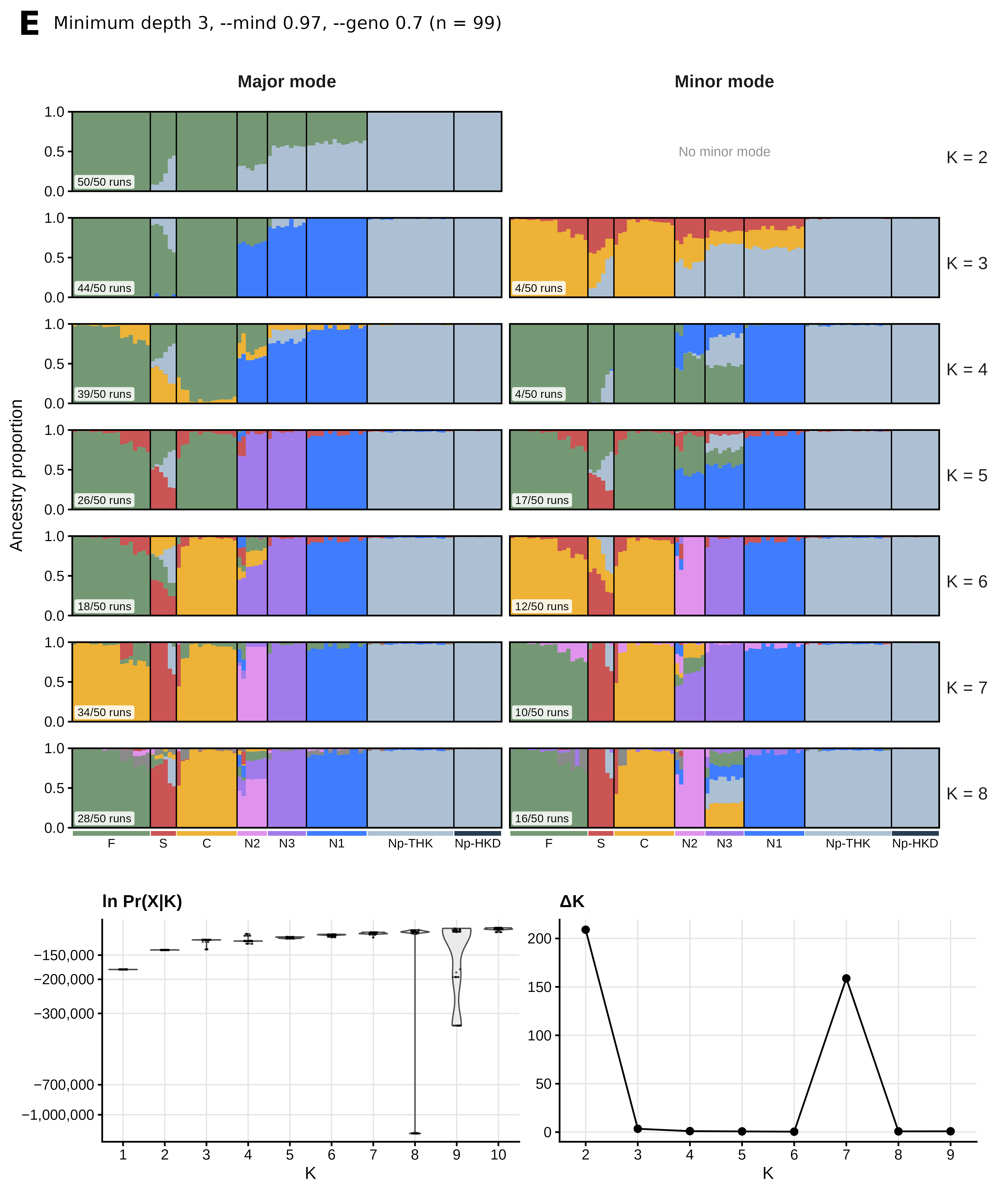
**Figure S11 (continued)**.

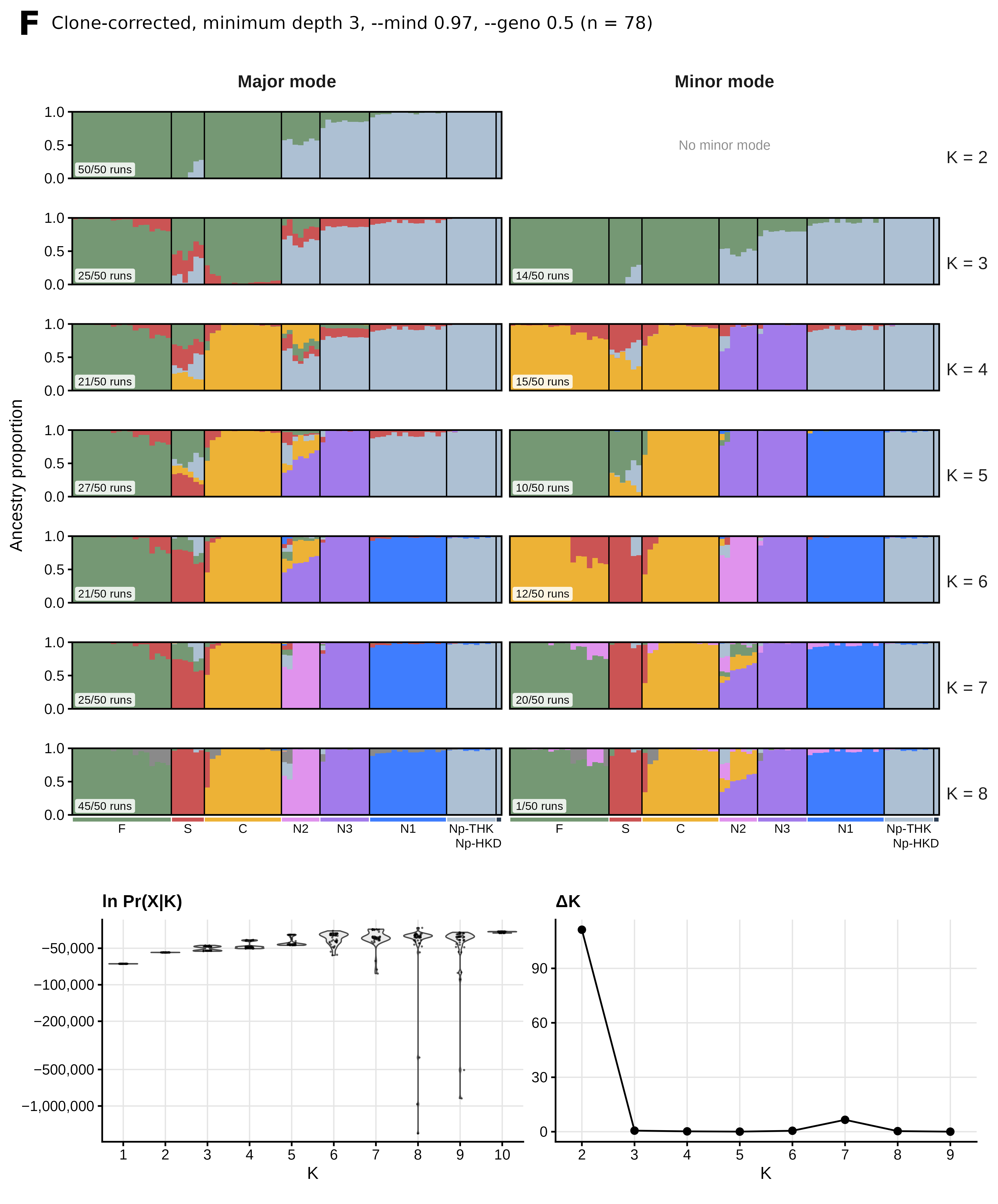

**Figure S11 (continued)**.

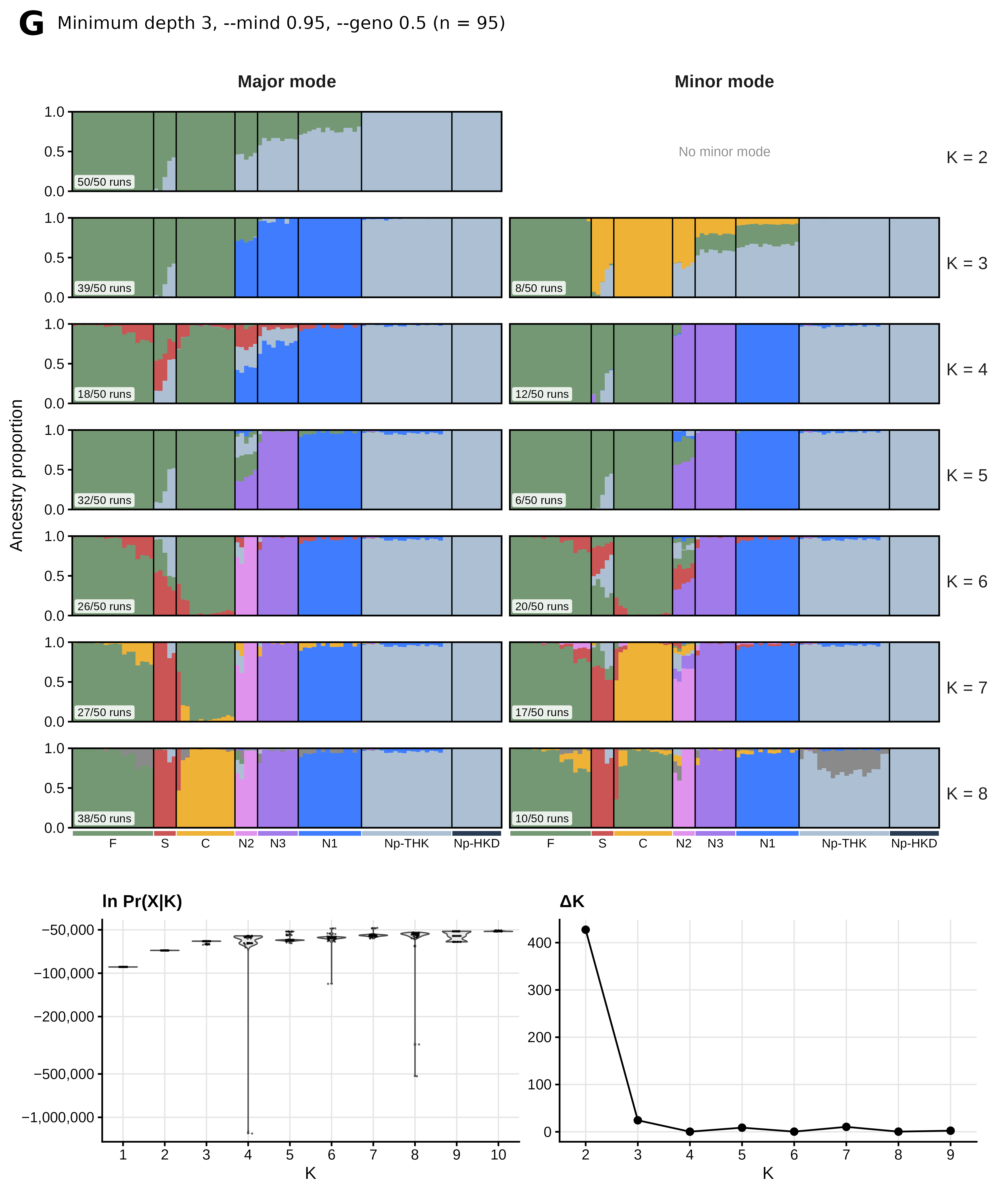
**Figure S11 (continued)**.

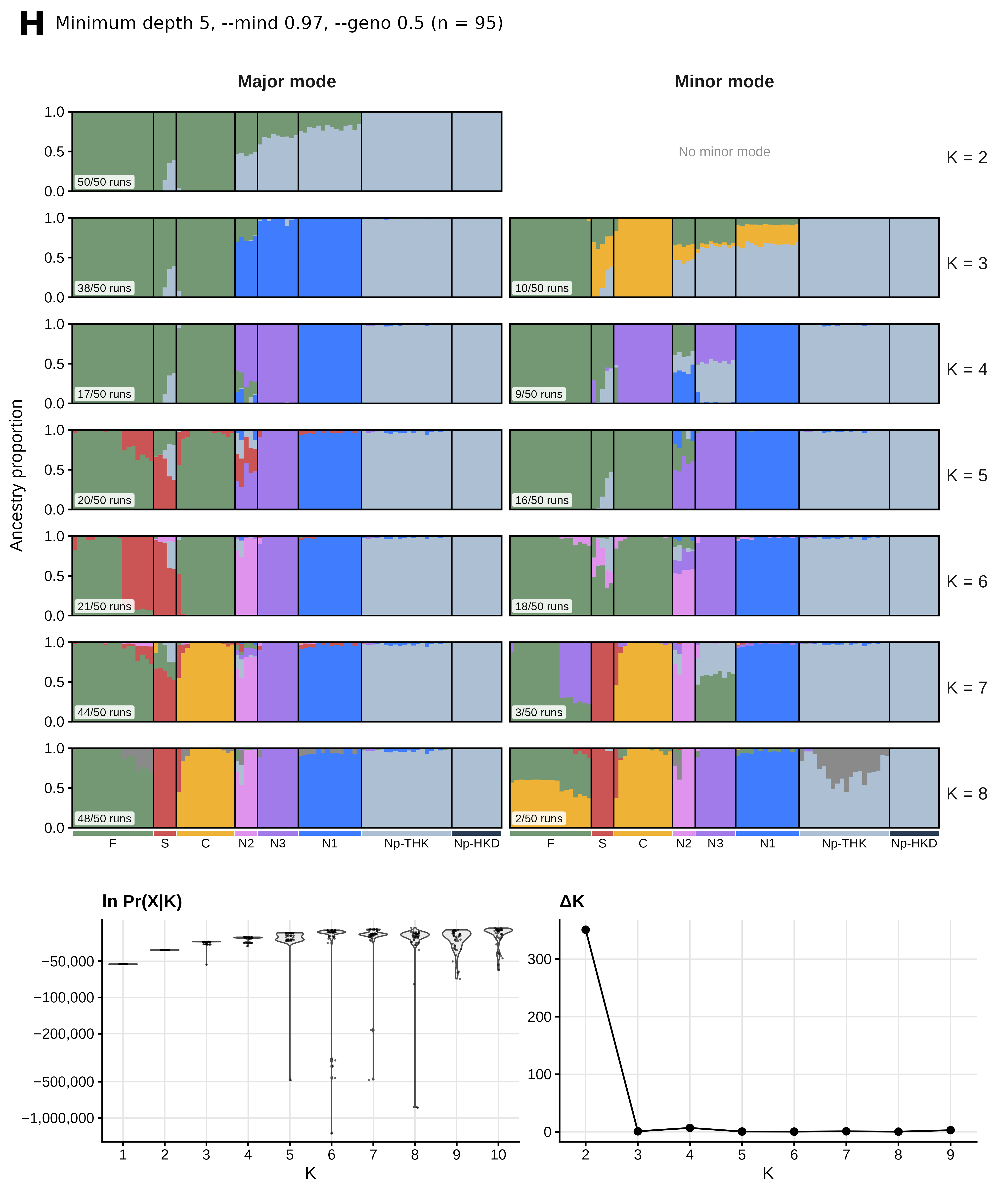

**Figure S11 (continued)**.

**Figure S11 (continued)**.

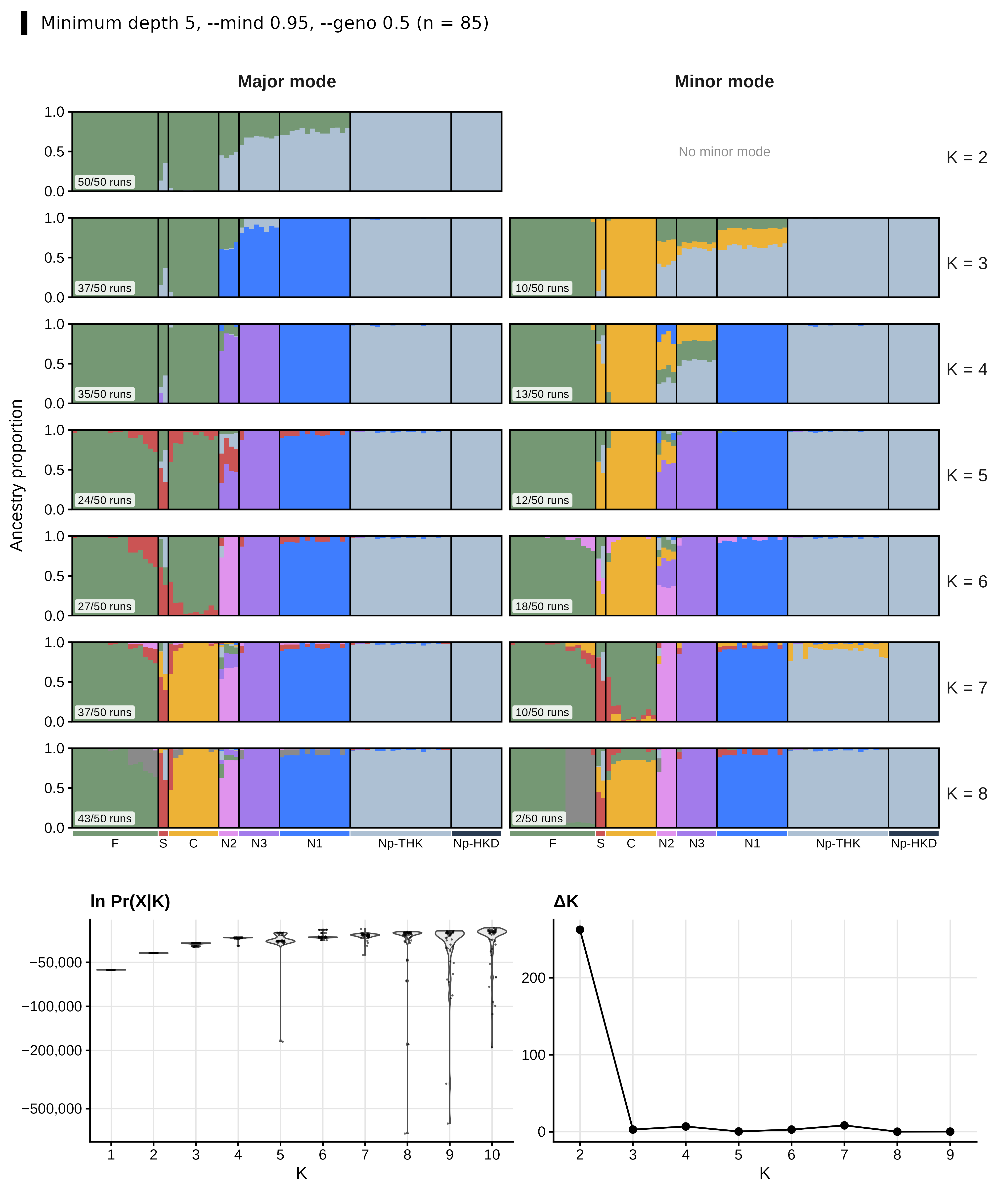

**Figure S12.** Number of genetic clusters estimated with the Puechmaille estimators from the STRUCTURE results. Sampling localities were used as groups. For each K, a cluster was counted when its median (MedMedK and
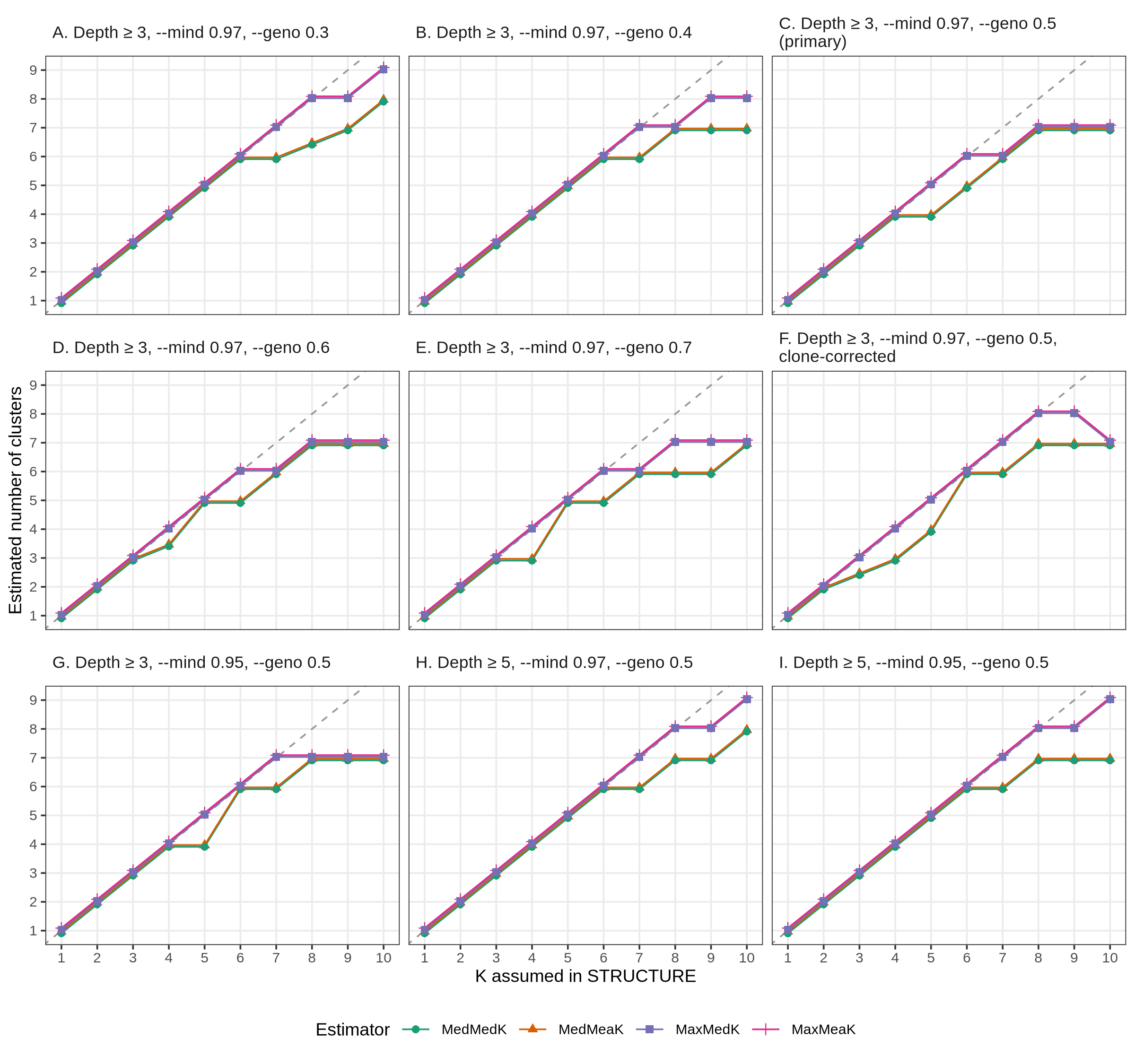
MaxMedK) or mean (MedMeaK and MaxMeaK) membership coefficient exceeded 0.5 in at least one locality. MedMedK and MedMeaK are medians, and MaxMedK and MaxMeaK are maxima, across the 50 replicate runs for each K; medians can take half-integer values. The dashed line indicates equality between the estimated and assumed K. (A–E) Datasets with a minimum depth of three and --mind 0.97 at --geno 0.3, 0.4, 0.5, 0.6, and 0.7. Panel C is the primary dataset. (F) Clone-corrected --geno 0.5 dataset, which retained one individual per multilocus lineage. (G–I) --geno 0.5 datasets with a minimum depth of three and --mind 0.95 (G), a minimum depth of five and --mind 0.97 (H), and a minimum depth of five and --mind 0.95 (I). Points are offset slightly so that overlapping estimators remain visible.

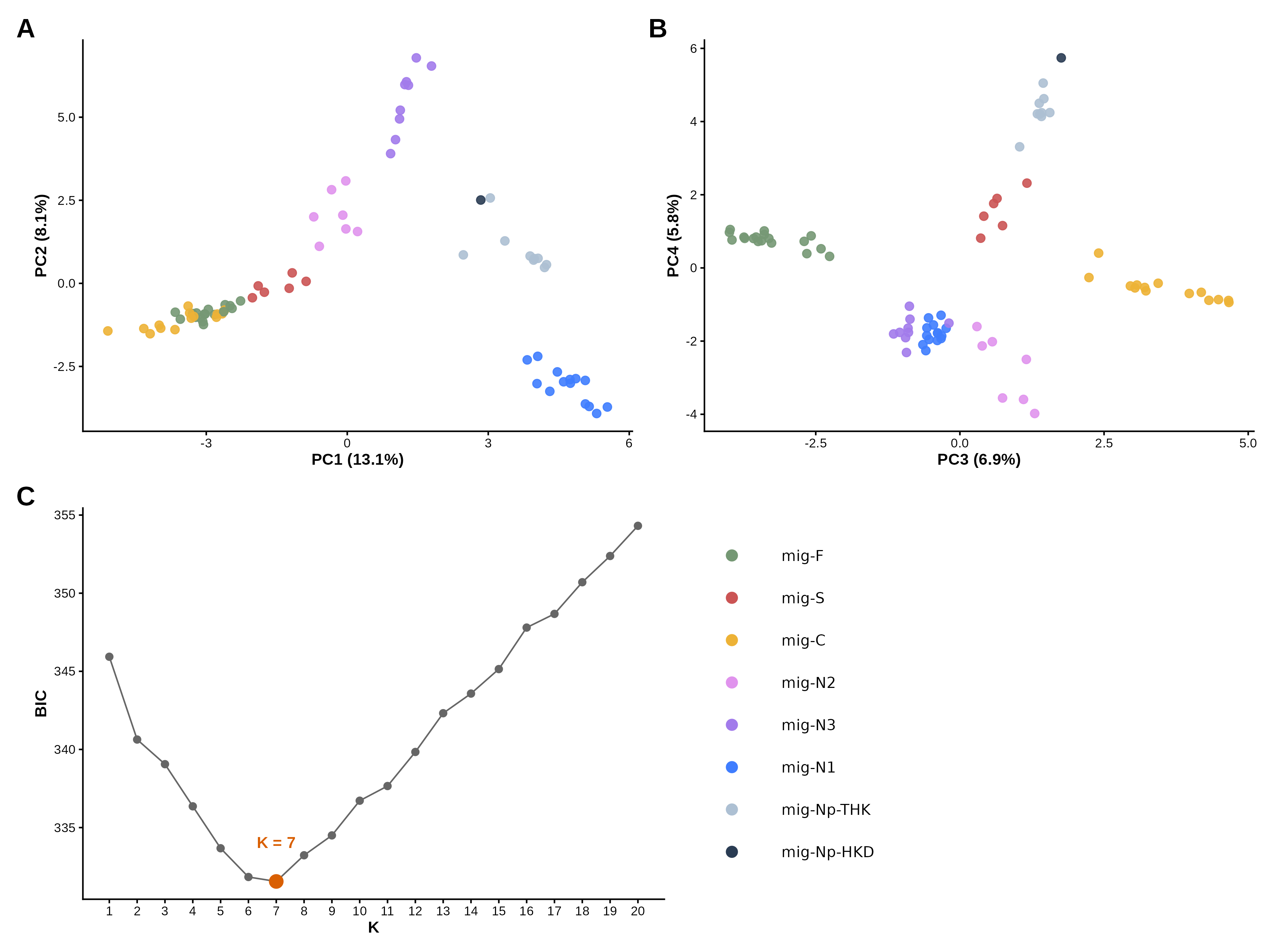

**Figure S13.** Principal component analysis (PCA) and k-means clustering of the clone-corrected --geno 0.5 dataset (n = 78; 4,307 LD-pruned SNPs). (A) PC1 and PC2. (B) PC3 and PC4. Colors indicate nuclear lineages, and percentages give the proportion of variance explained by each principal component. The open circle indicates the individual of mig-C assigned to the cluster of mig-S. (C) Bayesian information criterion (BIC) of k-means clustering on all principal components for K = 1–20. K = 7 was the smallest K after which the BIC increased. The seven k-means clusters matched the seven groups except for the individual marked in A and B.

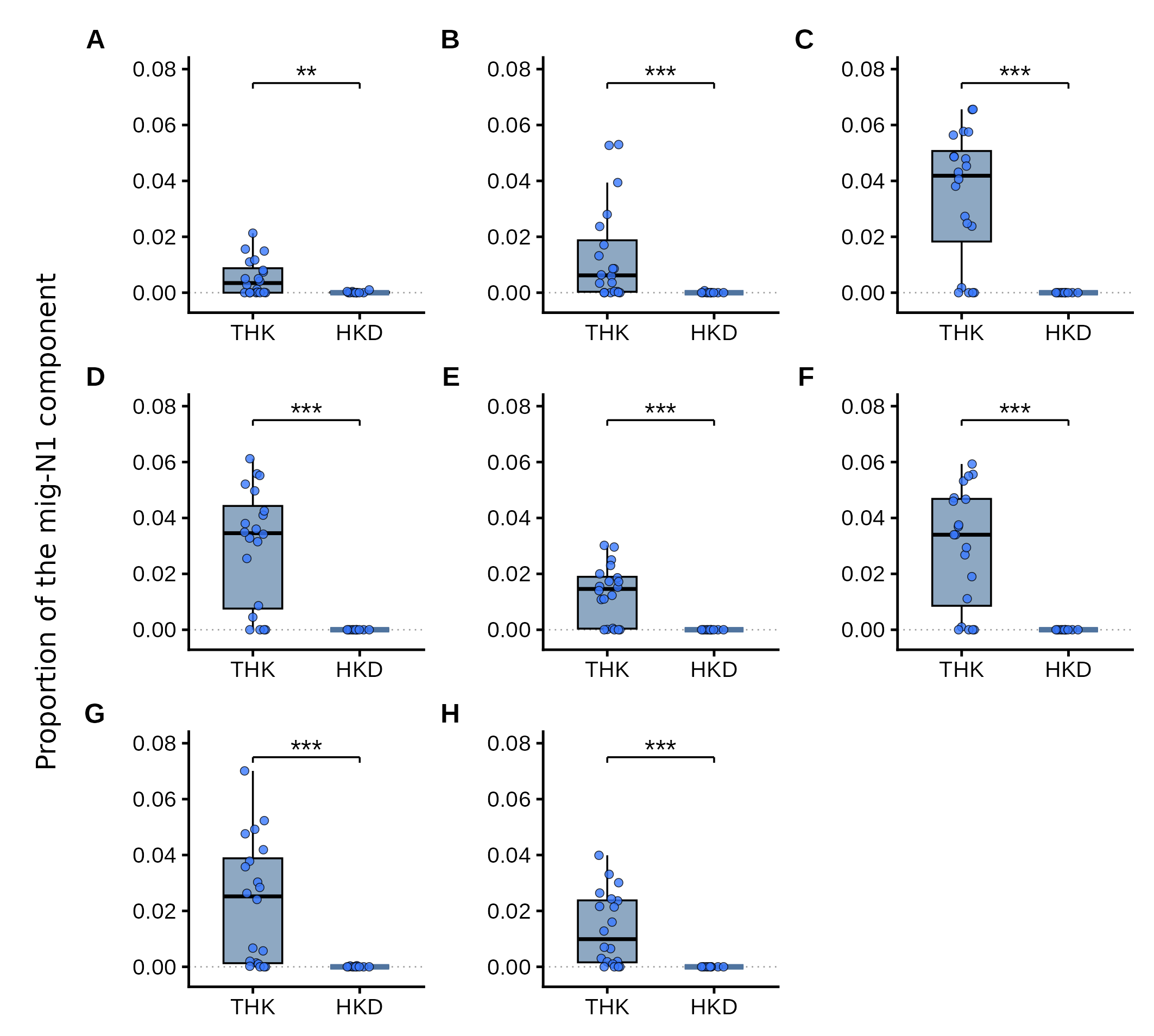

**Figure S14.** Proportion of the STRUCTURE component predominant in mig-N1 in individuals of mig-Np at K = 8 (major mode). Each point represents one individual, and boxes show the median and interquartile range for mig-Np-THK (THK; n = 20) and mig-Np-HKD (HKD; n = 11; n = 10 in panel H). The dotted line indicates zero. Asterisks indicate significant differences between THK and HKD (Wilcoxon rank-sum test; **P < 0.01, ***P < 0.001). P values are uncorrected; all remained below 0.01 after Holm correction. (A–E) Datasets with a minimum depth of three and --mind 0.97 at --geno 0.3 (A), 0.4 (B), 0.5 (C, primary dataset), 0.6 (D), and 0.7 (E). (F–H) --geno 0.5 datasets with a minimum depth of three and --mind 0.95 (F), a minimum depth of five and --mind 0.97 (G), and a minimum depth of five and --mind 0.95 (H). The clone-corrected dataset is not shown because it retained only one individual of mig-Np-HKD. Values are given in Table S15.

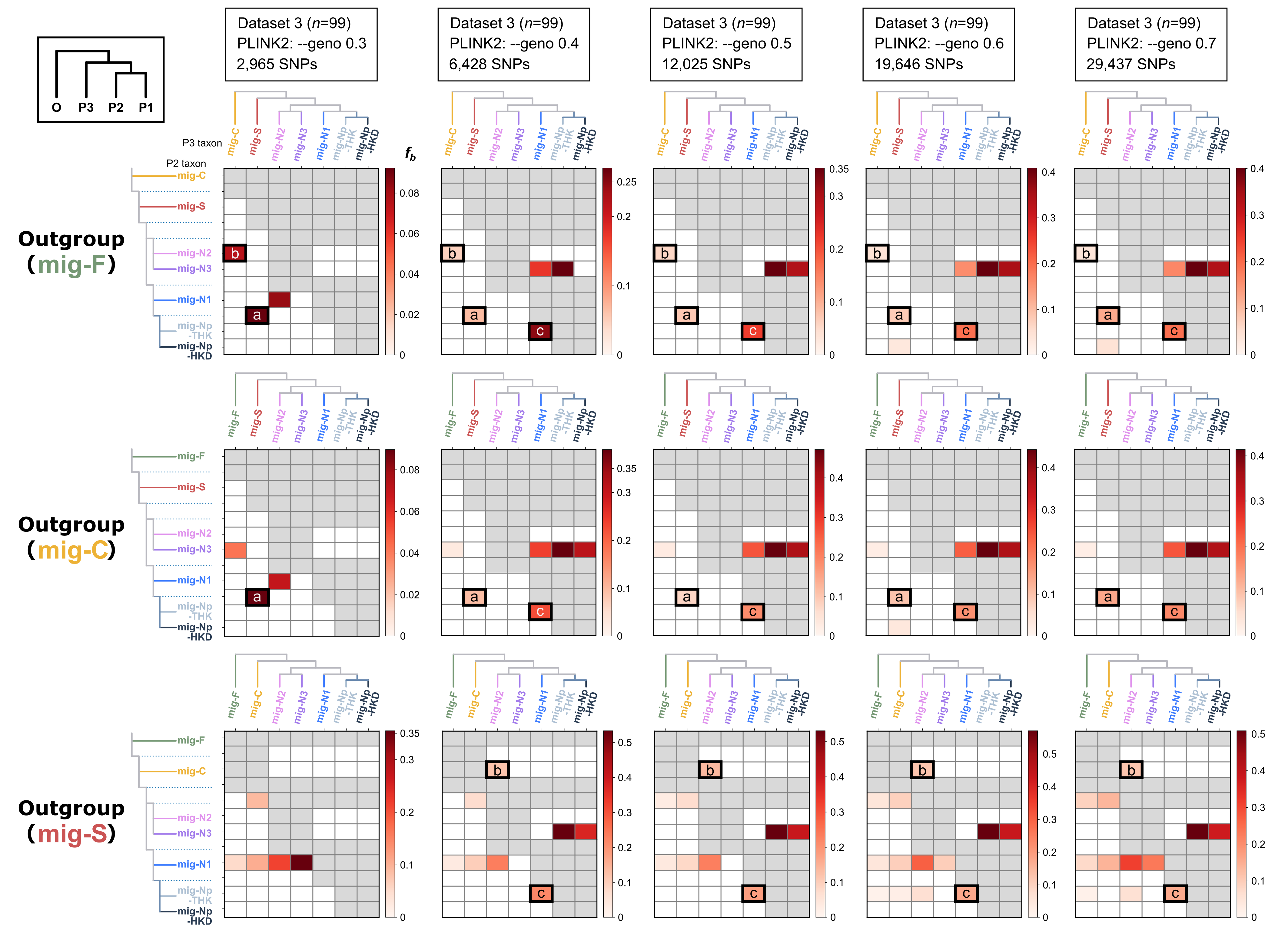
**Figure S15.** *f*-branch statistics for the *Catapionus nebulosus* species group, computed in Dsuite from the unpruned SNP datasets (Dataset 3 in Table S10; minimum depth of 3, --mind 0.97). Rows correspond to the three rooting assumptions and columns to the five --geno thresholds. In each matrix, cells give *f*_b between the lineage in the column (the P3 taxon) and the branch in the row of the tree shown at the left. Color scales differ among matrices. Gray cells mark comparisons that the topology does not define. Three recurring signals are outlined: (a) between mig-S and the common ancestor of mig-Np-THK and mig-Np-HKD, (b) between mig-C and mig-N2, and (c) between mig-N1 and mig-Np-THK. Signal (a) cannot be tested where mig-S is the outgroup. The inset at the top left shows the trio configuration. Trio-level *f*₄-ratios for signals (a) and (c) are shown in **Figures S16 and S17**, and statistics for significant trios are given in **Table S16**.

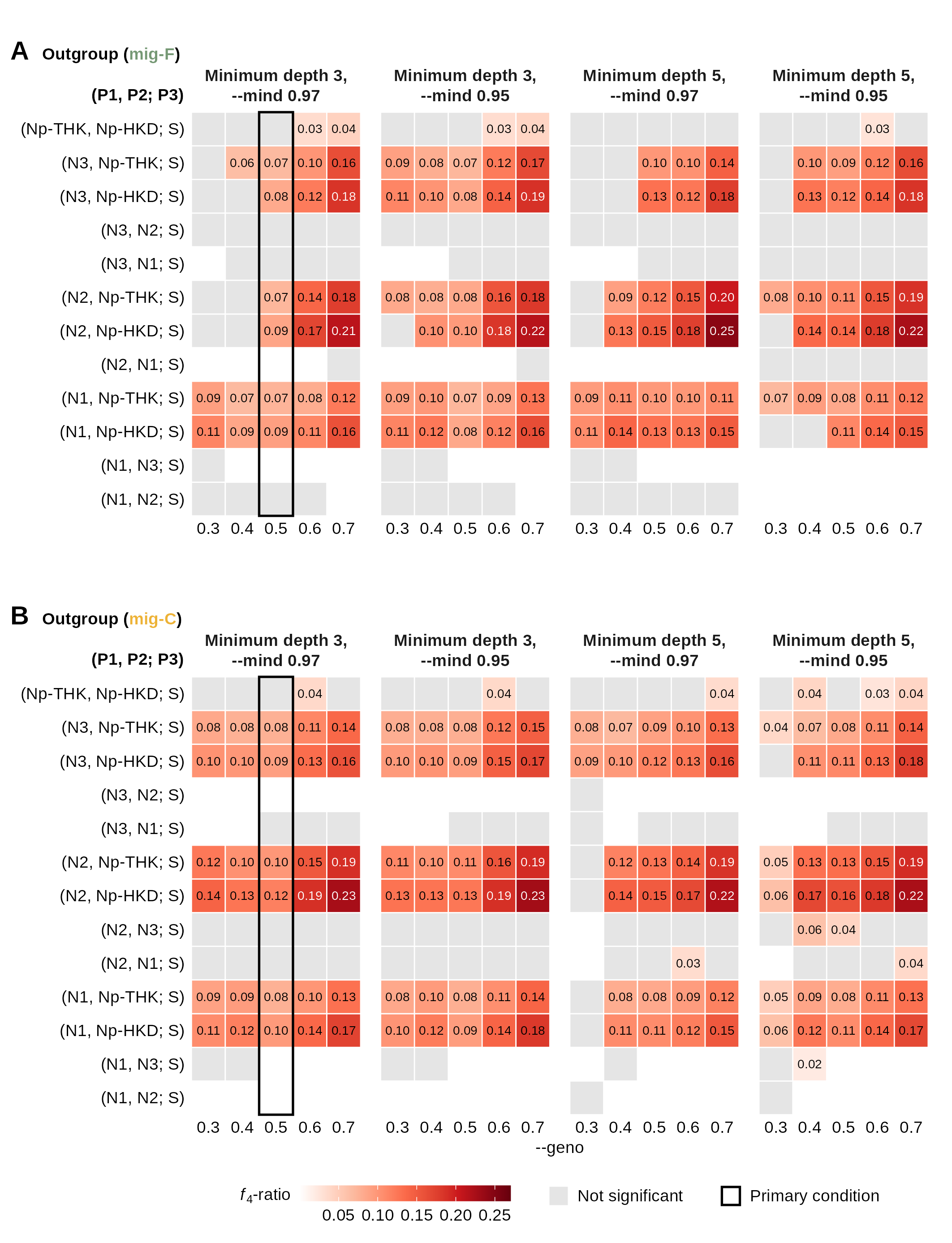
**Figure S16.** *f*₄-ratios of Dsuite trios with mig-S as P3. (A) mig-F as the outgroup. (B) mig-C as the outgroup. Columns show four combinations of minimum depth (3 or 5) and --mind (0.97 or 0.95), each at five --geno thresholds. Rows show trios as (P1, P2; P3), and lineage labels omit the prefix “mig-”. Colored cells show trios that were significant after Bonferroni correction, with the *f*₄-ratio given in each cell. Pale gray cells show trios that were not significant. Dsuite places the lineage that shares more derived alleles with P3 in the P2 position, so each trio appears in only one of the two rows for a given pair. Blank cells indicate that the result is shown in the row with the reverse order. Boxed columns show the primary condition (minimum depth of 3, --mind 0.97, --geno 0.5). These trio-level values underlie signal (a) in **Figure S15**. Statistics for significant trios are given in **Table S16**.

**Figure S17.** *f*₄-ratios of the Dsuite trio (mig-Np-HKD, mig-Np-THK; mig-N1) with mig-F, mig-C, or mig-S as the outgroup. Columns show four combinations of minimum depth (3 or 5) and --mind (0.97 or 0.95), each at five --geno thresholds. Colored cells show trios that were significant after Bonferroni correction, with the *f*₄-ratio given in each cell. Pale gray cells show trios that were not significant. The boxed column shows the primary condition (minimum depth of 3, --mind 0.97, --geno 0.5). These trio-level values underlie signal (c) in **Figure S15**. Statistics for significant trios are given in **Table S16**.
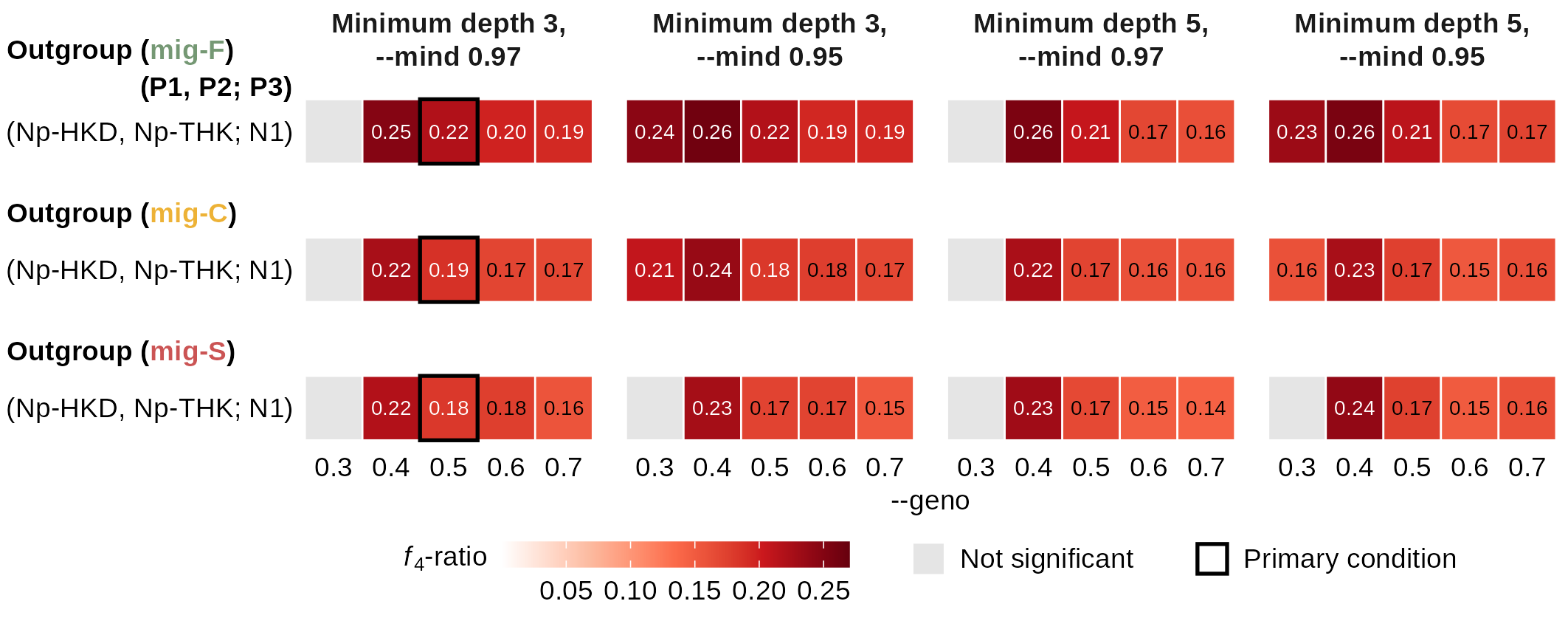

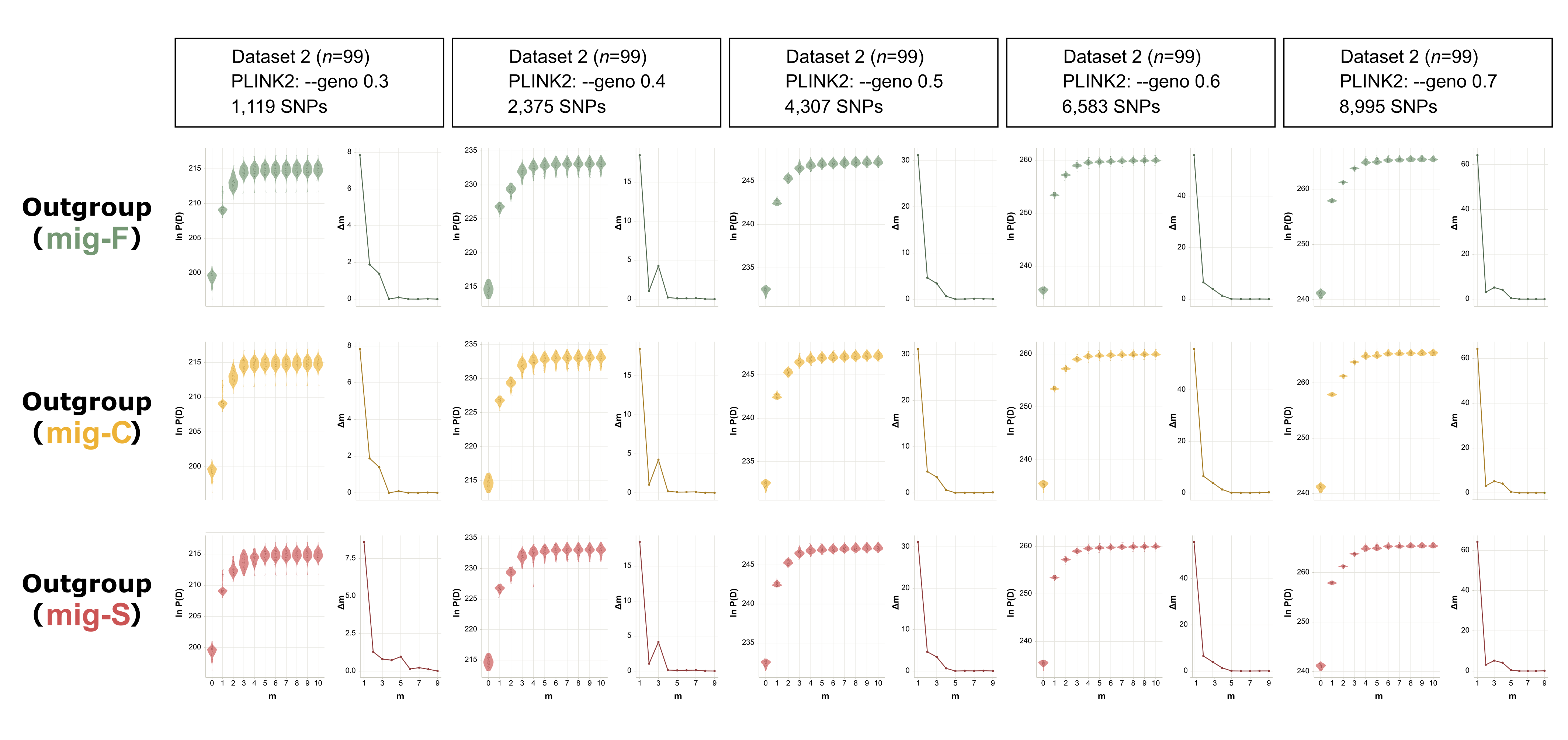
**Figure S18.** Selection of the number of migration edges for the *Catapionus nebulosus* species group, with OptM. Rows correspond to the three candidate rootings and columns to the five --geno thresholds. Within each panel, the left plot gives ln P(D) for m = 0 to 10, with violins showing the distribution across the 50 runs at each m, and the right plot gives the Δm statistic.

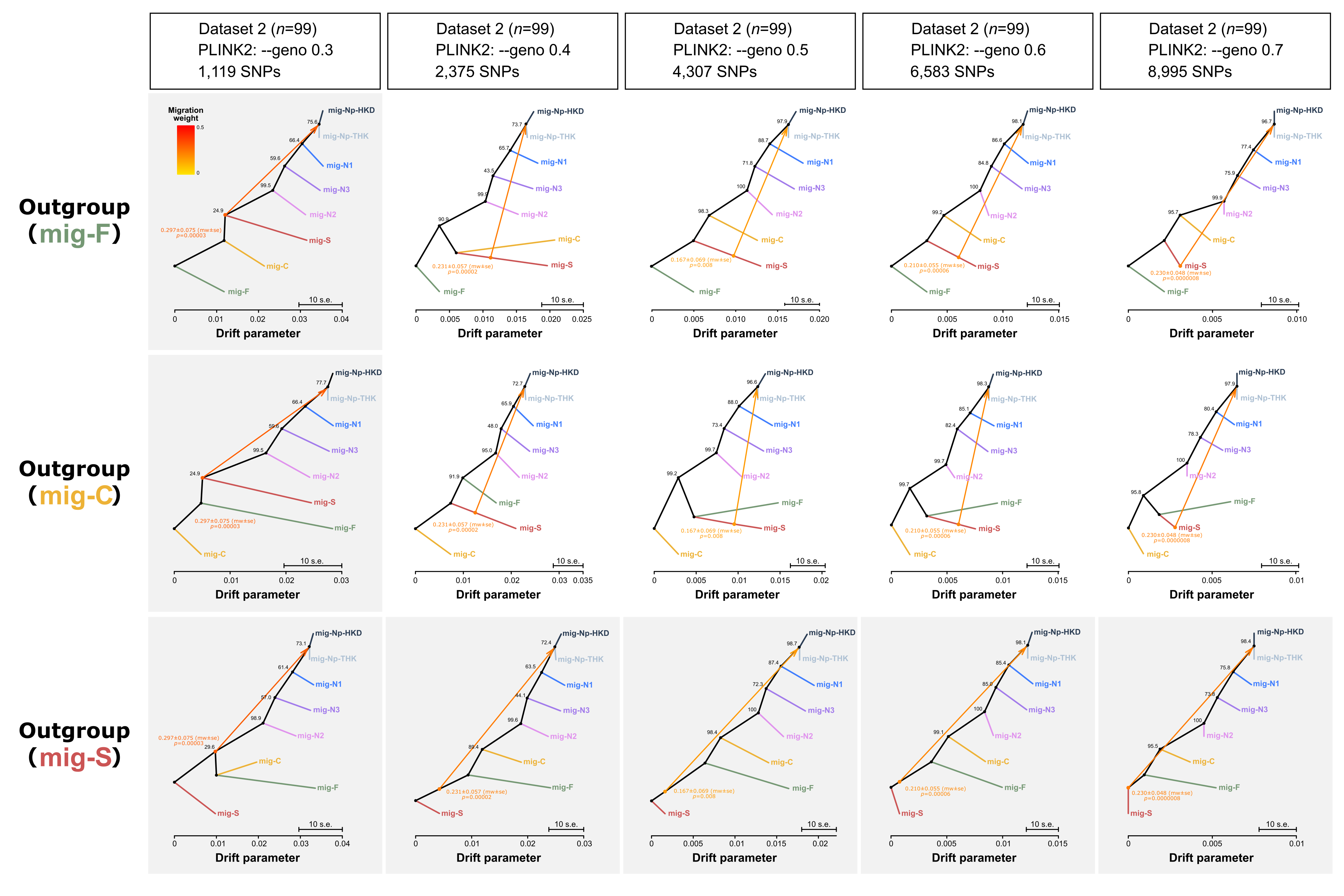
**Figure S19.** Admixture graphs for the *Catapionus nebulosus* species group, inferred with OrientAGraph. Rows correspond to the three candidate rootings and columns to the five --geno thresholds. Panels with a gray background show graphs in which the donor was placed on a branch ancestral to the recipient. Values are given in Table S17.

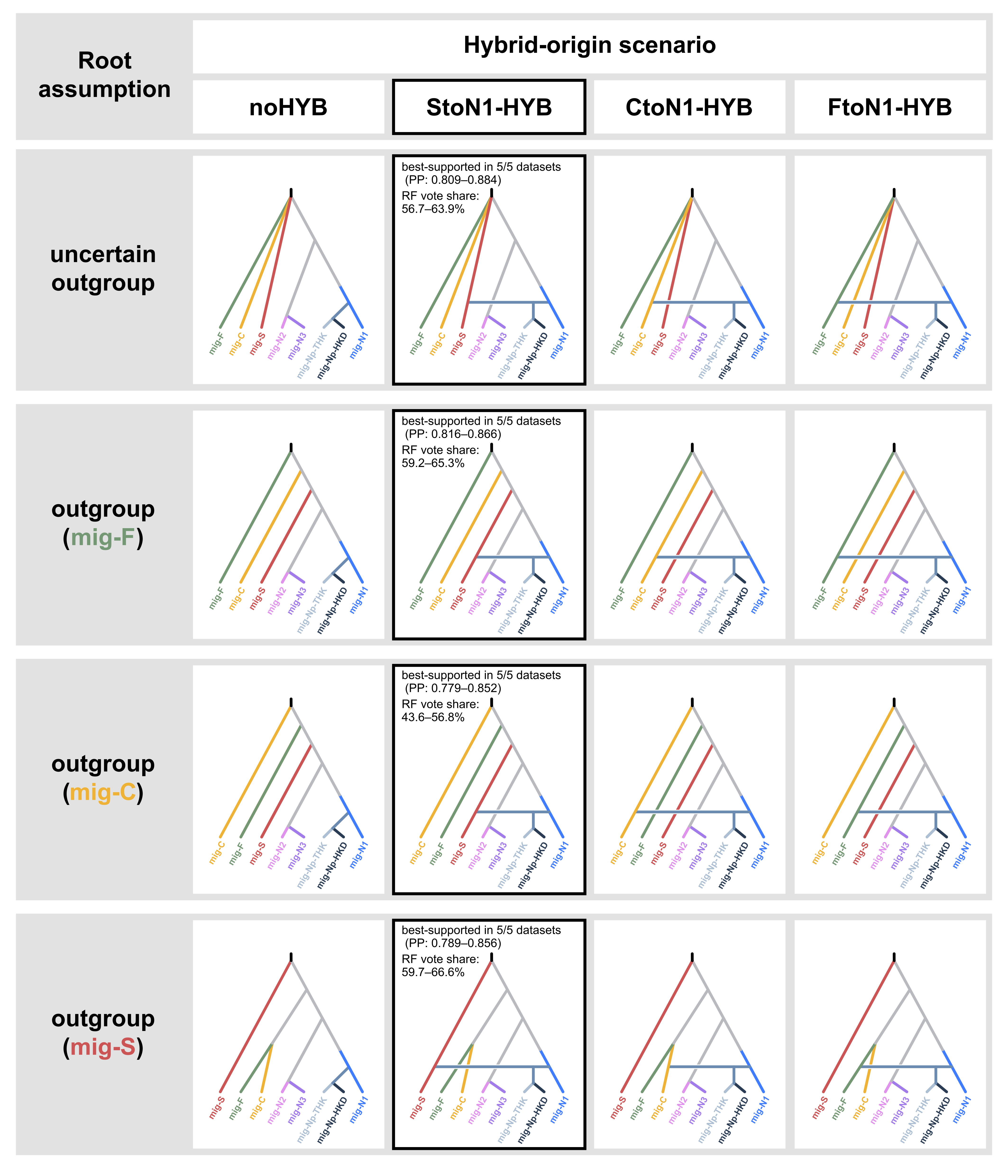
**Figure S20.** Four alternative origin scenarios evaluated using DIYABC-RF under each rooting scenario. Rows correspond to the rooting assumptions (the three candidate rootings and a root-agnostic set), and columns correspond to the origin scenarios. The best-supported scenario, mig-N1 × mig-S, is outlined in each row, with the posterior probability and vote share of the selected model across the five --geno thresholds. Values are given in **Table S18**.

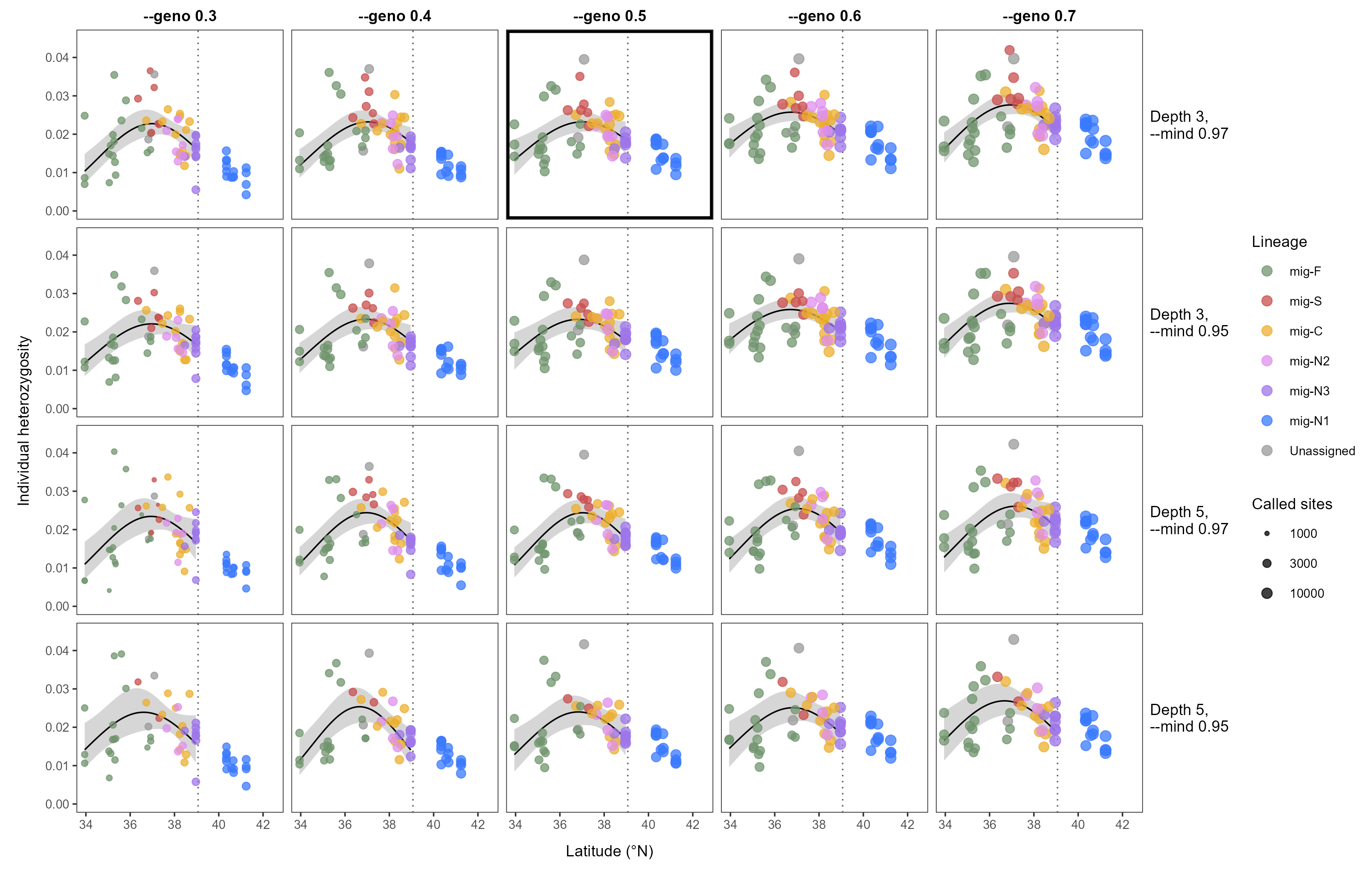
**Figure S21**. Latitudinal trend in individual heterozygosity of the sexual lineages under each filtering condition. Rows give the combinations of minimum depth and individual missingness, and columns give the --geno thresholds. The primary dataset (minimum depth of three, --mind 0.97, and --geno 0.5) is outlined. Each point represents one individual, and its size is proportional to the number of called sites. Colors indicate nuclear lineages, and gray points indicate the mig-X assemblage. Curves show the fit of a beta-binomial GLMM with latitude and its square as fixed effects, genotyping rate as a covariate held at its mean, and locality as a random intercept. Shading shows the 95% confidence interval of the fixed effects. The model was fitted to individuals south of 39°04′N (dotted line), and individuals north of this line are shown but were not included in the fit. Model statistics are given in **Table S20**.
